# A global overview of pleiotropy and genetic architecture in complex traits

**DOI:** 10.1101/500090

**Authors:** Kyoko Watanabe, Sven Stringer, Oleksandr Frei, Maša Umićević Mirkov, Tinca J.C. Polderman, Sophie van der Sluis, Ole A. Andreassen, Benjamin M. Neale, Danielle Posthuma

**Author notes:** Correspondence to: Danielle Posthuma, Department of Complex Trait Genetics, VU University, De Boelelaan 1085, 1081 HV, Amsterdam, The Netherlands.

## Abstract

After a decade of genome-wide association studies (GWASs), fundamental questions in human genetics are still unanswered, such as the extent of pleiotropy across the genome, the nature of trait-associated genetic variants and the disparate genetic architecture across human traits. The current availability of hundreds of GWAS results provide the unique opportunity to gain insight into these questions. In this study, we harmonized and systematically analysed 4,155 publicly available GWASs. For a subset of well-powered GWAS on 558 unique traits, we provide an extensive overview of pleiotropy and genetic architecture. We show that trait associated loci cover more than half of the genome, and 90% of those loci are associated with multiple trait domains. We further show that potential causal genetic variants are enriched in coding and flanking regions, as well as in regulatory elements, and how trait-polygenicity is related to an estimate of the required sample size to detect 90% of causal genetic variants. Our results provide novel insights into how genetic variation contributes to trait variation. All GWAS results can be queried and visualized at the GWAS ATLAS resource (http://atlas.ctglab.nl).

## MAIN TEXT

Since the first genome-wide association study (GWAS) on macular degeneration in 2005^1^, over 3,000 GWASs have been published, for more than 1,000 traits, reporting on over tens of thousands of significantly associated genetic variants^2^. Results from GWASs have increased our insight into the genetic architectures of investigated traits, and for some traits, GWAS results have led to further insight into disease mechanisms^3,4^, such as autophagy for Crohn’s disease^5^, immunodeficiency for Rheumatoid arthritis^6^ and transcriptome regulation through *FOXA2* in the pancreatic islet and liver for Type 2 diabetes^7^. The emerging picture after over a decade of GWASs is that the majority of studied traits are highly polygenic and thus influenced by many genetic variants each of small effect^4,8^, with disparate genetic architectures across traits^9^. Fundamental questions, such as whether all genetic variants or all genes in the human genome are associated with at least one trait, with many or even all traits, and whether the polygenic effects for specific traits are functionally clustered or whether they are randomly spread across the genome, are however still unanswered^4,10,11^. Answers to these questions would greatly enhance our understanding of how genetic variation leads to trait variation and trait correlation. Whereas GWAS primarily aims to discover genetic variants associated with specific traits, the current availability of a vast amount of GWAS results can be used to investigate some of these fundamental questions.

To this end, we compiled a catalogue of 4,155 GWAS results across 2,965 unique traits from 295 studies, including publicly available GWASs and new results for 600 traits from the UK Biobank (http://atlas.ctglab.nl). These GWAS results were used in the current study to achieve the following aims; *i*) charting the extent of pleiotropy at trait-associated locus, gene, SNP and gene-set levels, *ii*) characterizing the nature of trait-associated variants (i.e. the distribution of effect size, minor allele frequency and biological functionality of trait-associated or credible SNPs), and *iii*) understanding the nature of the genetic architecture across a variety of traits and domains in terms of SNP heritability and trait polygenicity (see **Extended Data Fig. 1)**.

### Catalogue of 4,155 GWAS summary statistics for 2,965 unique traits

We collected publicly available full GWAS summary statistics (last update 23^rd^ October 2018; see **Methods**). This resulted in 3,555 GWAS summary statistics from 294 studies. We additionally performed GWAS on 600 traits available from the UK Biobank release 2 cohort (UKB2; release May 2017)^12^, by selecting non-binary traits with >50,000 European individuals with non-missing phenotypes, and binary traits for which the number of available cases and controls were each >10,000 and total sample size was >50,000 (see **Methods**, **Supplementary Information 1** and **Supplementary Table 1-2**). In total, we collected 4,155 GWASs from 295 unique studies and 2,965 unique traits (see **Supplementary Table 3** for a full list of collected GWASs). Traits were manually classified into 27 standard domains based on previous studies^13,14^. The average sample size across curated GWASs was 56,250 subjects. The maximum sample size was 898,130 subjects for a Type 2 Diabetes meta-analysis^15^. The 4,155 GWAS results are made available in an online database (http://atlas.ctglab.nl). The database provides a variety of information per trait, including SNP-based and gene-based Manhattan plots, gene-set analyses^16^, SNP heritability estimates^17^, genetic correlations, cross GWAS comparisons and phenome-wide plots. For the present study, we restricted our analyses to reasonably powered GWASs (i.e. sample size >50,000), to avoid including SNP effect estimates with relatively large standard errors (see **Methods**). By selecting a GWAS with the largest sample size per trait, it resulted in 558 GWASs for 558 unique traits across 24 trait domains. The average sample size of these 558 GWASs was 256,276, and 478 GWASs (85.7%) were based on the UKB2 including 11 meta-analyses with UKB2, 46 (8.2%) on the UK Biobank release 1 cohort (UKB1) including 8 meta-analyses with UKB1, and the remaining were non-UKB cohorts. All results presented hereafter concern these selected 558 GWASs unless specified otherwise. The online database, however, allows researchers to reproduce similar analyses with custom selections of GWASs.

### The extent of pleiotropy

Results of previous GWASs have shown significant associations of thousands of genomic loci with a large number of traits^2,4^. Given a finite number of segregating variants on the human genome, this suggests the presence of widespread pleiotropy. Pleiotropy may be informative to the reasons of co-morbidity between traits, as it may indicate an underlying shared genetic mechanism, and may aid in resolving questions regarding causal effects of one trait on another. However, the exact extent of pleiotropy across the genome is currently unknown^4^. We therefore investigated pleiotropy at locus, gene, SNP and gene-set levels. We defined pleiotropy as the presence of statistically significant associations with more than one trait domain as traits within domain tend to show stronger phenotypic correlations than between domains (see **Supplementary Information 2** and **Extended Data Fig. 2**). Our definition thus refers to ‘statistical pleiotropy’, and includes situations of true pleiotropy (e.g. one SNP directly influences multiple traits), or situations where statistical associations to multiple traits are induced via causational effects of one trait on another, via phenotypic correlations between traits, or via a third common factor^18^. We defined the level of pleiotropy by the number of associated domains, and further grouped into four categories; multi-domain (associated with traits from multiple domains), domain-specific (associated with multiple traits from a single domain), trait-specific (associated with a single trait) and non-associated (**Methods**). We then assessed whether pleiotropic associations at the locus, gene, SNP or gene set level are structurally or functionally different from trait- or domain-specific associations or non-associated sites.

#### Pleiotropic genomic loci

The 558 GWASs yielded 41,511 trait-associated loci (from 470 traits, as 88 traits did not yield any genome-wide significant association after QC; see **Methods**). After grouping physically overlapping trait-associated loci, we obtained 3,362 grouped loci (**Methods**, **Extended Data Fig. 3**, and **Supplementary Table 4**). The total summed length of these loci (1706.0 Mb) covered 61.0% of the genome. Of these, 93.3% were associated with more than one trait and 90.0% were multi-domain loci (**Table 1** and **Extended Data Fig. 4a, b**). The multi-domain and domain-specific loci showed a significantly higher density of protein coding genes compared to non-associated genomic regions (*p*=5.3e-16 and *p*=2.6e-4; **Fig. 1a** and **Supplementary Table 5**).

**Table 1.** Count and proportion of pleiotropic trait-associated loci, genes, SNPs and gene-sets.

|  | Loci |  | Genes |  | SNPs |  | Gene-set |  |
| --- | --- | --- | --- | --- | --- | --- | --- | --- |
|  | Length (Mb) | % | Count | % | Count | % | Count | % |
| **Total in genome** | 2796.10 | 100.00 | 17,444 | 100.00 | 1,740,179 | 100.00 | 10,650 | 100.00 |
| **Associated** | 1706.00 | 61.01 | 11,443 | 65.60 | 236,388 | 13.58 | 1,106 | 10.38 |
| Pleiotropic* | 1592.53 | 93.35 | 9,252 | 80.85 | 142,376 | 60.23 | 606 | 54.79 |
| Multi-domain | 1535.76 | 90.02 | 7,657 | 66.91 | 76,650 | 32.43 | 361 | 32.64 |
| Domain specific | 56.77 | 3.33 | 1,595 | 13.94 | 65,726 | 27.80 | 245 | 22.15 |
| Trait specific | 113.48 | 6.65 | 2,191 | 19.15 | 94,012 | 39.77 | 500 | 45.21 |
| **Non-associated** | 1090.10 | 38.99 | 6,001 | 34.40 | 1,503,791 | 86.42 | 9,544 | 89.61 |

\*The count of pleiotropic loci, genes, SNPs and gene-sets is the sum of the multi-domain and domain specific categories. Proportion of pleiotropic, multi-domain, domain specific and trait specific categories are relative to the associated loci, SNPs, genes or gene-sets, respectively.

**Fig. 1.**
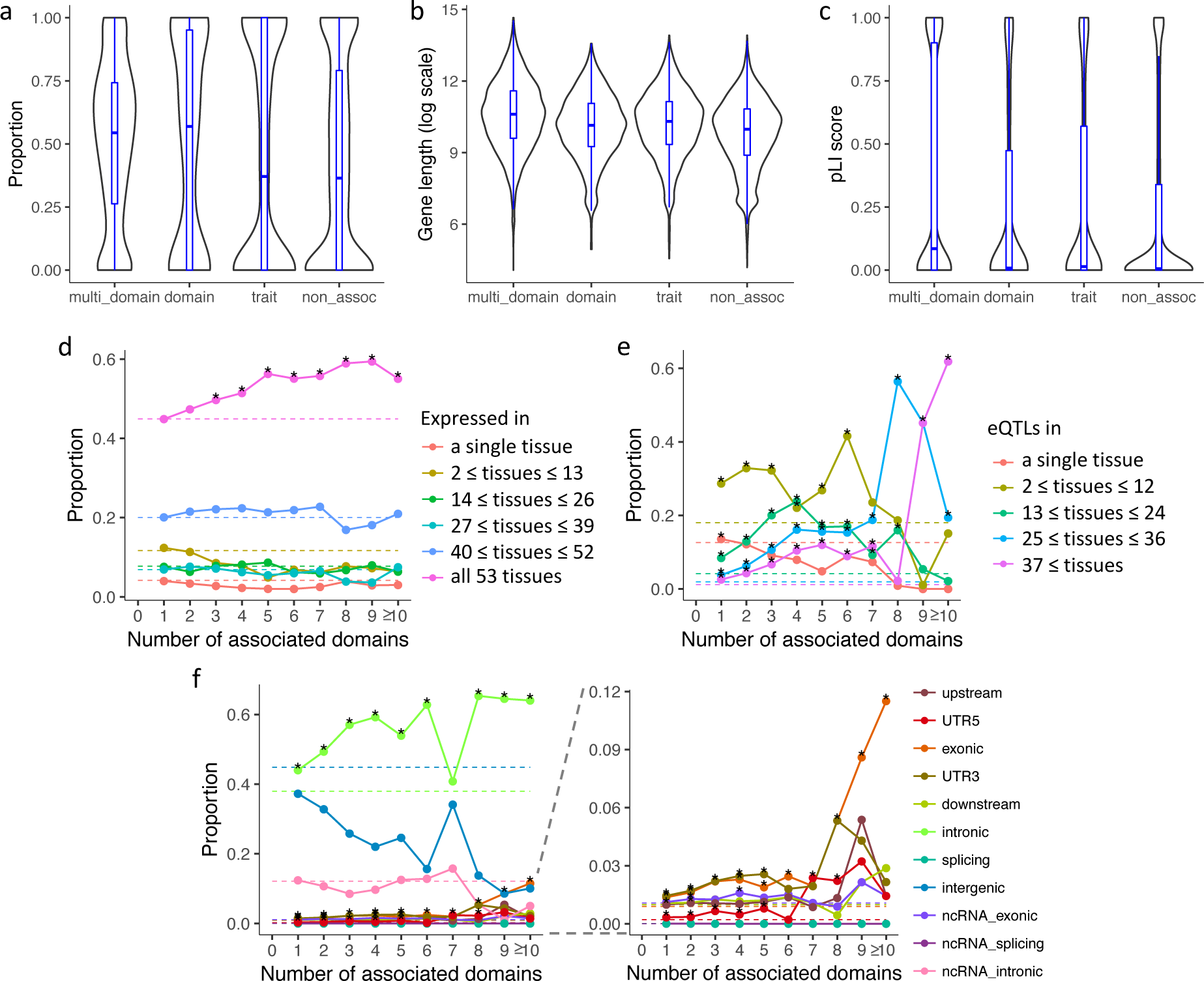
Trait-associated locus, gene and SNP pleiotropy across the genome. **a.** Distribution of gene density of loci with different association types. **b.** Distribution of gene length in log scale with different association types. **c.** Distribution of pLI score of genes with different association types. For **a-c**, multi_domain: associated with traits from >1 domain, domain: associated with >1 traits from a single domain, trait: associated with a single trait, non_assoc: not associated with any of 558 traits. **d.** Tissue specificity of genes at different levels of pleiotropy. Each data point represents a proportion of genes expressed in a given number of tissues for a specific number of associated domains. **e.** Proportion of SNPs with different functional consequences at different levels of pleiotropy. Each data point represents the proportion of SNPs with a given functional consequence for a specific number of associated domains. **f.** Tissue specificity of SNPs based on active eQTLs at different levels of pleiotropy. Each data point represents the proportion of SNPs being eQTLs in a given number of tissues for a specific number of associated domains. For **d-f**, dashed lines refer to the baseline proportions (relative to all 17,444 genes (d) or all 1,740,179 SNPs (e-f)), and stars denote significant enrichment relative to the baseline (Fisher’s exact test, one-sided).

The locus associated with the largest number of traits and domains (i.e. the most pleiotropic locus) was the MHC region (chr 6:25Mb-37Mb), which contained 441 trait-associated loci from 213 traits across 23 trait domains. The MHC region is well-known for its complex structure of linkage disequilibrium, spanning over 300 genes. The extremely pleiotropic nature of this region might, therefore, be explained by its long-ranged LD block due to overlap of multiple independent signals from multiple traits. Similarly, high locus pleiotropy, not limited to the MHC region, can occur purely due to the overlap of the LD blocks of the loci in the grouped locus, and they may not share the same causal SNPs. By performing colocalization (i.e. statistically identifying loci sharing the same causal SNP) for all possible pairs of physically overlapping trait-associated loci (see **Methods**, **Supplementary Information 3** and **Extended Data Fig. 3**), we indeed observed a decrease in the number of associated traits and trait domains per group of colocalized loci compared to loci defined by physical overlap (**Extended Data Fig. 4** and **Supplementary Table 6**). In addition, loci grouped based on physical overlap often contained multiple independent groups of colocalized loci (**Supplementary Table 6**). Therefore, physical overlap of trait-associated loci does not necessary mean that the same causal SNPs are involved in the traits associated with such a grouped locus. Examination of pleiotropy at the gene or SNP level will provide further insight into the nature of the pleiotropy observed at the locus level.

#### Pleiotropic genes

We next investigated the extent of pleiotropy at the gene level. For this, we conducted a gene-based analysis on 17,444 protein-coding genes using MAGMA for each trait^16^ (**Methods**). Of the 558 traits, 516 yielded at least one significantly associated gene and 11,443 (65.6%) genes were significantly associated to at least one trait (**Supplementary Table 7**). Of these, 81.0% were associated with more than one trait and 66.9% were associated with traits from multiple domains (**Table 1** and **Extended Data Fig. 5a, b**). We found that genes associated with at least one trait are significantly longer than genes that are not associated with any of the 558 tested traits (*p*=2.1e-194, *p*=8.7e-12 and *p*=3.8e-29 for multi-domain, domain-specific and trait-specific genes, respectively; **Fig. 1b** and **Supplementary Table 8**). As the MAGMA algorithm is insensitive to bias caused by gene-length, these findings are unlikely to be due to larger genes having an increased statistical probability to be significantly associated (**Supplementary Information 4, Extended Data Fig. 5c** and **Supplementary Table 9**). The multi-domain genes showed a significantly higher probability of being intolerant to loss of function mutations (pLI score)^19^ compared to trait-/domain-specific and non-associated genes (*p*=1.2e-79, *p*=4.8e-22 and *p*=2.8e-19, respectively; **Fig. 1c** and **Supplementary Table 10**), suggesting that more pleiotropic genes are on average less tolerant to loss of function variants. The most pleiotropic genes are located in the MHC region, yet a region on chromosome 3 also spanned multiple genes with high levels of pleiotropy (**Extended Data Fig. 5a**). In this region, *BSN* was associated with the largest number of trait domains (94 traits across 17 domains).

We next tested whether tissue specificity of genes was related to the level of pleiotropy by counting the number of active tissues per gene based on gene expression profiles for 53 tissue types obtained from GTEx^20^ (see **Methods**). The results showed that the proportion of genes expressed in all 53 tissue types increases along with the level of pleiotropy (*p*=9.7e-05, **Fig. 1d** and **Supplementary Table 11**). This indicates that more pleiotropic genes tend to be active in multiple tissue types, suggesting that those genes are involved in general biological functions across the human body.

#### Pleiotropic SNPs

The level of pleiotropy at a locus or gene level does not necessarily translate to pleiotropy at the level of the SNP. For example, within the same locus or gene, multiple SNPs may be significantly associated with different traits. A locus or gene can thus show a higher level of pleiotropy compared to individual SNPs. We, therefore, investigated the extent of pleiotropy at the level of the SNP. To do so, we extracted 1,740,179 SNPs that were present in all 558 GWAS results. We first confirmed that this selection of SNPs had the same distribution of their location across the genome and their functional consequences as all known SNPs on the genome (**Methods** and **Extended Data Fig. 6a, b**). We note that some of the observed SNP-pleiotropy may still be induced by LD, e.g. a SNP could reach genome-wide significance because of its strong LD with a causal SNP. However, the purpose of this analysis is to identify individual SNPs (not loci) that are associated with multiple trait domains and their functions. Of these, 237,120 (13.6%) were genome-wide significant (*p*<5e-8) in at least one of the 558 traits (**Extended Data Fig. 6c** and **Supplementary Table 12**). Out of 237,120 SNPs that were associated with at least one trait, 60.2% were associated with more than one trait and 32.4% were associated with more than one domain (**Table 1** and **Extended Data Fig. 6d**).

These pleiotropic SNPs spread broadly across the genome but were not evenly distributed, i.e. chromosome 1, 11, 12, 15, 17, 20 and 22 showed relative enrichment of pleiotropic SNPs (**Supplementary Information 5** and **Supplementary Table 13**). Of all associated SNPs, the most pleiotropic SNP, located in the MHC region (rs707939; an intronic SNP of *MSH5*) was associated with 48 traits from 13 domains. There were 45 SNPs associated with 12 trait domains, of which 35 were located on chromosome 3, 49.8Mb-50.1Mb overlapping with 5 protein coding genes, *TRAIP*, *CAMKV*, *MST1R*, *MON1A* and *RBM6*. These SNPs include two exonic SNPs, rs2681781 (synonymous on *CAMKV*) and rs2230590 (nonsynonymous on *MST1R*; **Supplementary Table 12**).

To investigate whether SNPs with a higher level of pleiotropy have different functional annotations than less pleiotropic SNPs, we investigated how functional consequence and tissue specificity in terms of expression quantitative trait loci (eQTLs) were represented across different levels of SNP pleiotropy (**Methods**). We found that the proportion of intronic and exonic SNPs increased as a function of the level of pleiotropy (*p*=2.2e-3 and *p*=1.7e-2, respectively); the proportion of exonic SNPs increased from less than 1% to over 5%, and the proportion of intronic SNPs increased from less than 40% to over 50% (**Fig. 1e** and **Supplementary Table 14**) with increasing levels of pleiotropy. The proportion of SNPs within flanking regions such as 5’ and 3’ untranslated regions (UTR) also increased with the number of associated domains. At the same time, we observed a steep decrease of the proportion of intergenic SNPs with increasing level of SNP pleiotropy (*p*=8.1e-4; **Fig. 1e** and **Supplementary Table 14**). Based on active eQTLs, the proportion of SNPs being eQTLs in a greater number of tissue types (>24 tissue types out of 48) increased along with the number of associated domains (*p*=8.4e-3 and *p*=1.1e-2 for eQTLs in between 25 and 36 tissues, and between 37 and 48 tissues, respectively) while SNPs in genes expressed in a single or less than half of available tissue types showed decreasing proportion (**Fig. 1f** and **Supplementary Table 15**). These results suggest that highly pleiotropic SNPs are more likely to be genic (exonic and intronic) and less likely to be tissue specific.

#### Pleiotropic gene-sets

Pleiotropy at the level of trait-associated loci, genes or SNPs do not necessarily suggest the presence of shared biological pathways across multiple traits. To assess the level of pleiotropy at the level of gene-sets, reflecting a biological meaningful grouping of genes, we performed MAGMA gene-set analyses for 558 traits using 10,650 gene-sets (Methods). In total, 235 (42.1%) traits showed significant association with one of 1,106 (10.4%) gene-sets. The most pleiotropic gene-set was ‘Regulation of transcription from RNA polymerase II promoter’ (GO biological process) associated with 61 traits from 9 domains, followed by 7 other gene-sets associated with 7 domains, of which 5 of them were also involved in regulation of transcription (**Supplementary Table 16**). We observed that the number of genes in a gene-set was significantly larger for highly pleiotropic gene-sets (associated with more than one domain) compared to other gene-sets (domain-specific, trait-specific and non-associated; *p*=4.1e-12, *p*=1.6e-13 and *p*=1.2e-29, respectively; **Extended Data Fig. 7a**, and **Supplementary Table 17**). Since GO terms (55.6% of tested gene-sets) have a hierarchical structure, the larger gene-sets are more likely to be located at the top of the hierarchy, representing more general functional categories.

In contrast to the pleiotropy at gene level where 80.9% genes were associated with more than one trait, we only found 54.8% of the associated gene-sets to be pleiotropic (**Table 1**). We observed that the proportion of pleiotropic genes per gene-set is not uniformly distributed, and pleiotropic genes tend to cluster into a subset of gene-sets, explaining the decreased proportion of pleiotropic gene-sets compared to pleiotropic genes (**Extended Data Fig. 7b, c**). At the same time, the higher proportion of trait-specific gene-sets (45.2%) compared to trait-specific genes (19.2%) suggests that, given current definitions of gene-sets, the combination of associated genes is rather unique to a trait and focusing on gene-sets to gain insight into trait-specific biological mechanisms may be more informative than focusing on single genes (**Supplementary Information 6**).

#### Genetic correlations across traits

Above we showed that of all trait-associated loci, genes and SNPs that are associated with at least one trait, 90.0%, 66.9% and 32.6% are associated with more than one domain, respectively. Such wide-spread pleiotropy indices non-zero genetic correlations between traits. To test whether genetic correlations are evenly present across traits or cluster into trait domains, we computed pairwise genetic correlations (*r_g_*) across 558 traits using LDSC^17^.

We calculated the proportion of trait pairs with an *r_g_* that is significantly different from zero across all 558 traits, within domains and between domains. Out of 155,403 possible pairs across 558 traits, 24,106 pairs (15.5%) showed significant genetic correlations after Bonferroni correction (*p*<0.05/155,403=3.2e-7) with an average |*r_g_*| of 0.38.

In principle, if the trait domains contain traits that are biologically related, we would expect that traits within the same domain have stronger genetic correlations than traits across domains. The proportion of pairs with a significant genetic correlation within a domain was especially high in cognitive, ‘ear, nose, throat’, metabolic and respiratory domains, and for most of domains, average |*r_g_*| across significant trait pairs was higher than 0.38 (across all traits). Note that the proportion of trait pairs with significant *r_g_* may be biased by sample size and *h*^*2*^_*SNP*_ of traits within a domain; across 558 traits, the worst case scenarios with the minimum observed *h*^*2*^_*SNP*_ (0.0045 with sample size 385,289) or the minimum sample size (51,750 with *h*^*2*^_*SNP*_ =0.0704) required *r_g_* to be above 0.39 or 0.18, respectively, to gain a power of 0.8 (**Methods**). Within domain, the majority of significant genetic correlations was positive and the average |*r_g_*| was above 0.5 in most of the domains (**Fig. 2a** and **Supplementary Table 18**). Between domains, the proportion of pairs with significant genetic correlations was generally lower than within domains, and most of the domain pairs showed average |*r_g_*|<0.4 (**Fig. 2b** and **Supplementary Table 19**). Some trait domains showed a predominance of negative genetic correlations with other domains, i.e. activity, cognitive, reproduction and social interaction domains. We further clustered traits based on genetic correlations, which resulted in the majority of clusters contained traits from multiple domains (**Methods**, **Supplementary Information 7** and **Extended Data Fig. 8**). These results suggest that although |*r_g_*| is higher within domain than across domains, the trait domains do not necessary reflect genetic similarity across traits.

**Fig. 2.**
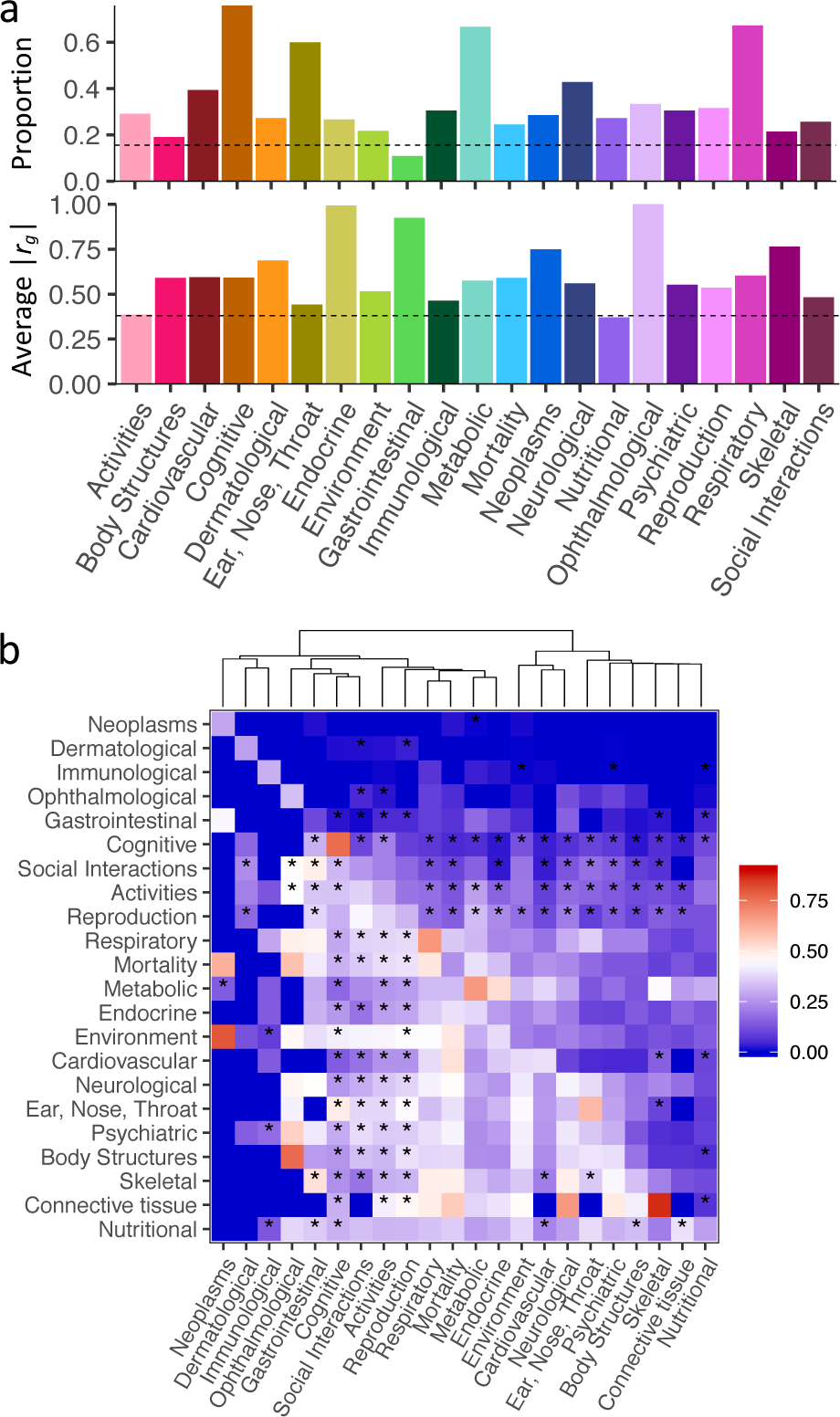
Within and between domains genetic correlations. **a.** Proportion of trait pairs with significant *r_g_* (top) and average |*r_g_*| for significant trait pairs (bottom) within domains. Dashed lines represent the proportion of trait pairs with significant *r_g_* (top) and average |*r_g_*| for significant trait pairs (bottom) across all 558 traits, respectively. Connective tissue, muscular and infection domains are excluded as these each contains less than 3 traits. **b.** Heatmap of proportion of trait pairs with significant *r_g_* (upper right triangle) and average |*r_g_*| for significant trait pairs (lower left triangle) between domains. Connective tissue, muscular and infection domains are excluded as each contains less than 3 traits. The diagonal represents the proportion of trait pairs with significant *r_g_* within domains. Stars denote the pairs of domains in which the majority (>50%) of significant *r_g_* are negative.

### The nature of trait-associated variants

We now address the question whether trait-associated variants differ from genetic variants that are not associated with any trait. For this purpose, we extracted all lead SNPs from each of the 558 GWASs. Lead SNPs were defined per trait at the standard threshold for genome-wide significance (*p*<5e-8) and using an *r*^*2*^ of 0.1 to obtain near-independent lead SNPs, based on the population-relevant reference panel (see **Methods**). Lead SNPs with minor allele count (MAC) ≤100 (based on MAF and sample size of the SNP) were excluded due to lower statistical power and a high false positive rate of effects of SNPs with extremely small MAF. This resulted in 82,590 lead SNPs for 476 traits, reflecting 43,455 unique SNPs. Out of 558 traits, 82 traits did not yield any genome-wide significant lead SNP after QC.

#### Distribution of MAF and effect sizes of lead SNPs

12.3% of the 43,455 (unique) lead SNPs derived from the 558 GWASs had a MAF below 0.01 which is significantly less than expected given the proportion of rare variants in the reference panels (*p*<1e-323; **Supplementary Information 8**), while the distribution of lead SNPs with a MAF above 0.01 was nearly uniform (**Fig. 3a**).

**Fig. 3.**
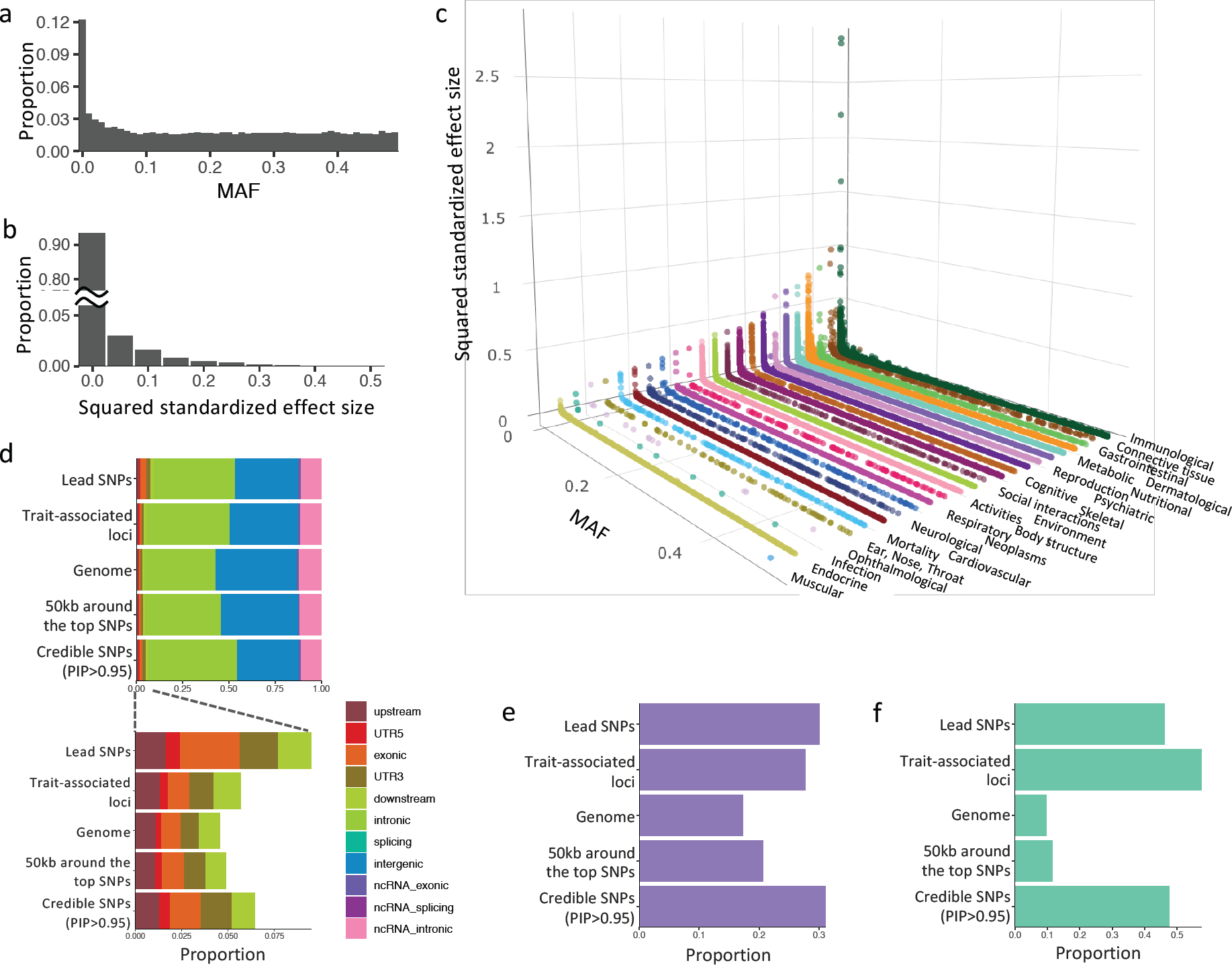
Distribution and characterization of lead SNPs and credible SNPs of 558 traits. **a.** Histogram of MAF of the unique lead SNPs. **b.** Histogram of squared standardized effect size of lead SNPs. **c.** Scatter plot of MAF and squared standardized effect sizes of lead SNPs grouped by trait domains. **d.** Distribution of functional consequences of SNPs. **e.** Proportion of SNPs that overlap with active consequence chromatin state (≤7) across 127 tissue/cell types. **f.** Proportion of SNPs overlapping with significant eQTLs from any of 48 available tissue types.

To gain insight into the distribution of effect sizes across lead SNPs, we calculated the standardized effect size (*β*) from Z-statistics as a function of MAF and sample size^21^, and inspected the distribution of the squared standardized effect sizes (*β^2^*) for lead SNPs across all traits (**Methods**). *β* ranged between 0.01 and 1.70, and *β^2^* is proportional to the variance explained. The median *β^2^* of the lead SNPs across all traits was 5.7e-4 (4.9e-4 and 6.0e-2 for lead SNPs with MAF≥0.01 and <0.01, respectively), and 94.6% of lead SNPs had a *β^2^* below 0.05 (**Fig. 3b**). Thus, the vast majority of lead SNPs thus explained less than 0.05% of the trait variance. We observed a relationship between MAF and standardized effect size, with rare variants (MAF<0.01) showing larger effect sizes (**Fig. 3c**). This is in line with the notion that rare variants are more likely to have large effects compared to common variants, as they are less likely to be under strong selective pressure^22^. However, we also note that statistical power for detecting the rare variants is un-stable^23^. Given that the proportion of rare lead SNPs is larger than the proportions in other MAF bins, it is possible that the distribution of the effect sizes has longer tails for SNPs with MAF<0.01. For most of the traits, a similar relationship between MAF and standardized effect size was observed (**Extended Data Fig. 9**), but large variation across traits was seen in terms of the number of rare lead SNPs, with e.g. a large proportion of rare variants influencing nutritional and connective tissue domains (see **Supplementary Information 8, Extended Data Fig. 10** and **Supplementary Table 20-21**).

#### Characterization of trait-associated loci and lead SNPs

Here we sought to characterise differences in the distribution of functional annotations when comparing SNPs within trait-associated loci to all SNPs in the genome, and comparing lead SNPs to SNPs in the trait-association loci (**Methods**). We first compared SNPs in the trait-associated loci against the entire genome. The strongest enrichment of SNPs in trait associated loci was seen in flanking regions (upstream, downstream, 5’ and 3’ UTR) with average fold enrichment (*E*) 1.31 (**Fig. 3d** and **Table 2**). Non-coding SNPs, in total, covered 93.1% of SNPs in the trait-associated loci, while intergenic SNPs were significantly depleted (*E*=0.83) and intronic SNPs significantly enriched compared to all SNPs in the genome (*E*=1.17; **Table 2**). SNPs in trait-associated loci were also slightly enriched for being exonic compared to the entire genome (*E*=1.07). Active chromatin states and eQTLs were also significantly enriched with notably high enrichment of eQTLs (*E*=1.61 and 5.95, respectively; **Table 2**).

**Table 2.**
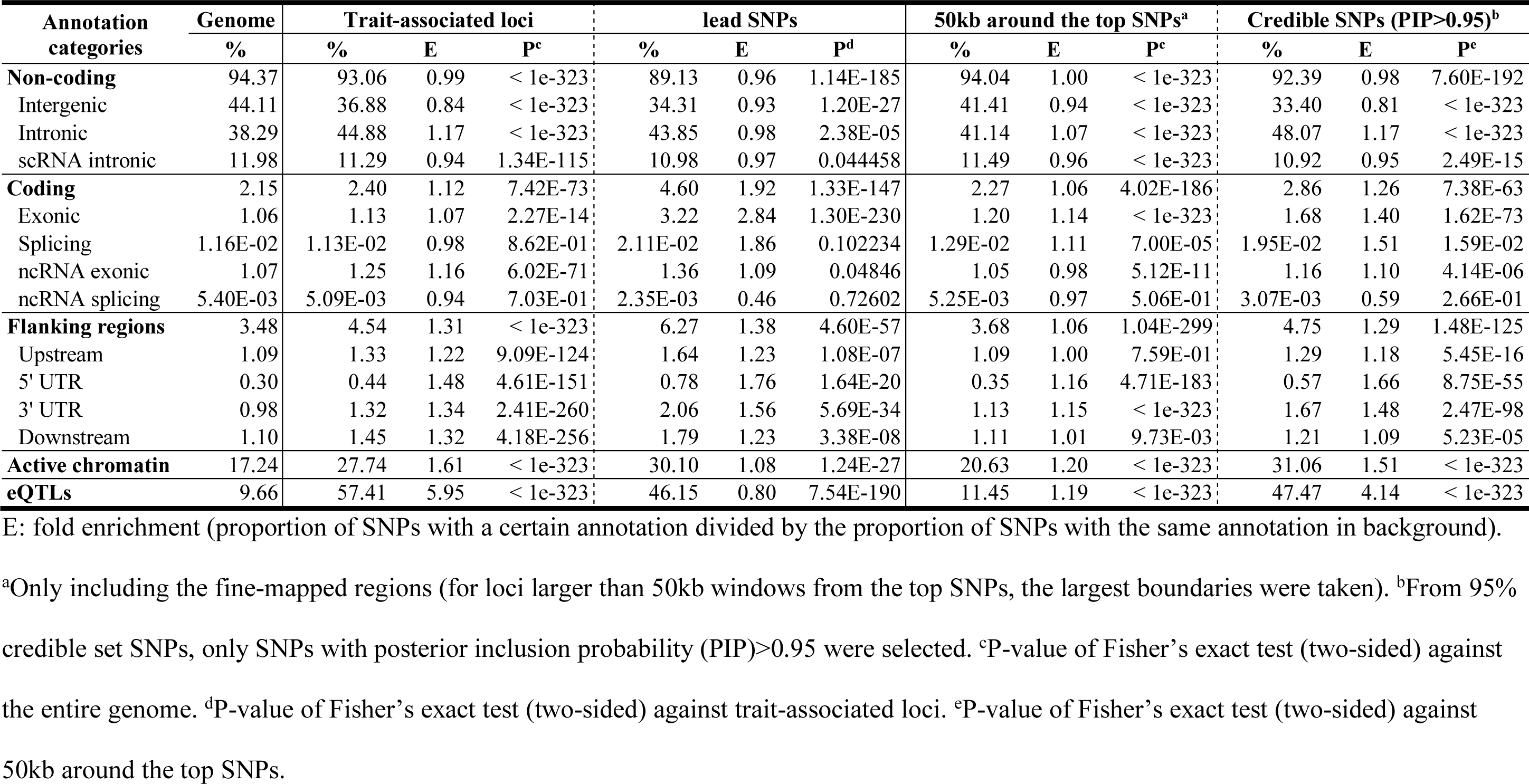
Characteristics of lead SNPs and credible SNPs with PIP>0.95 across 558 traits versus all SNPs in the genome.

We next compared lead SNPs with SNPs in the trait-associated loci. The strongest enrichment for lead SNPs was seen in exonic SNPs (*E*=2.84) followed by flanking regions (*E*=1.38), while intronic and intergenic regions were slightly depleted (average *E*=0.95; **Fig. 3d** and **Table 2**). These results clearly indicate that SNPs located in exonic and flanking regions tend to show stronger effect sizes than other SNPs in the trait-associated loci. On the other hand, active chromatin states showed slight enrichment (*E*=1.08) while eQTLs were significantly depleted (*E*=0.80; **Fig. 3e-f** and **Table 2**). This suggests that SNPs within the trait-associated loci largely overlap with regulatory elements but these elements do not always have the strongest effect sizes within the loci.

#### Characterization of credible set SNPs based on fine-mapping

Owing to the small effect sizes of variants in complex traits and extensive LD throughout the human genome, there is a reasonable chance that lead SNPs (i.e. defined based on LD and P-values) are not the causal SNPs in the trait-associated loci^24^, even when the causal SNPs are actually measured or imputed. Statistical fine-mapping utilizes evidence of the associations at each variant in the loci (effect sizes and LD structure) to assign posterior probability of each specific model at particular locus, which are then used to infer the posterior probabilities of each SNP being included in the model (posterior inclusion probability, PIP) and ascertain the minimum set of SNPs required to capture the likely causal variant. We performed fine-mapping using FINEMAP software^25^ for each trait-associated locus, setting the maximum number of SNPs in the causal configuration (*k*) to 10 and using randomly selected 100k individuals from UKB2 as a reference panel (see **Methods**). From all of the loci associated with at least one of the 558 traits, we obtained a list of credible SNPs with PIP>0.95 consists of 196,542 SNPs (**Supplementary Information 9**).

Next we characterized credible SNPs in respect to their functional annotations, similar as done above with lead SNPs. We thus compared SNPs in the fine-mapped regions to all SNPs in the genome, and credible SNPs to SNPs in the fine-mapped regions. The enrichment pattern of SNPs in the fine-mapped regions was similar to SNPs in the trait-associated loci; i) e. significant enrichment of SNPs in intronic and flanking regions but the fold enrichment was much smaller (**Fig. 3d** and **Table 2**). This is mainly because the fine-mapped regions are often larger than the trait-associated loci by taking 50kb around the top SNPs of the trait-associated loci. In contrast, fold enrichment of exonic SNPs was slightly higher than trait-associated loci (**Table 2**). As we observed higher gene-density around the trait-associated loci, expanding the loci resulted in larger proportion of exonic regions. Both active chromatin state and eQTLs were significantly enriched, however, fold enrichment of eQTLs was notably less than trait-associated loci (**Fig. 3e-f** and **Table 2**). Similar to the lead SNPs, credible SNPs showed strong enrichment in exonic (*E*=1.40) and flanking regions (*E*=1.29), as well as intronic regions (*E*=1.17; **Table 2**). Although an enrichment of active chromatin state is consistent with the result observed in the lead SNPs (*E*=1.51), eQTLs were also significantly enriched in credible SNPs with very strong fold increase (*E*=4.14; **Fig. 3e-f** and **Table 2**).

In summary, the number of credible SNPs is 4.5 times larger than the number of lead SNPs, since for determining lead SNPs, all SNPs that have high LD with lead SNPs are discarded while the fine-mapping captures likely causal SNPs given the observed pattern of association and LD structure. Lead SNPs and credible SNPs show different distributions of enrichment in tested biological functions. We observed a decreased proportion of exonic SNPs and an increased proportion of non-coding or regulatory SNPs within the credible SNPs compared to the lead SNPs. These findings may be due to the fact that coding SNPs tend to have higher effect sizes and are more often assigned as lead SNPs, while the fine-mapping in regions containing some of these causal coding variants may disperse a proportion of probability to adjacent variants. On the other hand, in loci where causal variants are acting through regulatory mechanisms, the credible sets may be more likely to capture the actual, single or multiple causal variants as compared to the lead SNPs.

### The nature of genetic architecture

The genetic architecture of a trait reflects the characteristics of genetic variants that contribute to the phenotypic variability, and is defined by e.g. the number of variants affecting the trait, the distribution of effect sizes, the MAF and the level of interactions between SNPs ^9^. To gain insight into how the genetic architecture varies across multiple complex traits, we assessed the SNP heritability (*h*^*2*^_*SNP*_) and the polygenicity of 558 traits.

#### SNP heritability

*h*^*2*^_*SNP*_ is an indication of the total amount of variance that is captured by the additive effects of all variants included in a GWAS. *h*^*2*^_*SNP*_ depends on several factors, such as the number of SNPs included in the analyses based on their MAF given the current sample size, the polygenicity of the trait (i.e. how many SNPs have an effect) and the distribution of effect sizes. We estimated *h*^*2*^_*SNP*_ for each trait using LDSC^17^ and SumHer from LDAK^26,27^ (**Methods**). The estimates of *h*^*2*^_*SNP*_ using LDSC and SumHer showed strong positive correlation (*r*=0.77 and *p*=3.8e-111; **Fig. 4a**). Therefore, we focus on estimates based on LDSC, hereafter, however complete results are available in **Supplementary Table 22** and discussed in **Supplementary Information 10** (**Extended Data Fig. 11**). The highest *h*^*2*^_*SNP*_ was observed for height (*h*^*2*^_*SNP*_=0.31) followed by bone mineral density (*h*^*2*^_*SNP*_=0.27). Of 558 traits, 214 traits, with an average sample size 292,267, showed *h*^*2*^_*SNP*_ less than 0.05. Most of these traits are classically regarded as ‘environmental’ (e.g. current employment status, illness of family members and transport types or activity traits including frequency and type of physical activities and type of accommodation), and tend to have a low *H*^214^. For these traits, the number of detected trait-associated loci is also very low with a median 3. Given the combination of current sample size of > 200,000 and low *h*^*2*^_*SNP*_, this suggests that for these traits increasing the sample size may not lead to a substantial increase in detected loci.

**Fig. 4.**
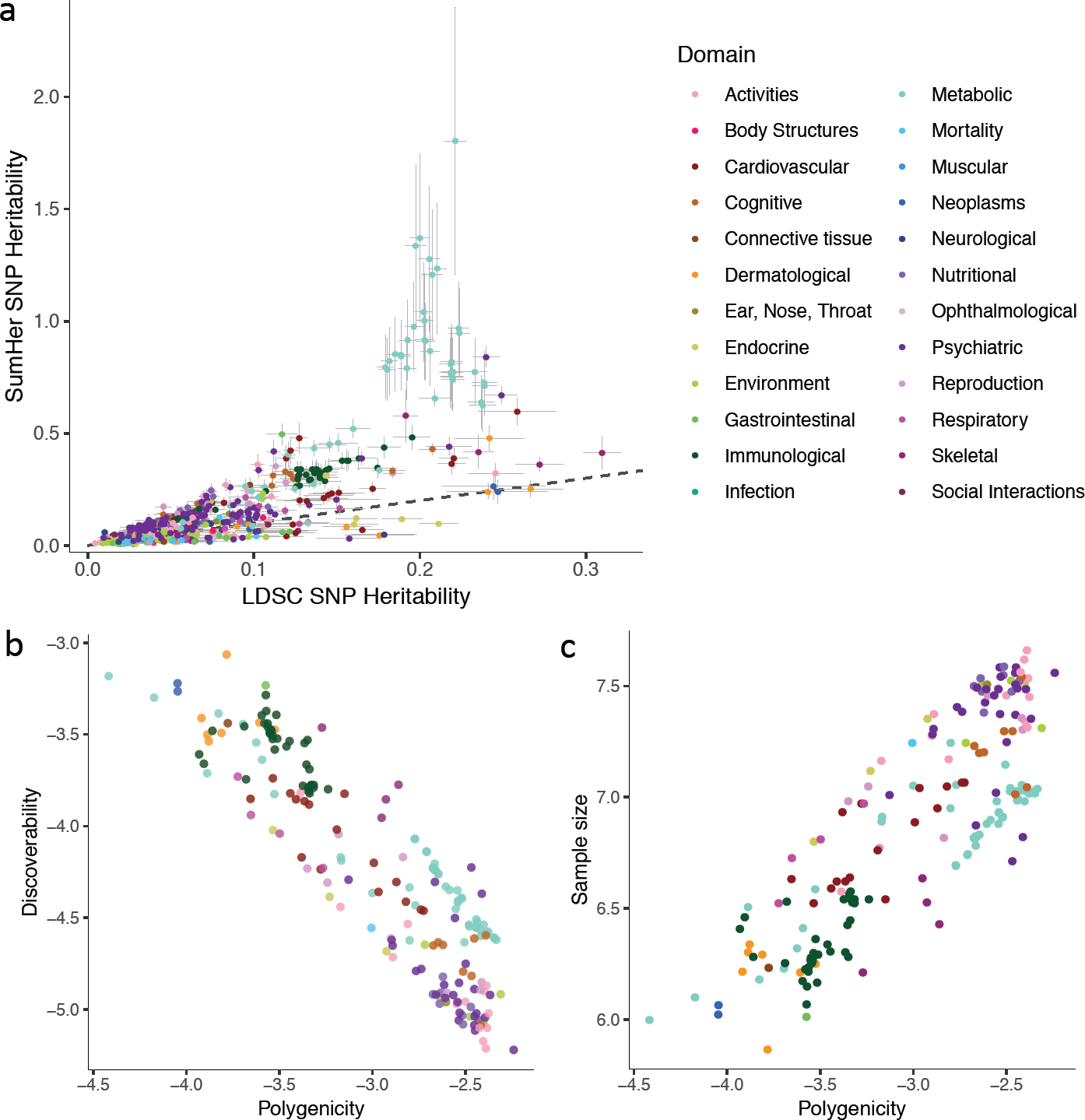
SNP heritability and polygenicity of 558 traits. **a.** Comparison of SNP heritability estimated by LDSC (x-axis) and SumHer (y-axis). Horizontal and vertical error bar represent standard errors of LDSC and SumHer estimates, respectively. **b.** Polygenicity and discoverability of traits, both on log 10 scale. Out of 558 traits, only 197 traits with reliable estimates (i.e. *h*^*2*^_*SNP*_>0.05 (estimated by MiXeR) and standard error of *π* is less than 50% of the estimated value) are displayed. Traits are colored by domain. **c.** Polygenicity and estimated sample size required to reach 90% of total SNP heritability explained by genome-wide significant SNPs, both in log 10 scale. Traits are colored by domain. Full results are available in **Supplementary Table 22**, **23**.

#### Polygenicity and discoverability of complex traits

The general observation from GWASs is that with increasing sample size, detected signals become not only more reliable but also more numerous, as with increasing power, smaller SNP effects may be detected. The total number of associated SNPs, the amount of variance they collectively represent, the distribution of effect sizes across the associated SNPs and how many additional individuals are expected to be needed for the detection of a fixed number of novel SNPs, are indications of the polygenicity of a trait. Such polygenicity may vary across traits, and can be informative for designing SNP-discovery studies.

To obtain an indication of trait-polygenicity, we applied the Causal Mixture Model for GWAS summary statistics (MiXeR)^28^ to estimate *π* (fraction of independent causal SNPs, polygenicity) and *σ_β_^2^* (variance of effect sizes of the causal SNPs, discoverability; see **Methods**). *π* ranges between 0 and 1, and a high *π* indicates a high level of polygenicity, while a high *σ_β_^2^* indicates a high level of discoverability of causal SNPs for the traits. Since the standard error of the model estimates become larger for traits with very small *h*^*2*^_*SNP*_ due to the small effect sizes, we only discuss the results of 197 out of 558 traits with *h*^*2*^_*SNP*_>0.05 and standard error of *π* less than 50% of the estimated value (as recommended by O. Frei; full results are available in **Supplementary Table 23**). We observed, as expected, a negative relationship between polygenicity and discoverability (*r*=-0.89 and *p*=4.93e-70), confirming that highly polygenic traits tend to have less causal SNPs with larger effect sizes (**Fig. 4b**).

The majority of traits (i.e. 116 traits) showed high polygenicity with *π*>1e-3 (more than 0.1% of all SNPs are causal). The highest polygenicity was observed in Major depressive disorder with 0.6% of SNPs being causal, while some traits, such as fasting glucose and serum urate level showed relatively low polygenicity (**Fig. 4b** and **Supplementary Table 23**). The traits with polygenicity >0.1% showed, on average, 8 times less discoverability compared to other traits with <0.1% of causal SNPs. The GWAS discoveries for traits with lower polygenicity and high discoverability will saturate with a lower sample size compared to the traits with higher polygenicity. Indeed, the estimated sample size, which is required to explain 90% of SNP heritability by genome-wide significant SNPs, is positively correlated with polygenicity (*r*=0.84 and *p*=6.30e-54), and extremely polygenic traits require tens of millions of subjects to identify 90% of causal SNPs at a genome-wide significant level (**Fig. 4c**).

## Discussion

The availability of hundreds of GWAS results provides the unique opportunity to gain insight into currently understudied questions regarding the genetic architecture of human traits. To facilitate such insight, we compiled a catalogue of 4,155 GWASs which can be queried online (http://atlas.ctglab.nl). We selected 558 well-powered GWASs to answer fundamental questions concerning the extent of pleiotropy of loci, genes, SNPs and gene-sets, characteristics of trait-associated variants and the polygenicity of traits.

We found that the total summed length of trait-associated loci for the 558 analysed traits covered more than half (60.1%) of the genome. 90% of the grouped loci contained associations with multiple traits across multiple trait domains. High locus pleiotropy can occur in two scenarios; *i*) when the same gene in a locus is associated with multiple traits or ii) when different genes or SNPs in the same locus are associated with multiple traits but due to LD the same locus is indicated. Our results showed that the proportion of pleiotropic associations dropped from 90% at the locus level to 63% at the gene level, and to 31% at the SNP level. These results show that although locus pleiotropy is widespread, pleiotropy at the level of genes and SNPs is much less abundant. This suggests that a gene can be involved in two distinct traits but how that gene is affected by the causal SNPs might differ. For instance, the function of the gene can be disrupted through a coding SNP for one trait, but expression of the same gene can be affected through a regulatory SNP for another trait.

Genes and SNPs that had a higher level of pleiotropy, were less tissue specific in terms of gene expression and active eQTLs. This suggests that SNPs and genes associated with multiple trait domains are more likely to be involved in general biological functions. Indeed, the top highly pleiotropic gene-sets were mostly involved in regulation of transcription which is an essential biological mechanism for any kind of cell to be functioning. Highly pleiotropic genes, therefore, can explain general vulnerability to a wide variety of traits, yet they may be less informative when the aim is to understand the causes of a specific trait. Although a large proportion of trait-associated genes are pleiotropic, the majority of trait associated gene-sets were trait-specific. Thus, the trait-specific combination of genes is highly informative, and future studies aimed at improved annotation of gene-functions will be needed to understand trait-specific gene association patterns.

It has been widely acknowledged that almost 90% of GWAS findings fall into non-coding regions^2^. Our results indeed show that 89.1% of the lead SNPs are non-coding, including intergenic (34.3%) and intronic (43.6%) SNPs. similarly, of the credible SNPs 92.4% were non-coding (intergenic 33.4% and intronic 48.1%). However, we showed different patterns when considering lead and credible SNPs; intergenic SNPs were depleted and the intronic SNPs were enriched in both the lead and credible SNPs. We also observed strong enrichment of the lead and credible SNPs in coding and flanking regions. These results indicate that both SNPs with the largest effect size (the lead SNPs) and the most likely causal SNPs (credible SNPs) within a locus tend to be located within or close to the genes. Although active chromatin states were enriched in both lead and credible SNPs, eQTLs were only enriched in credible SNPs but depleted in lead SNPs. This implies that likely causal regulatory SNPs do not necessarily have the strongest effect sizes in a locus.

Our analyses showed that the majority of analysed traits are highly polygenic with more than 0.1% of SNPs being causal. For those highly polygenic traits, over 10s of millions of individuals are required to identify all SNPs at genome wide significance (*p*<5e-8) that can explain at least 90% of the phenotypic variance explained by additive genetic effects. In the case of polygenic traits, individuals have almost unique combinations of risk/effect alleles for a specific disease or trait. With higher levels of polygenicity, and thus larger quantities of causal SNPs, the possible combinations of them also increase. This substantially increases the degree of genetic heterogeneity of the trait, and complicates the detection of genetic effects as the effect sizes of individual SNPs that are yet to be detected are even smaller than those observed in current GWASs.

In conclusion, our analyses have provided novel insight into the extent of pleiotropy, the nature of associated genetic regions and how traits differ in genetic architectures. This knowledge can guide the design of future genetic studies.

## METHODS

### Publicly available GWAS summary statistics

GWAS summary statistics were curated from multiple resources and were included only when the full set of SNPs were available. We excluded whole exome sequencing studies. This yielded 2,288 GWASs from 33 consortia and any other resources where summary statistics are available (last update 23^rd^ October 2018). From dbGAP, we obtained 2,659 unique datasets (ftp://ftp.ncbi.nlm.nih.gov/dbgap/Analyses_Table_of_Contents.txt, last accessed 4^th^ July 2017) and extracted 896 GWAS summary statistics in which a matched publication was available and sample size for a specific trait was explicitly mentioned in the original study. We excluded non-GWAS studies (e.g. PAGE (Prenatal Assessment of Genomes and Exomes) studies) and GWASs with immune-chip, whole exome sequencing and replication cohorts (exact reasons of exclusion for each dataset is available in **Supplementary Table 24**).

Together this resulted in a total of 3,555 GWAS summary statistics. The complete list and detailed information for each GWAS with summary statistics is available in **Supplementary Table 3** (atlas ID 1-3184, 3785-4155).

### UK Biobank GWAS summary statistics

Additional to the summary statistics available from external studies, we performed GWASs of traits from UK Biobank release 2 cohort (UKB2)^12^ under application ID 16404. We only used phenotype fields with first visit and first run (e.g. f.xxx.0.0) with exceptions for multi-coded phenotypes, which allowed to assign more than one code for a single subject (see **Supplementary Information 1, 2**). From the 1,940 unique field IDs to which we had access, 755 had >50,000 subjects with non-missing values. They are assigned to field name using ukb_field.tsv obtained from http://biobank.ctsu.ox.ac.uk/crystal/download.cgi (last accessed 31^st^ August 2017). Note that for newly available phenotypes for release 2, we annotated field names manually based on the UK biobank data showcase. From these phenotypes, we excluded baseline characteristics, phenotypes used as covariates, date and place phenotypes, status phenotypes (i.e. completion status, answered a specific question), ethnicity, genomic phenotypes and any other phenotypes that are not relevant for performing a GWAS. For each phenotype, we provided reason of exclusions in **Supplementary Table 1**. This resulted in 434 unique fields including 49 multi-coded phenotypes. 385 phenotypes were considered quantitative when the phenotype value was quantitative or categorical, and could be ordered. Phenotypes coded by yes/no were considered as binary with a few exceptions (**Supplementary Table 1**). For quantitative and binary phenotypes, subjects with phenotype codes −1 for “Do not know” or −3 for “Prefer to not answer” were excluded and the original phenotype code as described in the UK biobank data showcase was used unless specified in Supplementary Text or **Supplementary Table 1, 2**. For 49 multi-coded phenotypes, we dichotomized each code to dummy binary phenotypes (cases for 1 and controls for 0) and included subjects with phenotype code −7 for “None of the above” as controls. Again, subjects with phenotype codes −1 for “Do not know” or −3 for “Prefer to not answer” were excluded. For example, field 670 based on UKB Data-Coding 100286 is coded from 1 to 5 and dichotomization results in five phenotypes such as 1 vs all others, 2 vs all others and so on. Detailed definitions of multi-coded phenotypes are described in **Supplementary Table 2**. After phenotyping, we selected phenotypes that had at least 50,000 European subjects. For binary traits, we further restricted to traits with at least 10,000 cases and controls. This resulted in a total of 600 traits (260 quantitative and 340 binary traits). Note that the final total sample size encoded in the atlas database (http://atlas.ctglab.nl) might be less than 50,000 due to lack of genotype data or missing values in covariates.

GWAS was performed for up to 10,846,944 SNPs with MAF > 0.0001 using PLINK 2^29^, while correcting for array, age (f.54.0.0), sex (f.31.0.0), Townsend deprivation index (f.189.0.0), assessment centre (f.21003.0.0) and 20 PCs. Linear or logistic models were used for quantitative or binary traits, respectively.

The complete list of traits from UK biobank release 2 analysed in this study is available in **Supplementary Table 3** (atlas ID 3185-3784).

### Pre-processing of GWAS summary statistics

Curated summary statistics were pre-processed to standardize the format. SNPs with p≤0 or >1, or non-numeric values such as “NA” were excluded. For summary statistics with non-hg19 genome coordinates, liftOver software was used to align to hg19. When only rsID was available in the summary statistics file without chromosome and position, genome coordinates were extracted from dbSNP 146. When rsID was missing, it was assigned based on dbSNP 146. When only the effect allele was reported, the other allele was extracted from dbSNP 146.

### Definition of lead SNPs and trait-associated loci

For each GWAS, we defined lead SNPs and genomic trait-associated loci as described before ^30^. First, we defined independent significant SNPs with *p*<5e-8 and independent at *r*^*2*^<0.6, and defined LD blocks for each of independent significant SNPs based on SNPs with p<0.05. Of these SNPs, we further defined lead SNPs that are independent at *r*^*2*^<0.1. We finally defined genomic trait-associated loci by merging LD blocks closer than 250kb. Each trait-associated locus was then represented by the top SNP (with the minimum P-value) and its genomic region was defined by the minimum and maximum position of SNPs which are in LD (*r*^*2*^≥0.6) with one of the independent significant SNPs within the (merged) locus. We used 1000 genome phase 3 (1000G)^31^ as a reference panel to compute LD for most of the GWASs in the database. For each GWAS, the matched population (from AFR, AMR, EAS, EUR, SAS) was used as the reference based on the information obtained from the original study. For trans-ethnic GWASs, the population with the largest total sample size was used. When the GWAS was based on the UKB release 1 cohort (UKB1), we used 10,000 randomly sampled unrelated White British subjects from UKB1 as reference. For other GWASs performed in this study or GWASs based on the UKB2, 10,000 randomly selected unrelated EUR subjects were used as a reference. Non-bi-allelic SNPs were excluded from any analyses.

The reference panel used for each GWAS is provided in the column “Population” of **Supplementary Table 3**. For trans-ethnic GWASs, the first population was used as reference, e.g. EUR+EAS+SAS means EUR had the largest sample. GWASs based on the UKB cohort was encoded either “UKB1 (EUR)” for UKB release 1 or “UKB2 (EUR)” for UKB release 2.

### MAGMA gene and gene-set analysis

We performed MAGMA v1.06^16^ gene and gene-set analyses for every GWAS in the database. For gene-analysis, 20,260 protein-coding genes were obtained using the R package BioMart (Ensembl build v92 GCRh37). SNPs were assigned to genes with 1kb window at both sides. The reference panel of corresponding populations used for each GWAS was based on either 1000G, UKB1 or UKB2 as described in the previous section. The gene-set analysis was performed with default parameters (snp-wise mean model). Gene-set analysis was performed for 4,737 curated gene-sets (C2) and 5,917 GO terms (C5; 4,436 biological processes, 580 cellular components and 901 molecular functions) from MsigDB v6.1 (http://software.broadinstitute.org/gsea/msigdb, last accessed 20 Apr 2018)^32^.

### SNP heritability and genetic correlation with LD score regression

We performed LD score regression (LDSC)^17^ for each GWAS to obtain SNP heritability and pairwise genetic correlations. Pre-calculated LD scores for 1000G EUR and EAS populations were obtained from https://data.broadinstitute.org/alkesgroup/LDSCORE/ (last accessed 26 Nov 2016) and LD score regression was only performed for GWASs with either an EUR or EAS population and when the number of SNPs in the summary statistics file was > 450,000. LDSR was performed only for HapMap3 SNPs excluding the MHC region (25Mb-34Mb). When the signed effect size or odds ratio was not available in the summary statistics file, “-- a1-inc” flag was used. As recommended previously^33^, we excluded SNPs with chi-square >80. For binary traits, the population prevalence was curated from the literature (only for diseases whose prevalence was available, **Supplementary Table 25**) to compute SNP heritability at the liability scale with “--samp-prep” and “--pop-prep” flags. For most of the personality/activity (binary) traits from UKB2 cohort, we assumed that the sample prevalence is equal to the population prevalence since the UK Biobank is a population cohort and not designed to study a certain disease/traits. Likewise, when population prevalence was not available, sample prevalence was used as population prevalence for all other binary traits. Genetic correlations were computed for pair-wise GWASs with the following criteria as suggested previously^33^:

- GWASs of EUR population or more than 80% of samples are EUR.
- The number of SNPs >450,000
- Signed effect size or odds ratio is available
- Effect and non-effect alleles are explicitly mentioned in the header or elsewhere.
- SNP heritability Z score >2

In total, pairwise genetic correlations were computed for 1,090 GWASs in the database.

### Selection of GWASs for cross-phenotype analyses

From the 4,155 curated GWASs in the database, we selected 558 GWASs with unique traits for cross-phenotype analyses based on the following criteria.

- Minimum sample size 50,000 and both cases and controls are >10,000 for binary phenotypes.
- The number of SNPs in the summary statistics is >450,000.
- GWAS is based on EUR population or >80% of the samples are EUR. If summary statistics of both trans-ethnic and EUR-only are available, use EUR-only GWAS.
- Exclude sex-specific GWAS, unless the phenotype under study is only available for a specific sex (e.g., age at menopause). If sex-specific and sex-combined GWASs are available, use sex-combined GWAS.
- Z-score of *h*^*2*^_*SNP*_ computed by LDSC is >2
- Signed effect size (beta or odds ratio) is available in the summary statistics.
- Effect and non-effect alleles are explicitly mentioned in the header or elsewhere.
- From GWASs that met the above criteria, we selected a GWAS per trait with the maximum sample size.

UKB2 GWASs performed in this study are further filtered based on the following:

- Exclude cancer screening or test phenotypes.
- Exclude item level phenotypes (i.e., Neuroticism and Fluid intelligence tests)
- Exclude phenotypes of parents’ age and parents’ still alive.
- Exclude medication, treatment, supplements and vitamin traits.
- If exactly the same traits were diagnosed by an expert (e.g. doctor) and self-reported, use the expert qualification.
- If exactly the same traits were present as main and secondary diagnoses, both are included.
- Phenotypes with large extremes were excluded from the analyses when the difference between the maximum value and 99 percentiles of the standardized phenotype value is >50.

There was one exception for height GWAS, where a meta-analysis by Yengo et al.^34^ (ID 4044) has the larger sample size, however the meta-analysis was limited to ∼2.4 million HapMap 2 SNPs. Since over 10 million SNPs are included in most of the selected GWASs, this smaller number of SNPs can bias our analyses. Therefore, the second largest GWAS (UKB2 GWAS performed in this study, ID 3187) was used instead. This resulted in total of 558 GWASs, across 24 domains, which were subsequently used in the cross-phenotype analyses in this study. These 558 GWASs are specified in **Supplementary Table 3**.

### Pleiotropic trait-associated loci

To define pleiotropic loci for the 558 traits (GWASs), we first extracted trait-associated loci on autosomal chromosomes. We excluded any locus with a single SNP (no other SNPs have *r*^*2*^>0.6) as these loci are more likely to be false positives. We then grouped physically overlapping loci across 558 traits. In a group of loci, it is not required that all individual trait-associated loci are physically overlapping but merging them should result in a continuous genomic region. For example, when trait-associated loci A and B physically overlap and trait-associated loci B and C also physically overlap, but A and C do not, these three trait-associated loci were grouped into a single group of loci (**Extended Data Fig. 3**). Therefore, a grouped locus could contain more than one independent locus from a single trait when gaps between them were filled by loci from other traits. The grouped loci were further assigned to three categories, *i*) multi-domain locus when a loci group contained traits from more than one domain, *ii*) domain specific locus when a loci group contained more than one trait from the same domain, and *iii*) trait specific locus when a locus did not overlap with any other loci. We compared the distribution of gene density across four association categories of the loci; multi-domain, domain specific and trait specific loci, and non-associated genomic regions. To define non-associated genomic regions, we extracted the minimum and maximum positions that were covered by 1000G, and the gap regions of grouped trait-associated loci were defined as non-associated regions. The gene density was computed as a proportion of a region that was overlapping with one of 20,260 protein-coding genes obtained from Ensembl v92 GRCh37. We then performed pairwise Wilcoxon rank sum test (two sided).

### Colocalization of trait-associated loci

To evaluate if physically overlapping trait-associated loci also share the same causal SNPs, we performed colocalization using the *coloc.abf* (Approximate Bayes Factor colocalization analysis) function of the coloc package in R^35^. Colocalization analysis was performed for all possible pairs of physically overlapping trait-associated loci across 558 traits. When two loci from different traits were physically overlapping but there were no SNPs that were present in both GWAS summary statistics in that overlapping region, colocalization was not performed. The inputs of the *coloc.abf* function are P-value, MAF and sample size for each SNP. When MAF was not available in the original summary statistics, it was extracted from the matched reference panel. For binary traits, sample prevalence was additionally provided based on total cases and controls of the study.

The *coloc.abf* function assumes a single causal SNP for each trait and estimates the posterior probability of the following 5 scenarios for each testing region; *H*_*0*_: neither trait has a genetic association, *H*_*1*_: only trait 1 has a genetic association, *H*_*2*_: only trait 2 has a genetic association, *H*_*3*_: both trait 1 and 2 are associated but with different causal SNPs and *H*_*4*_: both trait 1 and 2 are associated with the same single causal SNP. In this study, as we pre-define the trait-associated loci for each trait which already discard scenarios *H*_*0*_ to *H*_*2*_, we are only interested whether *H*_*4*_ is most likely. We therefore defined, a pair of loci as colocalised when the posterior probability of *H*_*4*_ is >0.9. We note that it is possible that genomic regions outside of the pre-defined trait-associated loci can also colocalize with other traits. However, we limited the analyses to the pre-defined trait-associated loci in this study, to be consistent with the level of pleiotropy measured by physical overlap of the loci.

Within a grouped locus defined based on physical overlap (see above), we further grouped loci based on a colocalization pattern. To do so, we considered colocalization pattern across group of physically overlapping loci as a graph in which nodes represent trait-associated loci and edges represent colocalization of the loci First, loci which did not colocalized with any other loci were considered as independent loci. For the rest of the loci, we identified connected components of the graph (**Extended Data Fig. 3**). This does not require all loci within a component to be colocalized with each other. For example, when locus A is colocalized with locus B, and locus B is colocalized with locus C, but locus A is not colocalized with locus C, all loci A, B and C are grouped into a single connected component. Detailed results are discussed in the **Supplementary Information 3**.

### Pleiotropic genes

For gene level pleiotropy, we extracted MAGMA gene analysis results for the 558 traits where 17,444 genes on autosomal chromosomes were tested in all GWASs. For each trait, genes with *p*<2.87e-6 (0.05/17,444) were considered as significantly associated. We did not correct the P-value for testing 558 traits since our purpose is not to identify genes associated with one of the 558 traits but to evaluate the overlap of trait-associations (when GWAS was performed for a single trait) across the 558 traits, and this applies to SNPs and gene-set level pleiotropy. The trait associated genes were further categorized into three groups in a similar way as for trait-associated loci, i.e. *i*) multi-domain genes that were significantly associated with traits from more than one domain, *ii*) domain-specific genes that were significantly associated with more than one trait from the same domain and *iii*) trait-specific genes that were significantly associated with a single trait.

We compared gene length and pLI score across genes in three different association categories and non-associated genes. Gene length was based on the start and end position of genes extracted from the R package biomaRt and pLI score was obtained from ftp://ftp.broadinstitute.org/pub/ExAC_release/release0.3.1/functional_gene_constraint (last accessed 27 April 2017). We performed t-tests for gene length in log scale and Wilcoxon rank sum tests for pLI scores (both two sided).

For each protein coding gene, we first assessed whether a gene is expressed or not in each of 53 tissue types based on expression profile obtained from GTEx v7^20^. We defined genes as expressed in a given tissue type if the average TPM is >1. For each of 17,444 genes, we then counted the number of tissue types where the gene is expressed and grouped them into six categories, i.e. genes expressed in *i*) a single tissue type (tissue specific genes), *ii*) between 2 and 13, *iii*) between 14 and 26, *ix*) between 27 and 39, *x*) between 40 and 52, and *xi*) 53 (all) tissue types. At each number of associated domains (from 1 to 10 or more domains), we recalculated the proportion of genes in each of the 6 categories, and performed the Fisher’s exact tests (one-sided) against baseline (the proportion relative to all 17,444 genes) to evaluate if the proportion is higher than expected.

### Pleiotropic SNPs

We extracted 1,740,179 SNPs that were present in all 558 GWASs. To evaluate if the select ion of ∼1.7 million SNPs biased the results, we compared distribution of these analysed SNPs with the all known SNPs in the genome (SNPs exist in 1000G EUR population, UKB1 and UKB2 reference panels) by computing the proportion of SNPs per chromosome. In addition, distribution of functional consequences of SNPs annotated by ANNOVAR^36^ was also compared with the all SNPs in the genome. For each SNP, we counted the number of traits to which a SNP was significantly associated at *p*<5e-8, and then grouped the associated SNPs into multi-domain, domain-specific and trait-specific SNPs using the same definitions as at the gene level.

Functional consequences of SNPs were annotated using ANNOVAR^36^. To test if a SNP from a certain functional category is enriched at a given number of associated domains compared to all analysed SNPs, a baseline proportion was calculated from the 1,740,179 SNPs for each functional category. At each number of associated domains (from 1 to 10 or more domains), we re-calculated the proportion of SNPs with each functional category and performed the Fisher’s exact test (one-sided) against the baseline (the proportion relative to all 1,740,179 SNPs), to test if the proportion if higher than expected.

eQTLs for 48 tissue types were obtained from GTEx v7 (https://www.gtexportal.org/home/; last accessed 20 January 2018)^20^ and we considered SNPs with gene q-value <0.05 with any gene in any tissue as eQTLs. For each eQTL, we counted the number of tissue types of being eQTL (regardless of associated genes) and categorized them into five groups, i.e. being eQTLs in *i*) a single tissue type (tissue specific eQTLs), *ii*) between two and 12, *iii*) between 13 and 24, *ix*) between 25 and 36 and *x*) and being in more than 37 tissue types. At each number of associated domains, we re-calculated the proportion of SNPs in each of the 5 categories, and performed the Fisher’s exact test (one-sided) against baseline (the proportion relative to all 1,740,179 SNPs), to test if the proportion if higher than expected.

### Pleiotropic gene-sets

For gene-set level pleiotropy, we extracted 10,650 gene-sets tested in all 588 traits. We then considered gene-sets with *p*<4.69e-6 (0.05/10,650) as significantly associated. The trait associated gene-sets were grouped into multi-domain, domain-specific and trait-specific gene-sets with the same definitions as at the gene level.

We compared the number of genes and average gene-length across gene-sets in different association categories and non-associated genes. Gene length was based on the start and end position of genes extracted from R package, biomaRt. We performed two-sided t-test in log scale of the number of genes and average gene-length.

### Power calculation of genetic correlation

Power calculations were performed using the bivariate analysis of GCTA-GRML power calculator (http://cnsgenomics.com/shiny/gctaPower/)^37^, to estimate the minimum *r_g_* that obtain a power of 0.8 in the worst case scenario. From 558 traits, two traits with the worst case scenarios were selected, one with the minimum *h*^*2*^_*SNP*_ estimated by LDSC and another with the minimum sample size. For each case, we obtained the minimum *r_g_* to obtain power of 0.8 by assuming both traits are quantitative with same sample size and *h*^*2*^_*SNP*_ and have phenotypic correlation 0.1.

### Hierarchical clustering of trait based on genetic correlation

Hierarchical clustering was performed on the matrix of pair-wise *r_g_*’*s* as calculated between the 558 traits. After Bonferroni correction for all possible trait pairs, non-significant genetic correlations were replaced with 0. The number of clusters *k* was optimized between 50 and 250 by maximizing the silhouette score with 30 iterations for each *k*.

### Estimated standardized effect size of lead SNPs

To enable comparison of effect sizes across GWASs from different studies, we first converted P-values into Z-statistics (two sided) and expressed the estimated effect size as a function of MAF and sample size as described previously^21^ using the following equations:

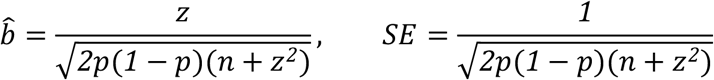

where *p* is MAF and n is the total sample size. We used the MAF of a corresponding European reference panel (either 1000G, UKB1 or UKB2) as described in the previous section “Definition of lead SNPs and genomic trait-associated loci”. Since we were not interested in the direction of effect, we used squared standardized effect sizes for analyses in this study.

### Fine-mapping of trait-associated loci

We defined the region to fine-map by taking 50kb around the top SNPs of the trait-associated loci. When trait-associated loci were larger than the 50kb window, the largest boundary was taken. Due to the complex LD structure, loci overlapping with the MHC region (chr6:25Mb-36Mb) were excluded. The fine-mapping was performed using the FINEMAP software (http://www.christianbenner.com/#) with shotgun stochastic search algorithm^25^. Since the coverage of true causal SNPs is affected by the sample size of the reference panel and GWASs^38^, we used randomly selected unrelated 100k EUR individuals from UKB2 cohort for all 558 GWASs. We limited the number of maximum causal SNPs (*k*) per locus to 10. When the number of SNPs within a locus is relatively small (around 30 or less), the algorithm can fail to converge. In that case, k was decreased by 1 until FINEMAP was successfully run. Loci with less than 10 SNPs were excluded from the fine-mapping.

FINEMAP outputs a set of models (all possible combination of *k* causal SNPs in a locus) with posterior probability (PP) of being a causal model. A 95% credible set was defined by taking models from the highest PP until the cumulative sum of PP reached 0.95. Then 95% credible set SNPs were defined as unique SNPs included in the 95% credible set of models. For each SNP, a posterior inclusion probability (PIP) was calculated as the sum of PPs of all models that contains that SNP. To select most likely causal SNPs, we further defined credible SNPs consists of SNPs with PIP>0.95. Detailed results are discussed in **Supplementary Information 9.**

### Annotation and characterization of lead SNPs and credible SNPs

Functional consequences of SNPs were annotated using ANNOVAR^36^ based on Ensembl gene annotations on hg19. Prior to ANNOVAR, we aligned the ancestral allele with dbSNP build 146. 15-core chromatin states of 127 cell/tissue types were obtained from Roadmap^39^ (http://egg2.wustl.edu/roadmap/data/byFileType/chromhmmSegmentations/ChmmModels/coreMarks/jointModel/final/all.mnemonics.bedFiles.tgz; last accessed 16 Mar 2016) and we annotated one of the 15-core chromatin states to each of the lead SNPs based on chromosome coordinates. Subsequently, consequence state was assigned for each SNP by taking the most common state across 127 cell/tissue types. SNPs with consequence state≤7 were defined as active. eQTLs in 48 tissue types were obtained from GTEx v7^20^ and we only used the significant eQTLs at gene q-value<0.05. eQTLs were assigned to SNPs by matching chromosome coordinate and alleles.

As we showed that trait-associated loci have higher gene density compared to non-associated regions, and GWAS signals are known to be enriched in regulatory elements^40^, we first identified background enrichment by comparing SNPs within trait-associated loci or fine-mapped regions with the entire genome. For this all known SNPs were extracted by combining all SNPs in 1000G, UKB1 and UKB2 reference panels (∼28 million SNPs in total). SNPs within the trait-associated loci were defined as the ones with P-value<0.05 and *r*^*2*^>0.6 with one of the independent significant SNPs as described above (see section ‘Definition of lead SNPs and trait-associated loci’). Therefore, it does not necessary include all SNPs physically located within the trait-associated loci. On the other hand, SNPs within fine-mapped region include all SNPs physically located within 50kb window from the most significant SNP of a locus. To characterize lead SNPs and credible SNPs given background enrichments, we compared these SNPs against all SNPs within trait-associated loci or fine-mapped regions, respectively.

### SNP heritability estimation with SumHer using LDAK model

We estimated SNP heritability of 558 traits using the SumHer function from the LDAK software v5.0 (http://dougspeed.com/ldak/) ^27^. Since our purpose was to compare estimates from LDSC and SumHer, we used the 1000G EUR reference panel and extracted HapMap3 SNPs as consistent with LDSC. We used unique ID’s of SNPs (consisting of chromosome:posision:allele 1:allele2) instead of rsID to maximize the match between GWAS summary statistics and the reference panel. The MHC region (chr6:25Mb-34Mb) was excluded. As recommended by the author, SNPs with large effects (*Z*^*2*^/(*Z*^*2*^+*n*)>100 where *Z*^*2*^ is chi-squared statistics and *n* is sample size of the SNP) were excluded.

To obtain SNP heritability in a liability scale, we provided population prevalence and sample prevalence with flags ‘--prevalance’ and ‘--ascertainment’ for binary traits. The same population prevalence was used as described in the section of SNP heritability estimate with LDSC (**Supplementary Table 25**). Details results are discussed in **Supplementary Information 10**.

### Estimation of polygenicity and discoverability with MiXeR

In the causal mixture model for GWAS summary statistics (MiXeR) proposed by Holland et al., the distribution of SNP effect sizes is treated a mixture of two distributions for causal and non-causal SNPs as the following^28^:

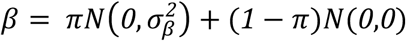

where *π* is the proportion of (independent) causal SNPs and *σ_β_^2^* is the variance of the effect sizes of causal SNPs. Therefore, *π* and *σ_β_^2^* respectively represent polygenicity and discoverability of the trait. We estimated both parameters for the 558 traits using MiXeR software (https://github.com/precimed/mixer)^28^. As recommended in the original study, we used 1000G EUR as a reference panel and restricted to HapMap 3 SNPs. SNPs with χ^2^>80 and the MHC region (chr6:26Mb-34Mb) were excluded. To estimate the sample size required to explain 90% of the additive genetic variance of a phenotype, we used an output of GWAS power estimates calculated in the MiXeR software, which contains 51 data points of sample size and the proportion of chip heritability explained^28^. We then estimated the sample size required to reaches 90% by using the *interp1* function from the pracma package in R.

## Data and materials availability

All publicly available GWAS summary statistics (original) files curated in this study are accessible from the original links provided at http://atlas.ctglab.nl. GWAS summary statistics for 600 traits from UK Biobank performed in this study are also provided at http://atlas.ctglab.nl and an archived file will be made available upon publication from https://ctg.cncr.nl/software/summary_statistics.

## END NOTES

### Acknowledgement

We thank all consortiums and all other individual labs for making GWAS summary statistics publicly available. We also thank Peter Visscher and Naomi Wray for their thoughtful suggestions and discussions. We additionally thank Anders Dale for his suggestions for the manuscript. This work was funded by Netherlands Organization for Scientific Research (NWO VICI 453-14-005 and NWO VIDI 452-12-014).

## Author contribution

D.P. designed the study. K.W. curated the database and performed analyses. T.J.C.P assisted with harmonization of phenotype labels of the database. S.S. performed QC on the UK Biobank data and wrote the analysis pipeline for UKB analyses. M.U.M assisted with the fine-mapping analyses. O.F. and O.A.A. developed software UGMG and assisted with the analyses. S.v.d.S and B.M.N discussed and provided valuable suggestions for analyses. K.W. and D.P. wrote the paper. All authors critically reviewed the paper.

## REFERENCES

1. Edwards, A. O. et al. Complement factor H polymorphism and age-related macular degeneration. Science (80-.). 308, 421–425 (2005).

2. Welter, D. et al. The NHGRI GWAS Catalog, a curated resource of SNP-trait associations. Nucleic Acids Res. 42, D1001–D1006 (2014).

3. Lander, E. S. Initial impact of the sequencing of the human genome. Nature 470, 187–197 (2011).

4. Visscher, P. M. et al. 10 Years of GWAS Discovery?: Biology, Function, and Translation. Am. J. Hum. Genet. 101, 5–22 (2017).

5. Henderson, P. & Stevens, C. The role of autophagy in Crohn’s Disease. Cells 1, 492–519 (2012).

6. Okada, Y. et al. Genetics of rheumatoid arthritis contributes to biology and drug discovery. Nature 506, 376–81 (2014).

7. Gaulton, K. J. et al. Genetic fine mapping and genomic annotation defines causal mechanisms at type 2 diabetes susceptibility loci. Nat. Genet. 47, 1415–1425 (2015).

8. Canela-Xandri, O., Rawlik, K. & Tenesa, A. An atlas of genetic associations in UK Biobank. Nat. Genet. 50, 1593–1599 (2018).

9. Timpson, N. J., Greenwood, C. M. T., Soranzo, N., Lawson, D. J. & Richards, J. B. Genetic architecture: The shape of the genetic contribution to human traits and disease. Nat. Rev. Genet. 19, 110–124 (2018).

10. Boyle, E. A., Li, Y. I. & Pritchard, J. K. An expanded view of complex traits: from polygenic to omnigenic. Cell 169, 1177–1186 (2017).

11. Wray, N. R., Wijmenga, C., Sullivan, P. F., Yang, J. & Visscher, P. M. Common disease is more complex than implied by the core gene omnigenic model. Cell 173, 1573–1580 (2018).

12. Bycroft, C. et al. The UK Biobank resource with deep phenotyping and genomic data. Nature 562, 203–209 (2018).

13. Goh, K. et al. The human disease network. Proc. Natl. Acad. Sci. 104, 8685–8690 (2007).

14. Polderman, T. J. C. et al. Meta-analysis of the heritability of human traits based on fifty years of twin studies. Nat. Publ. Gr. 47, 702–709 (2015).

15. Mahajan, A. et al. Fine-mapping type 2 diabetes loci to single-variant resolution using high-density imputation and islet-specific epigenome maps. Nat. Genet. (2018).

16. de Leeuw, C. A., Mooij, J. M., Heskes, T. & Posthuma, D. MAGMA: generalized gene-set analysis of GWAS data. PLoS Comput. Biol. 11, e1004219 (2015).

17. Bulik-sullivan, B. K. et al. LD Score regression distinguishes confounding from polygenicity in genome-wide association studies. Nat. Genet. 47, 291–295 (2015).

18. Solovieff, N., Cotsapas, C., Lee, P. H., Purcell, S. M. & Smoller, J. W. Pleiotropy in complex traits: Challenges and strategies. Nat. Rev. Genet. 14, 483–495 (2013).

19. Lek, M. et al. Analysis of protein-coding genetic variation in 60,706 humans. Nature 536, 285–291 (2016).

20. The GTEx Consortium. Genetic effects on gene expression across human tissues. Nature 550, 204–213 (2017).

21. Zhu, Z. et al. Integration of summary data from GWAS and eQTL studies predicts complex trait gene targets. Nat. Genet. 48, 481–487 (2016).

22. Manolio, T. A. et al. Finding the missing heritability of complex diseases. Nature 461, 747–753 (2009).

23. Lee, S., Abecasis, G. R., Boehnke, M. & Lin, X. Rare-variant association analysis: Study designs and statistical tests. Am. J. Hum. Genet. 95, 5–23 (2014).

24. van de Bunt, M., Cortes, A., Brown, M. A., Morris, A. P. & McCarthy, M. I. Evaluating the performance of fine-mapping strategies at common variant GWAS loci. PLoS Genet. 11, e1005535 (2015).

25. Benner, C. et al. FINEMAP?: efficient variable selection using summary data from genome-wide association studies. Bioinformatics 32, 1493–1501 (2016).

26. Speed, D. et al. Reevaluation of SNP heritability in complex human traits. Nat. Genet. 49, 986–992 (2017).

27. Speed, D. & Balding, D. J. Better estimation of SNP heritability from summary statistics provides a new understanding of the genetic architecture of complex traits. Nat. Genet. (2018).

28. Holland, D. et al. Beyond SNP heritability: polygenicity and discoverability estimated for multiple phenotypes with a univariate gaussian mixture model. bioRxiv (2018).

29. Purcell, S. et al. PLINK: A tool set for whole-genome association and population-based linkage analyses. Am. J. Hum. Genet. 81, 559–575 (2007).

30. Watanabe, K., Taskesen, E., van Bochoven, A. & Posthuma, D. Functional mapping and annotation of genetic associations with FUMA. Nat. Commun. 8, 1826 (2017).

31. Auton, A. et al. A global reference for human genetic variation. Nature 526, 68–74 (2015).

32. Liberzon, A. et al. Molecular signatures database (MSigDB) 3.0. Bioinformatics 27, 1739–1740 (2011).

33. Zheng, J. et al. LD Hub: a centralized database and web interface to perform LD score regression that maximizes the potential of summary level GWAS data for SNP heritability and genetic correlation analysis. Bioinformatics 33, 272–279 (2017).

34. Yengo, L. et al. Meta-analysis of genome-wide association studies for height and body mass index in ∼700000 individuals of European ancestry. Hum. Mol. Genet. 27, 3641–3649 (2018).

35. Giambartolomei, C. et al. Bayesian Test for Colocalisation between Pairs of Genetic Association Studies Using Summary Statistics. PLoS Genet. 10, e1004383 (2014).

36. Wang, K., Li, M. & Hakonarson, H. ANNOVAR: functional annotation of genetic variants from high-throughput sequencing data. Nucleic Acids Res. 38, e164 (2010).

37. Visscher, P. M. et al. Statistical power to detect genetic (co)variance of complex traits using SNP data in unrelated samples. PLoS Genet. 10, e1004269 (2014).

38. Benner, C. et al. Prospects of fine-mapping trait-associated genomic regions by using summary statistics from genome-wide association studies. Am. J. Hum. Genet. 101, 539–551 (2017).

39. Roadmap Epigenomics Consortium. Integrative analysis of 111 reference human epigenomes. Nature 518, 317–330 (2015).

40. Tak, Y. G. & Farnham, P. J. Making sense of GWAS: using epigenomics and genome engineering to understand the functional relevance of SNPs in non-coding regions of the human genome. Epigenetics Chromatin 8, 57 (2015).

