## Extended Data for "A global overview of pleiotropy and genetic architecture in complex traits"

**Extended Data Figures**

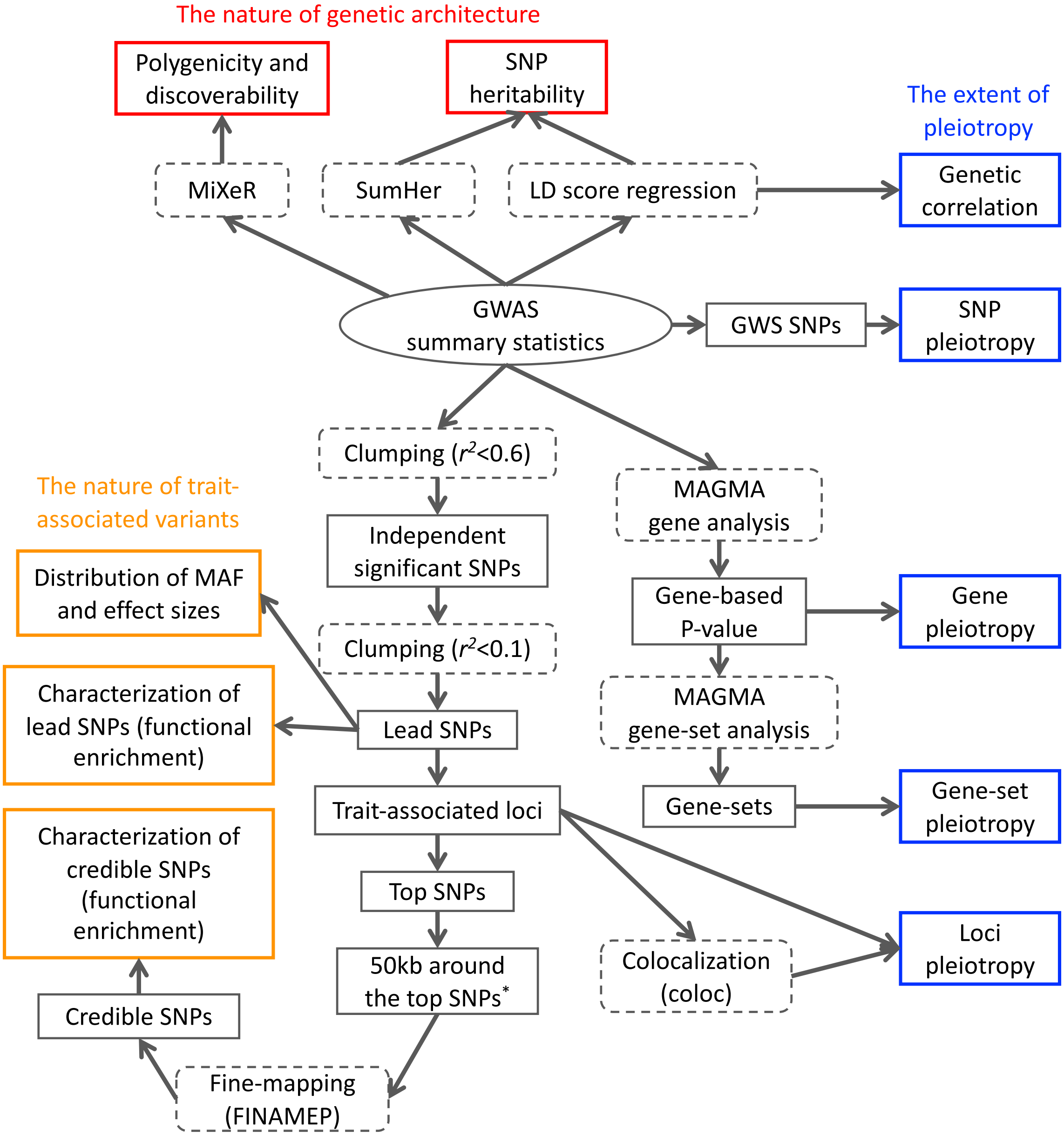

**Fig. 1 Schematic overview of the analyses**
Dashed grey boxes represent processes and/or software. Solid grey boxes represent results or outputs of the processes. Colored boxes are main analyses in this study, each corresponds a sub-headings in the main text. *Fine-mapped regions were defined by taking the largest range of 50kb around the top (most significant) SNPs and trait-associated loci.

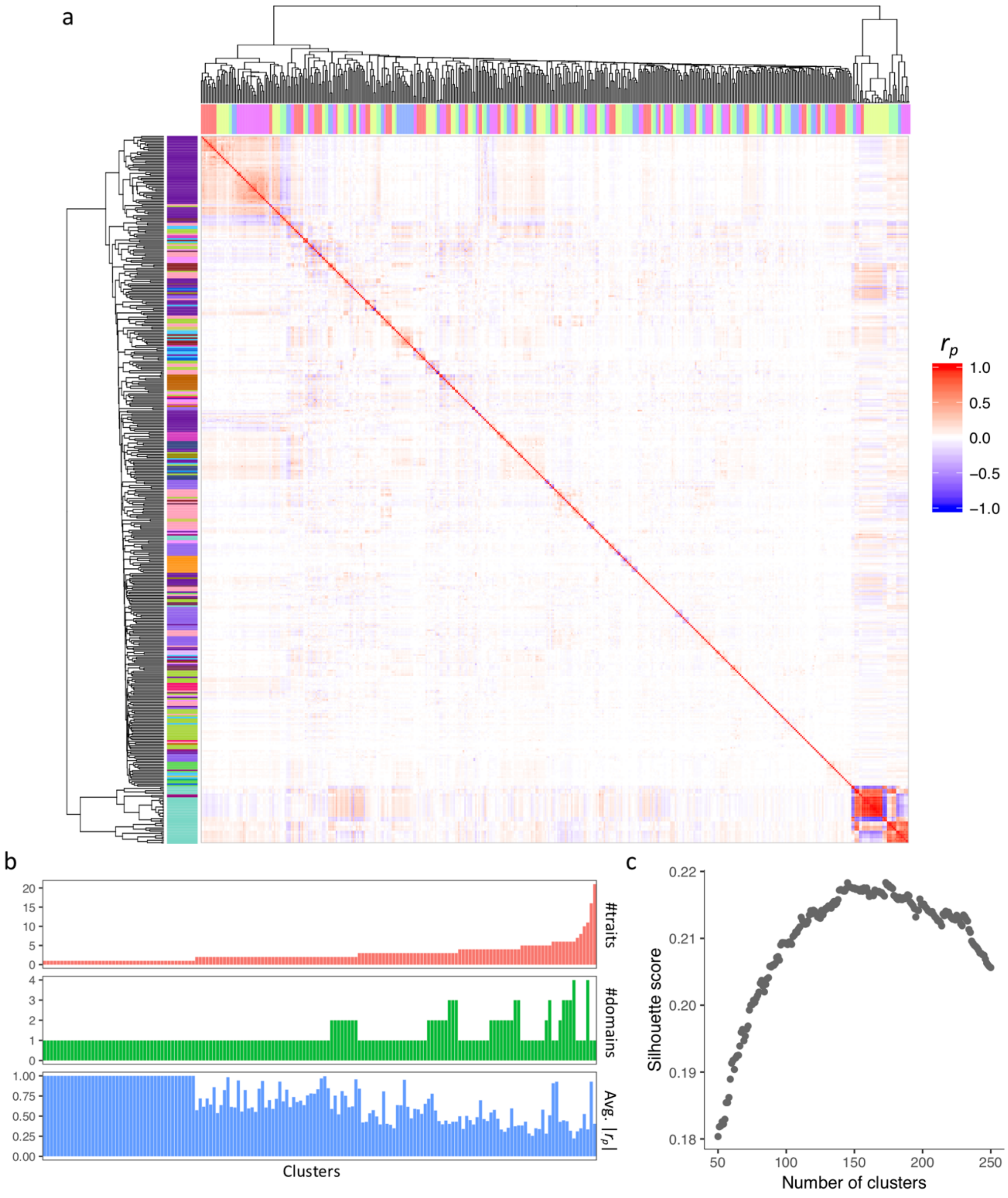

**Fig. 2. Phenotypic correlation across 457 UKB2 phenotypes. a**. Heatmap of pair-wise phenotypic correlation matrix. Both the columns and rows are hierarchically clustered. The colors at the top (x-axis) represent the clusters (consecutive traits with the same color are in the same cluster) and the colors at the left (y-axis) represent the domain of the trait. **b.** Number of traits, domains and average phenotypic correlation per cluster. **c**. Silhouette score with different number of clusters.

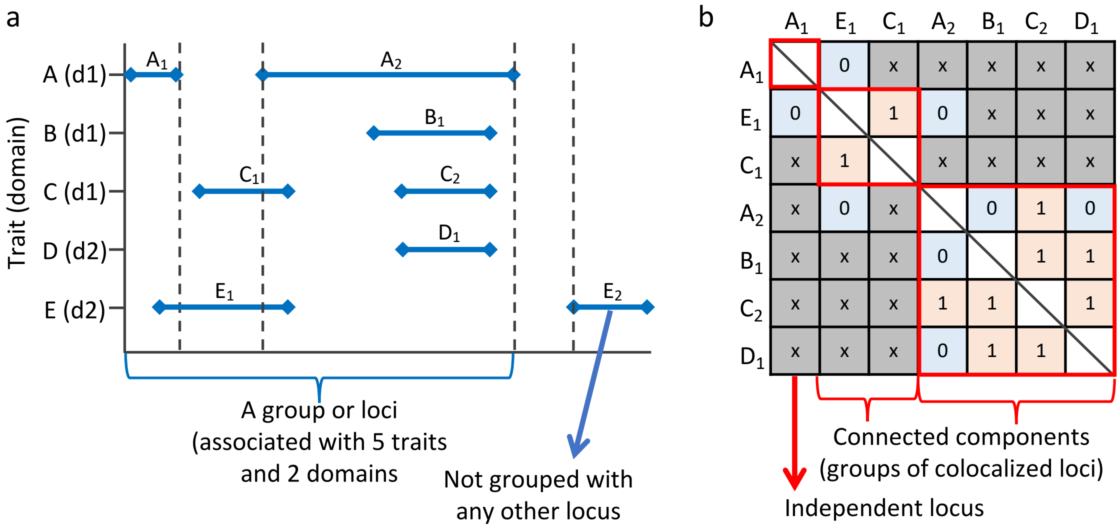

**Fig. 3. Definition of grouped loci based on physical overlap and colocalization. a.** A group of loci defined by physical overlap. X-axis is a chromosome coordinate, Y-axis is trait (domain in parentheses). Horizontal blue lines represent trait-associated loci. Seven loci except $E_1$ are, in this case, grouped into a single consecutive genomic region. This loci group is considered to be associated with 5 traits and 2 domains in the analyses. **b.** A matrix of colocalization pattern across loci within a physically overlapping group defined in **A**. Pair of loci is denoted as 'x' when there is no physical overlapped (colocalization was not performed), 0 when loci are not colocalized and 1 when loci are colocalized (sharing the same causal SNP). Red boxes represent connected component by considering the matrix as a undirected graph and cells with 1 as edges. In this case, there are 3 independent loci grouped based on colocalization and, for each group, the number of traits and domains are counted.

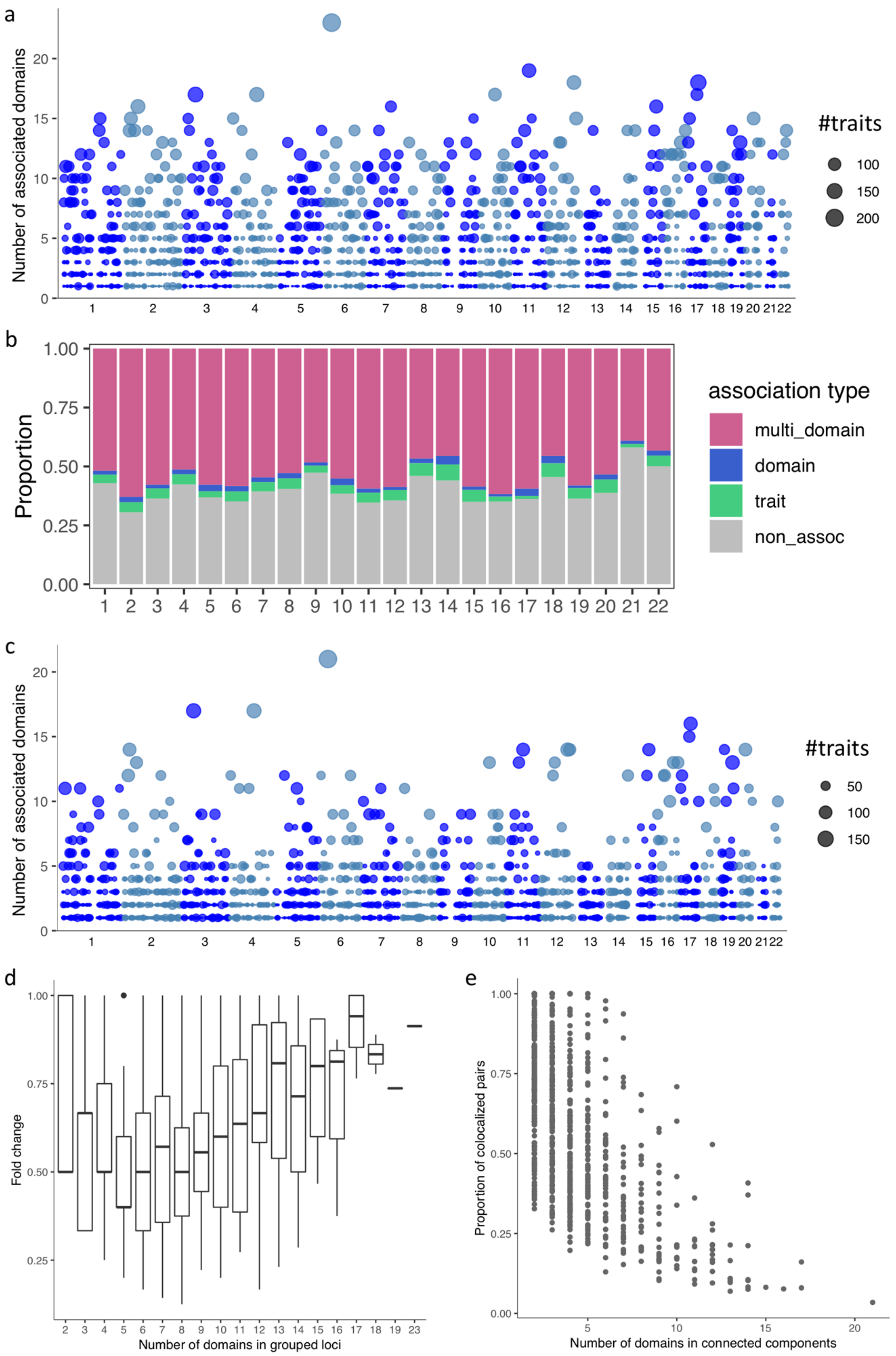

**Fig. 4. Distribution of pleiotropic risk loci and colocalization. a**. Manhattan like plot of pleiotropic trait-associated loci. Each data point represents a group of trait-associated loci

based on physical overlap. Full results are available in **Supplementary Table 4**. **b.** Proportion of the length of trait-associated loci with different association types per chromosome. multi_domain: associated with traits from >1 domain, domain: associated with >1 traits from a single domain, trait: associated with a single trait, non_assoc: not associated with any of 558 traits. **c**. Manhattan like plot for groups of colocalized loci (connected components of colocalized loci). Each data point represent the largest connected component of colocalized loci per a group of physically overlapping loci. **d.** Box plot of the proportional decrease of the number of domains in a connected component of colocalized loci compared to the grouped loci based on physical overlap. Outliers are plotted as dots. For each grouped locus, the largest connected component was selected. The x-axis is the number of domains in grouped loci based on physical overlap. **d.** The proportion of colocalized locus pairs per connected component of colocalized loci.

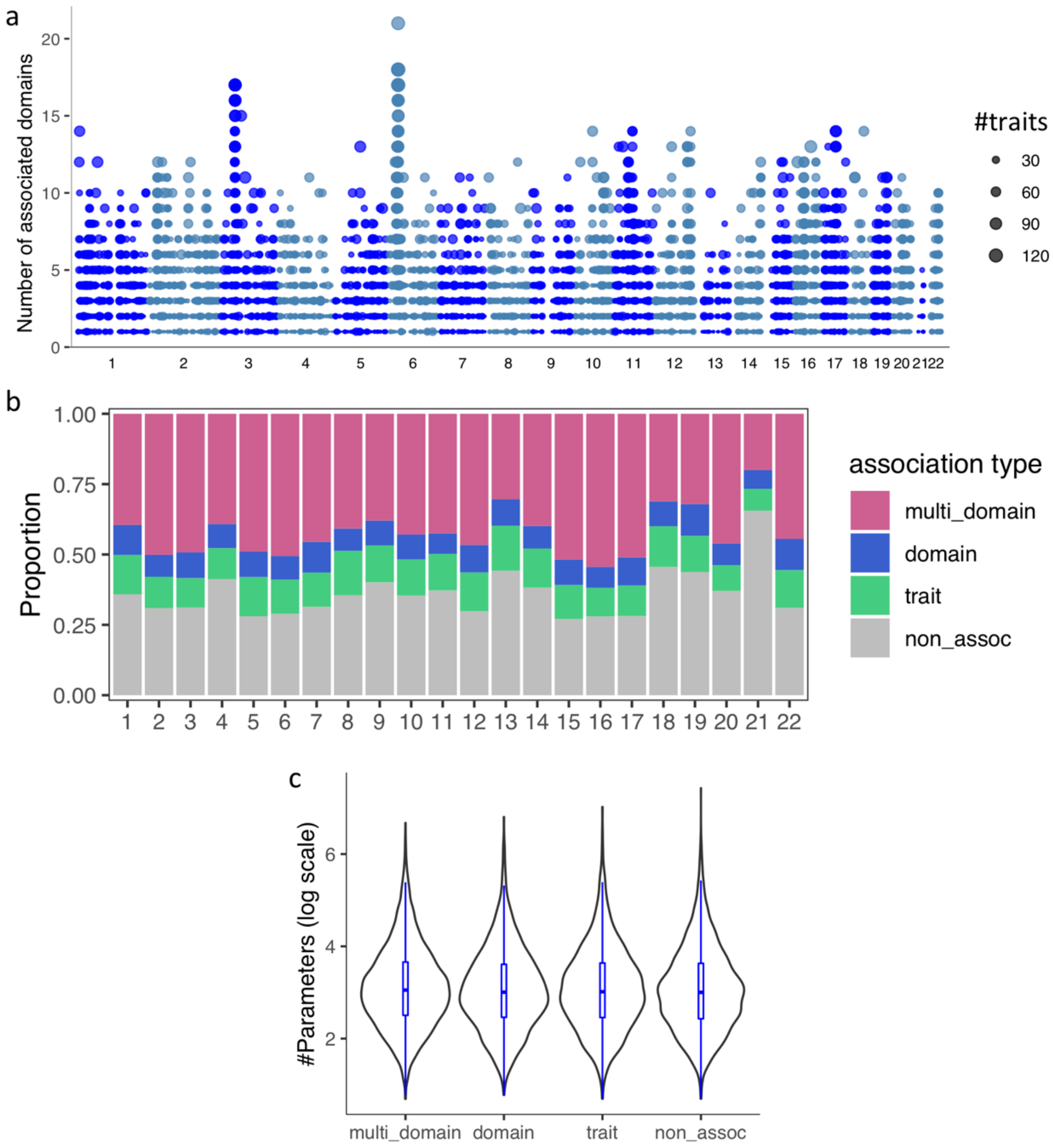

**Fig. 5. Distribution of pleiotropic genes and features of genes and gene sets. a.** Manhattan like plot of pleiotropic genes, every dot represents a single gene. Full results are available in **Supplementary Table** 7. **b.** Proportion of genes with different association types per chromosome. multi_domain: associated with traits from >1 domain, domain: associated with >1 traits from a single domain, trait: associated with a single trait, non_assoc: not associated with any of 558 traits. **c.** Distribution of the number of parameters used in the MAGMA gene analysis with different association types.

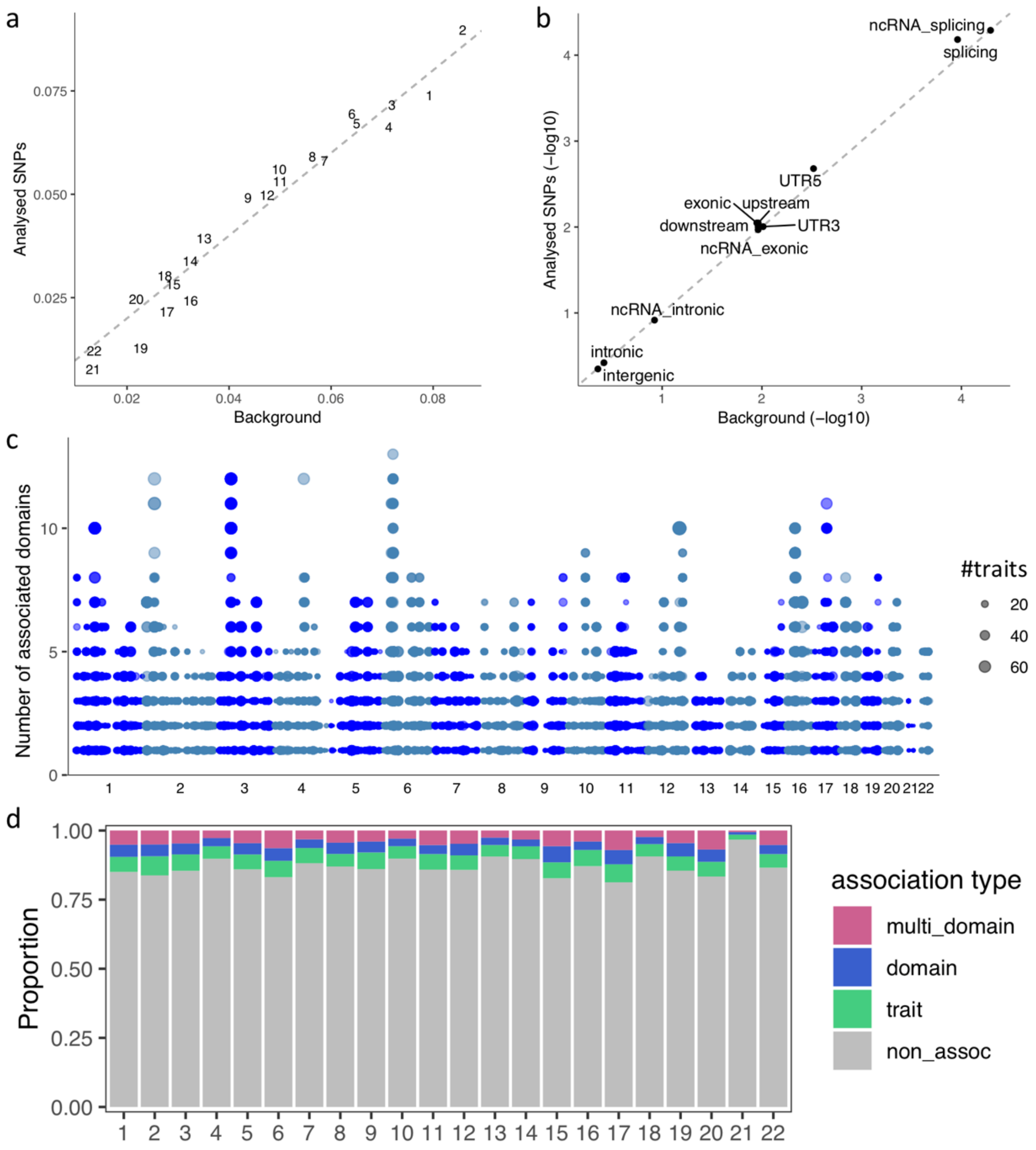

**Fig. 6. Distribution of pleiotropic SNPs. a.** Distribution of all SNPs in the genome (x-axis) and analyzed SNPs (y-axis) across chromosomes. **b.** Distribution of functional consequences of all SNPs in the genome (x-axis) and analyzed SNPs (y-axis). **c.** Manhattan like plot of pleiotropic SNPs. Each data point represent a single SNP and only SNPs associated with >1 trait are displayed. Full results are available in **Supplementary Table 12**. **d.** Proportion of SNPs with different association types per chromosome. multi_domain: associated with traits from >1 domain, domain: associated with >1 traits from a single domain, trait: associated with a single trait, non_assoc: not associated with any of 558 traits.

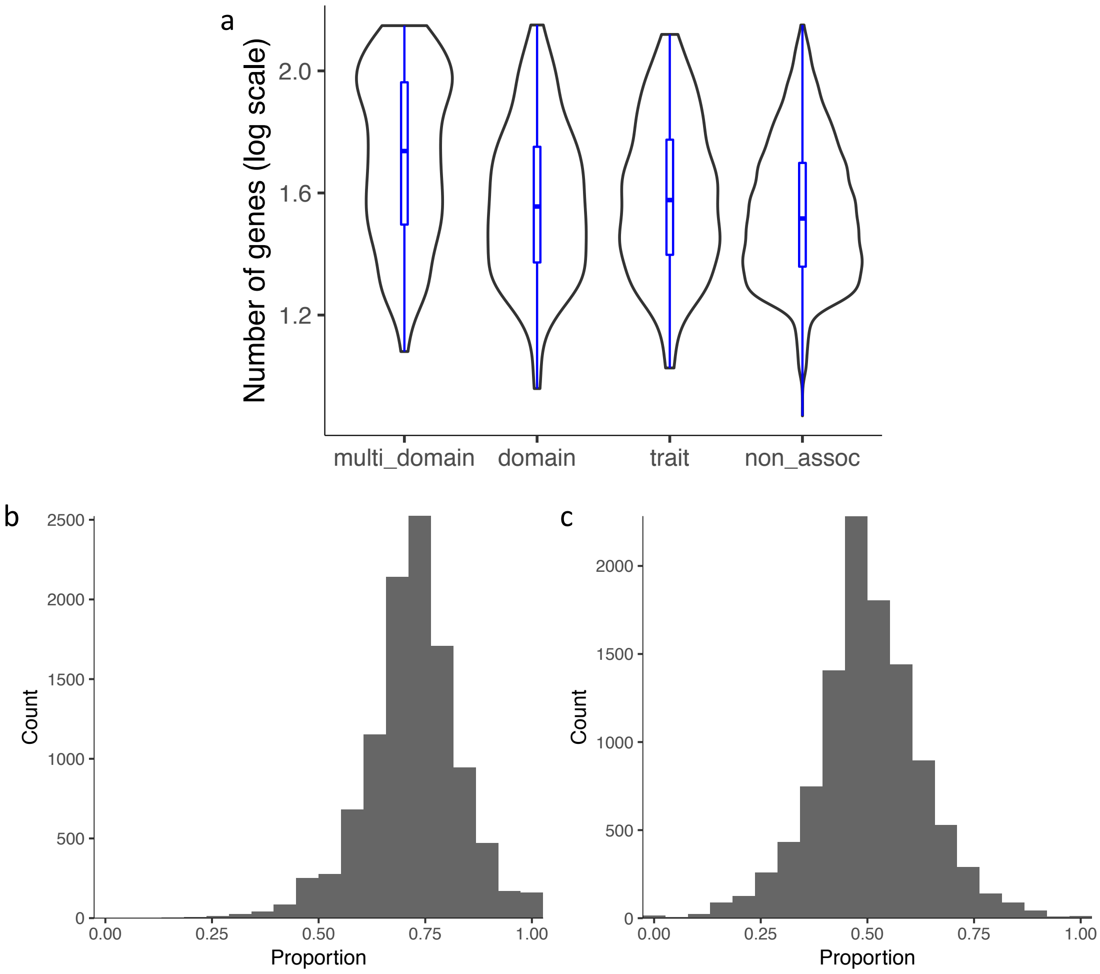

**Fig. 7. Distribution of pleiotropic gene-sets. a.** Distribution of the number of genes in the gene-sets with different association types. multi_domain: associated with traits from >1 domain, domain: associated with >1 traits from a single domain, trait: associated with a single trait, non_assoc: not associated with any of 558 traits. **b.** Histogram of the proportion of genes associated with at least one of the 558 traits in a gene-set. **c.** Histogram of the proportion of genes associated with traits from more than one domain in a gene-set.

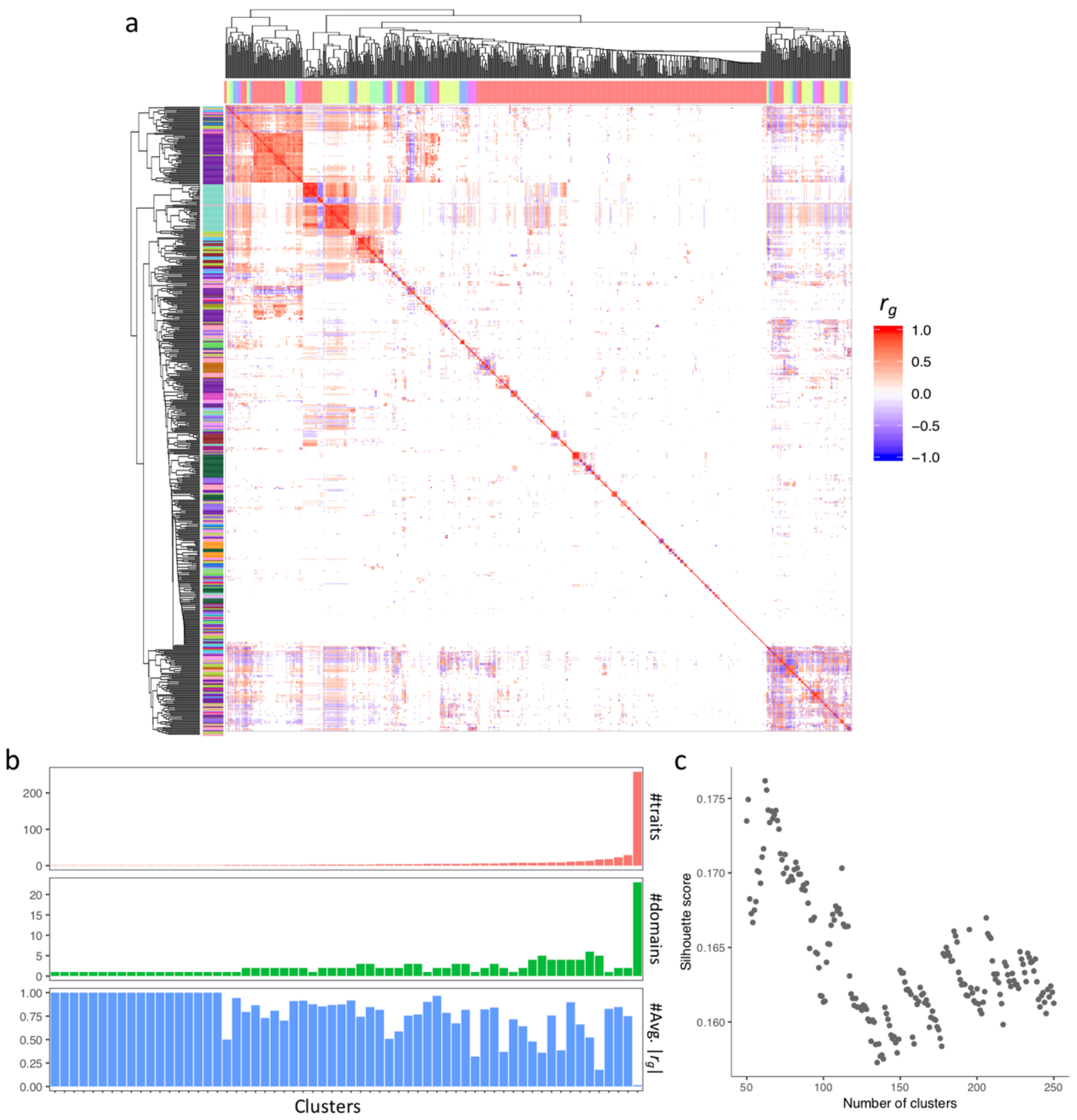

**Fig. 8. Features of trait clusters based on genetic correlation. a.** Heatmap of the genetic correlation ($r_g$) matrix. Both columns and rows are hierarchically clustered. The color bar at the top (x-axis) represents the clusters (consecutive traits with the same color are in the same cluster) and the color bar on the left (y-axis) represent domains of the traits. **b.** Number of traits, domains and average genetic correlation per cluster. **c.** Silhouette score with different number of clusters.

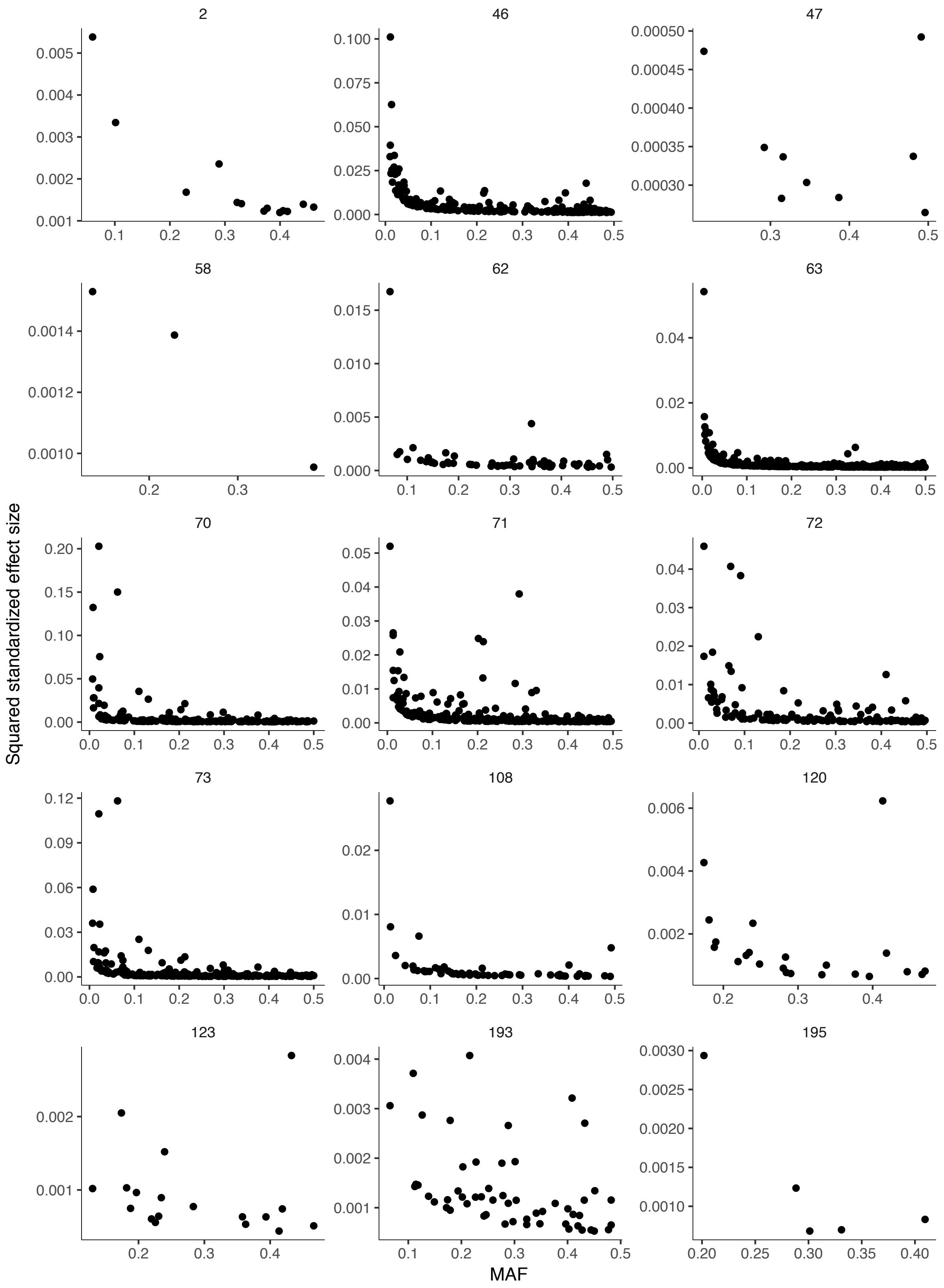

2
46
47
58
62
63
70
71
72
73
108
120
123
193
195
Squared standardized effect size
MAF

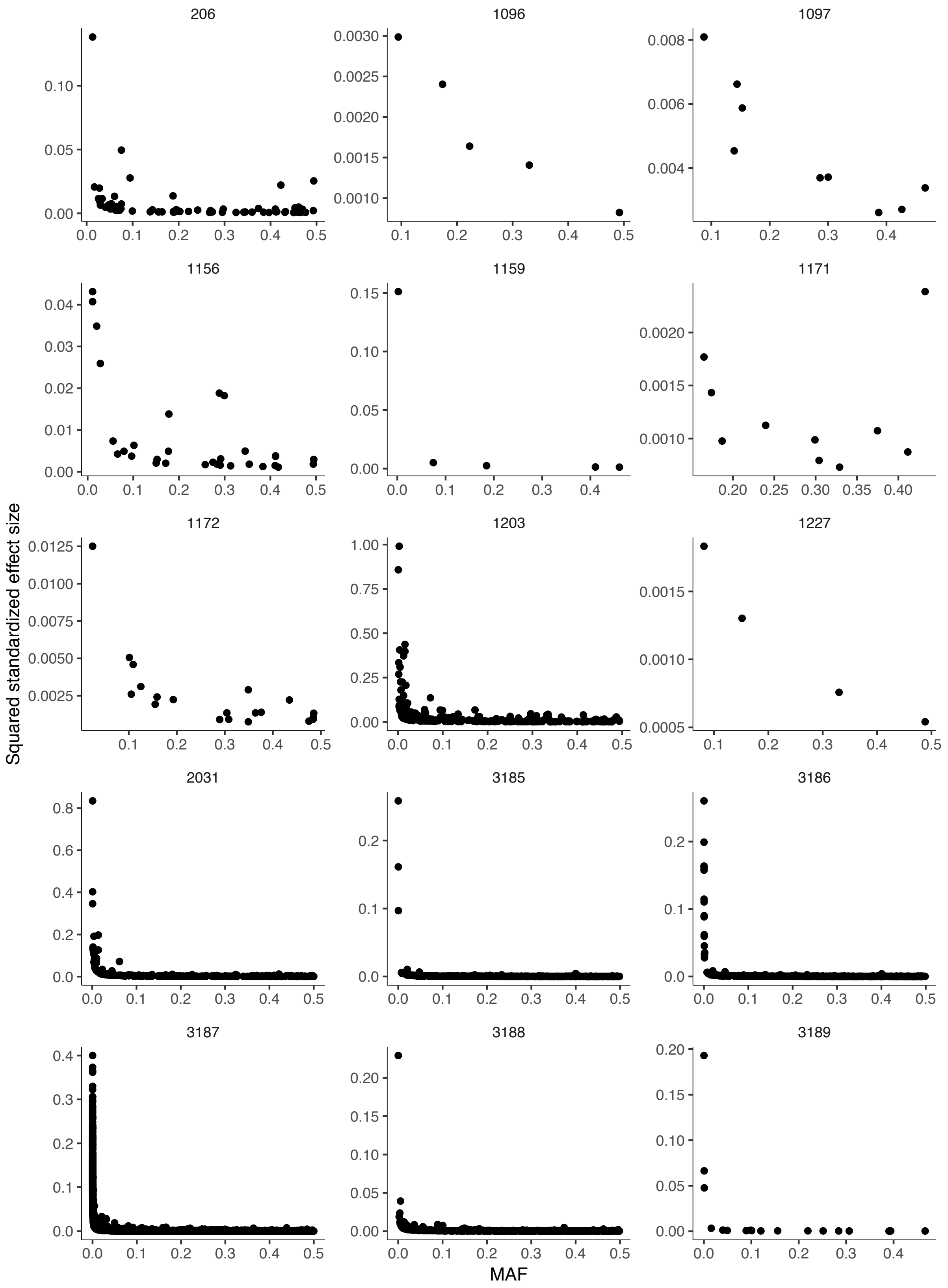
206
1096
1097
1156
1159
1171
1172
1203
1227
2031
3185
3186
3187
3188
3189
Squared standardized effect size
MAF

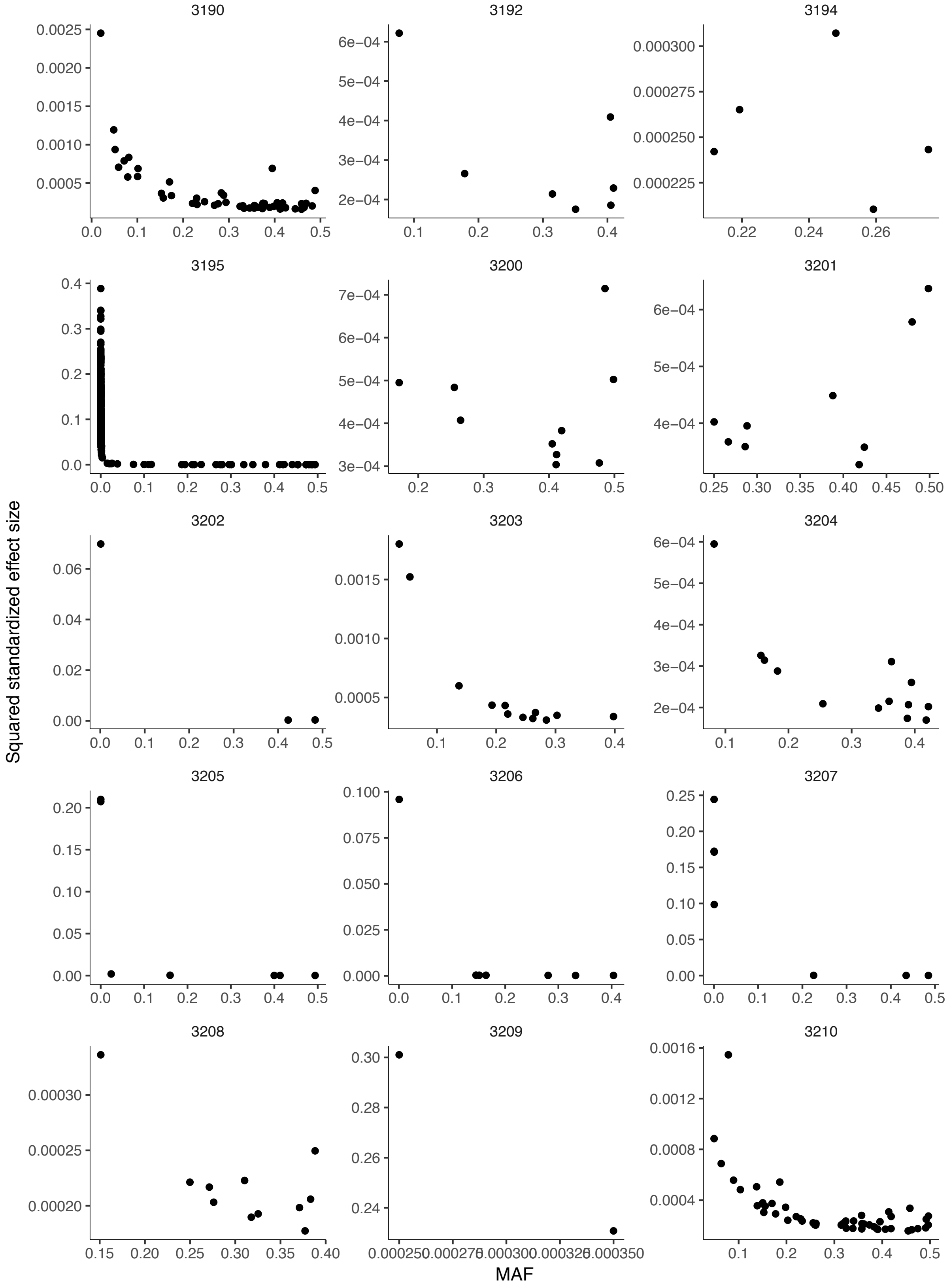
3190
3192
3194
3195
Squared standardized effect size
MAF

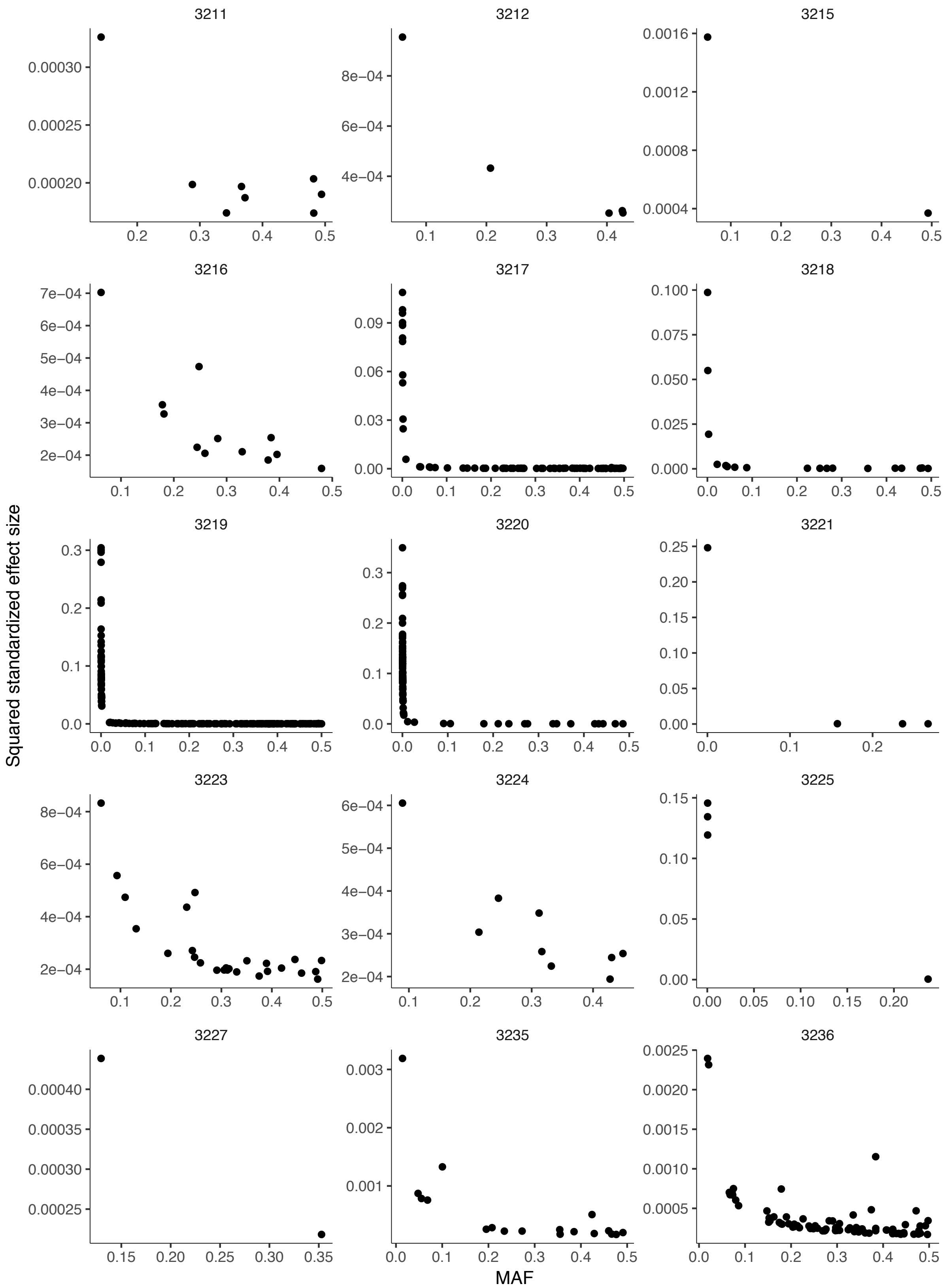
Squared standardized effect size
MAF
3211
3212
3215
3216
3217
3218
3219
3220
3221
3223
3224
3225
3227
3235
3236

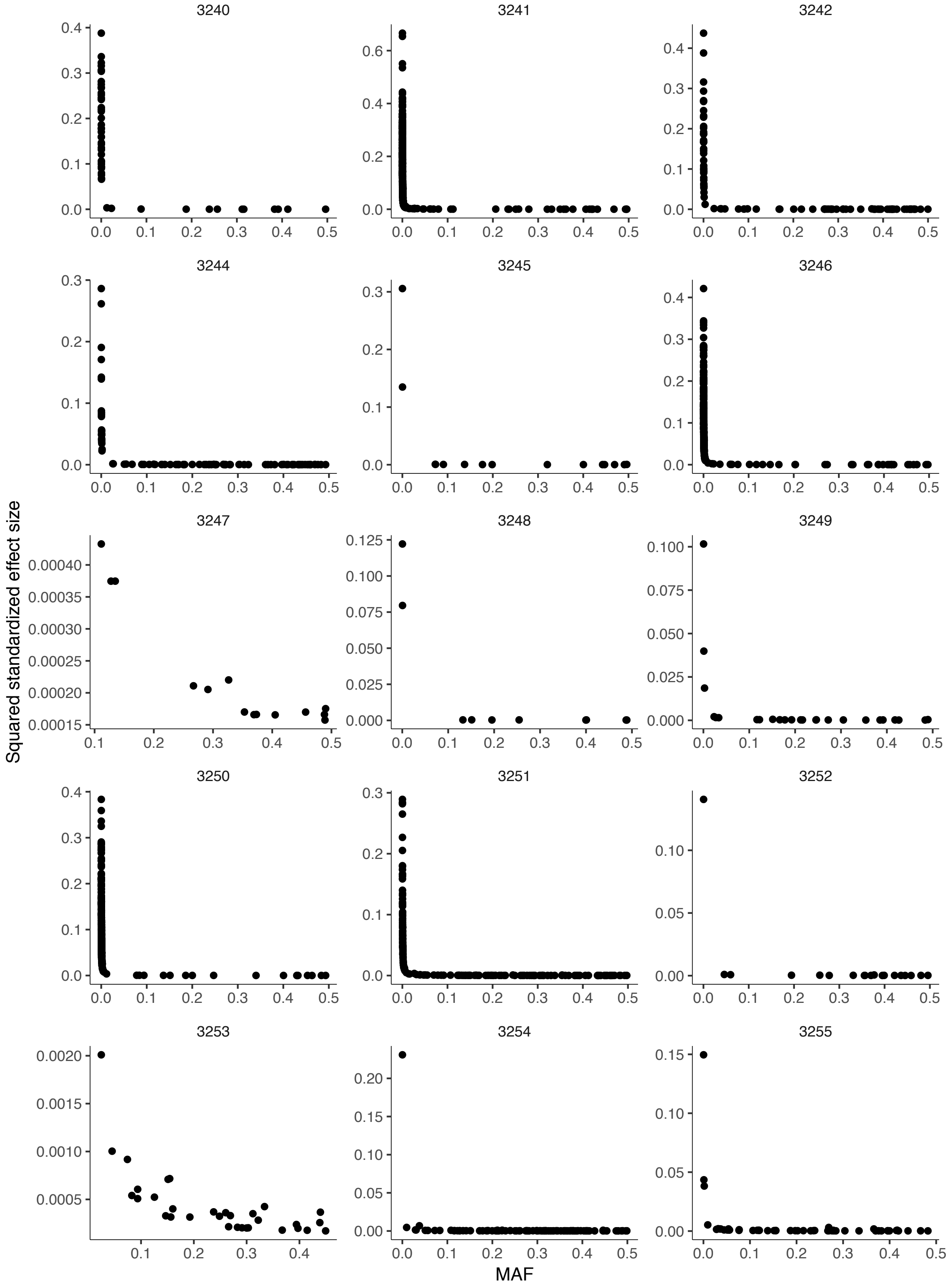
3240
3241
3242
Squared standardized effect size
MAF

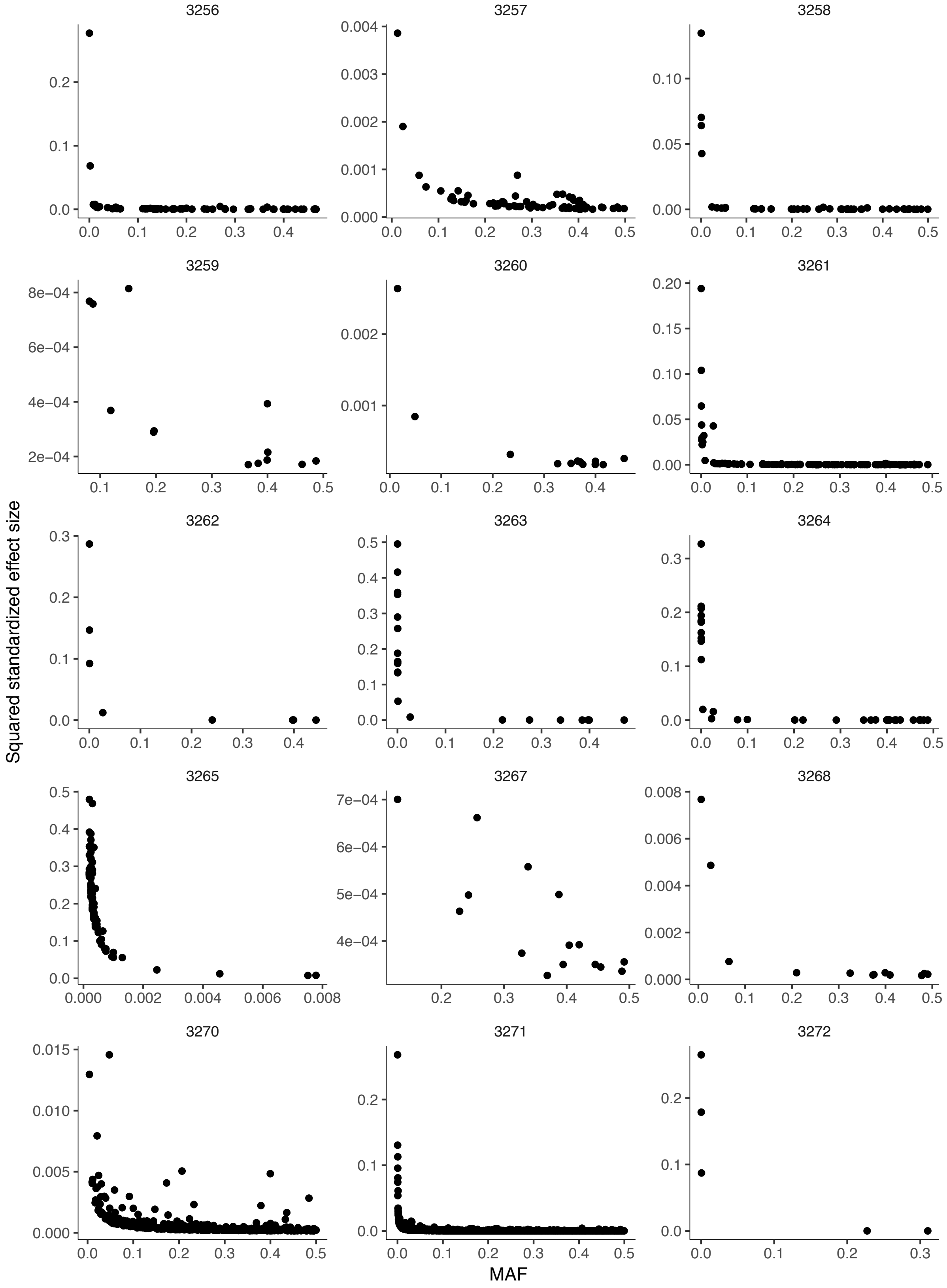
3267
3268
3270
3271
3272
Squared standardized effect size
MAF

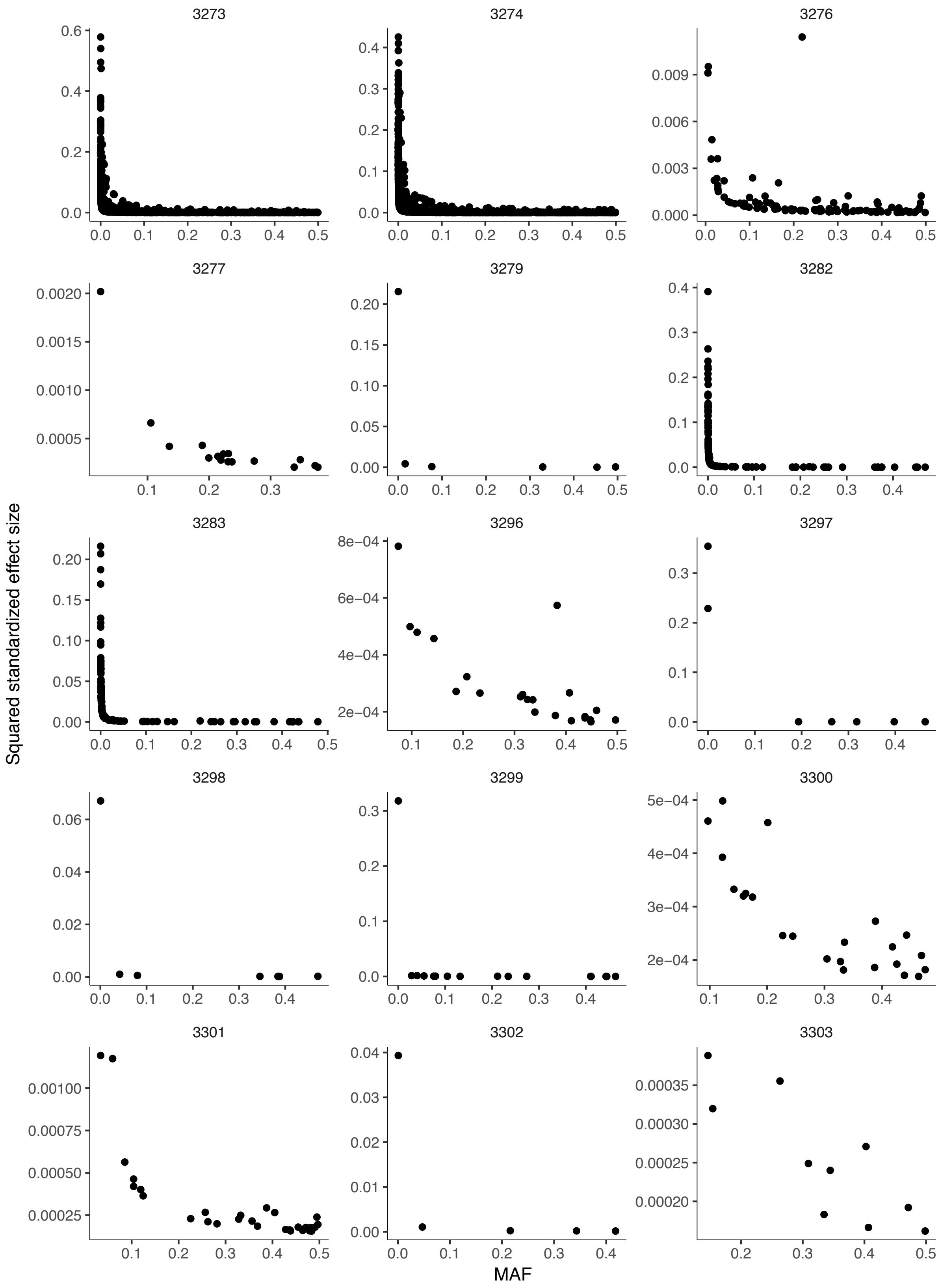
3273
3274
3276
3277
3279
3282
3283
3296
3297
3298
3299
3300
3301
3302
3303
Squared standardized effect size
MAF

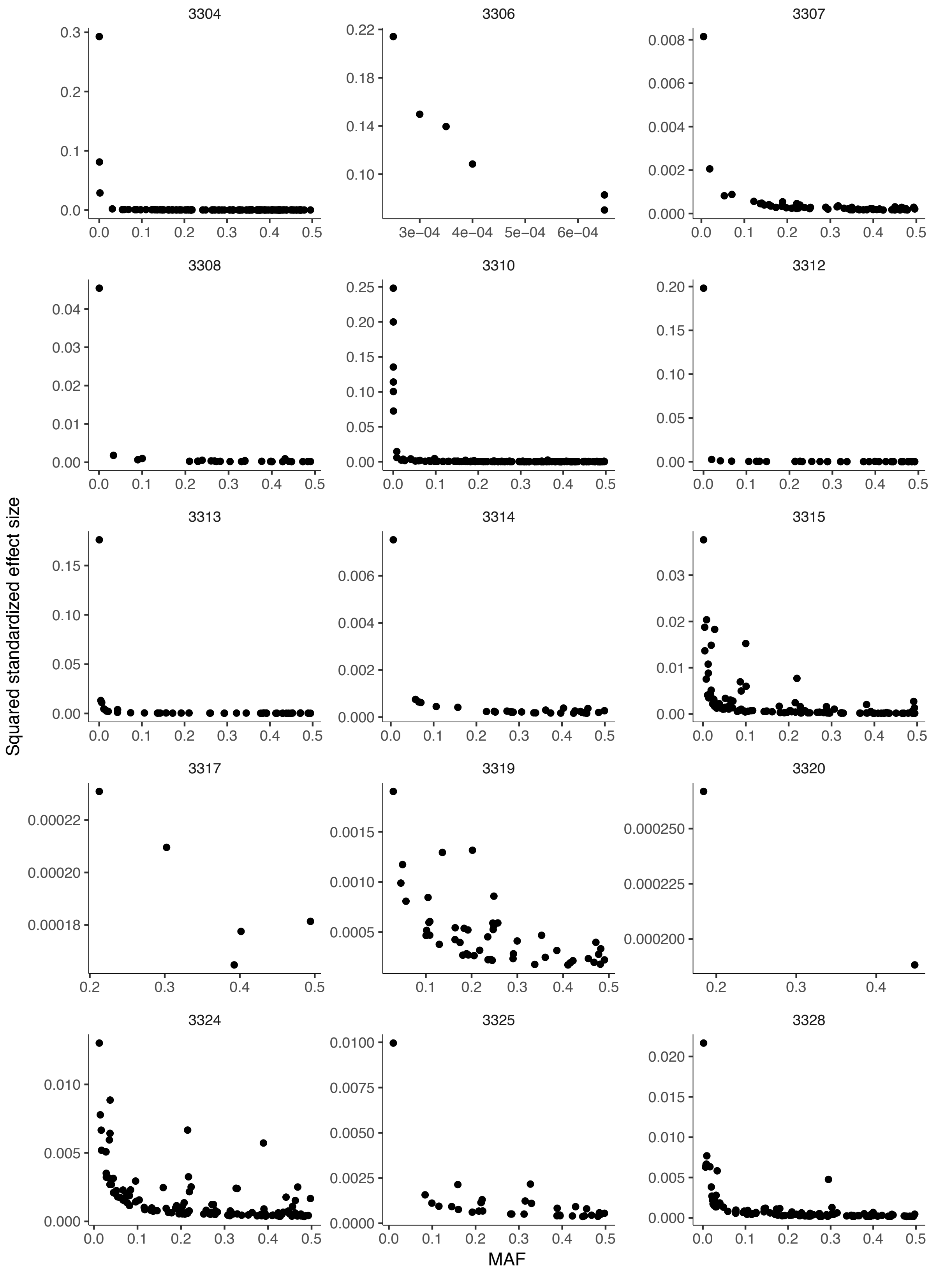
3304
3306
3307
3308
3310
3312
3313
3314
3315
3317
3319
3320
3324
3325
3328
Squared standardized effect size
MAF

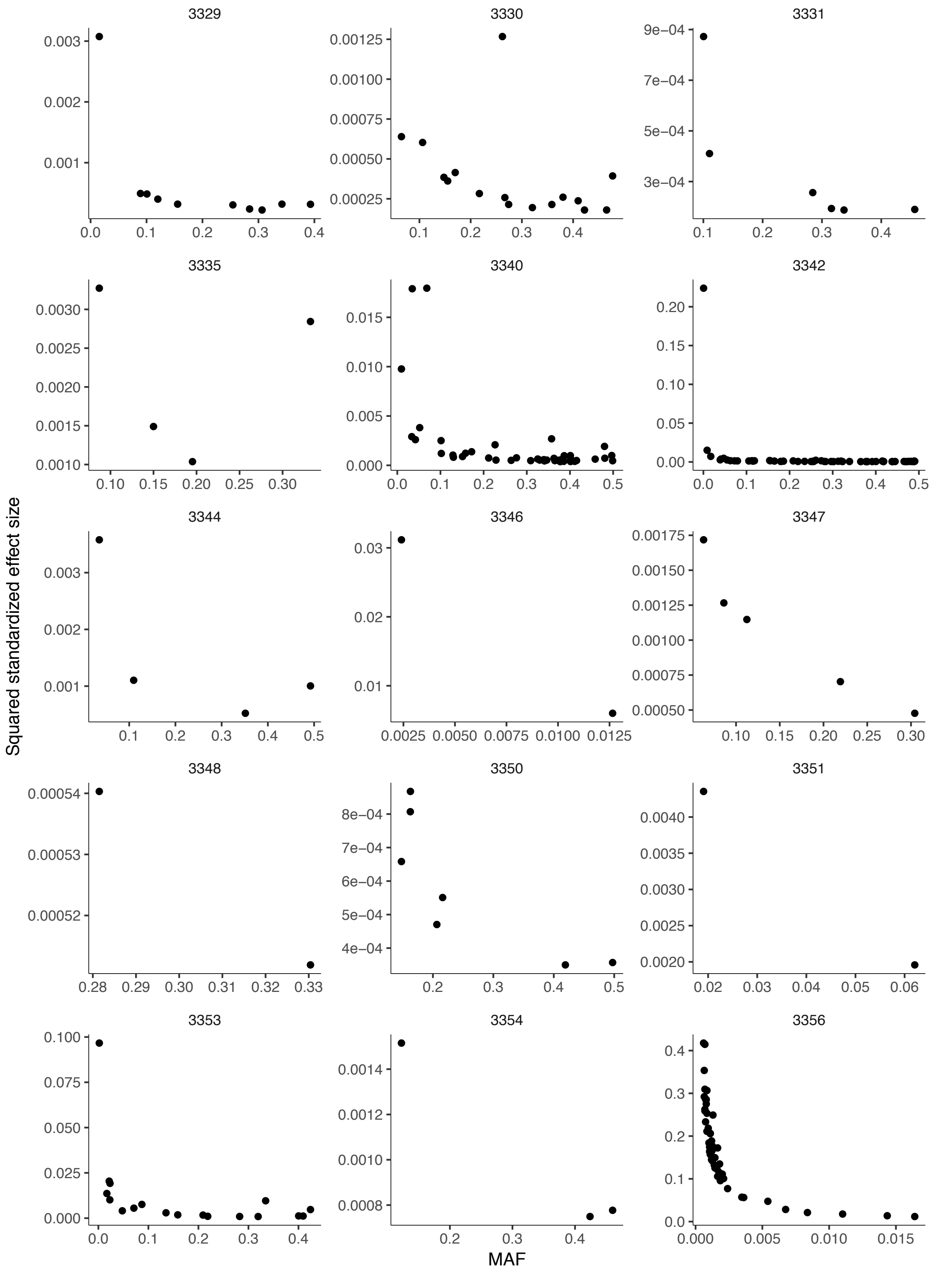
3329
3330
3331
3335
3340
3342
3344
3346
3347
3348
3350
3351
3353
3354
3356
Squared standardized effect size
MAF

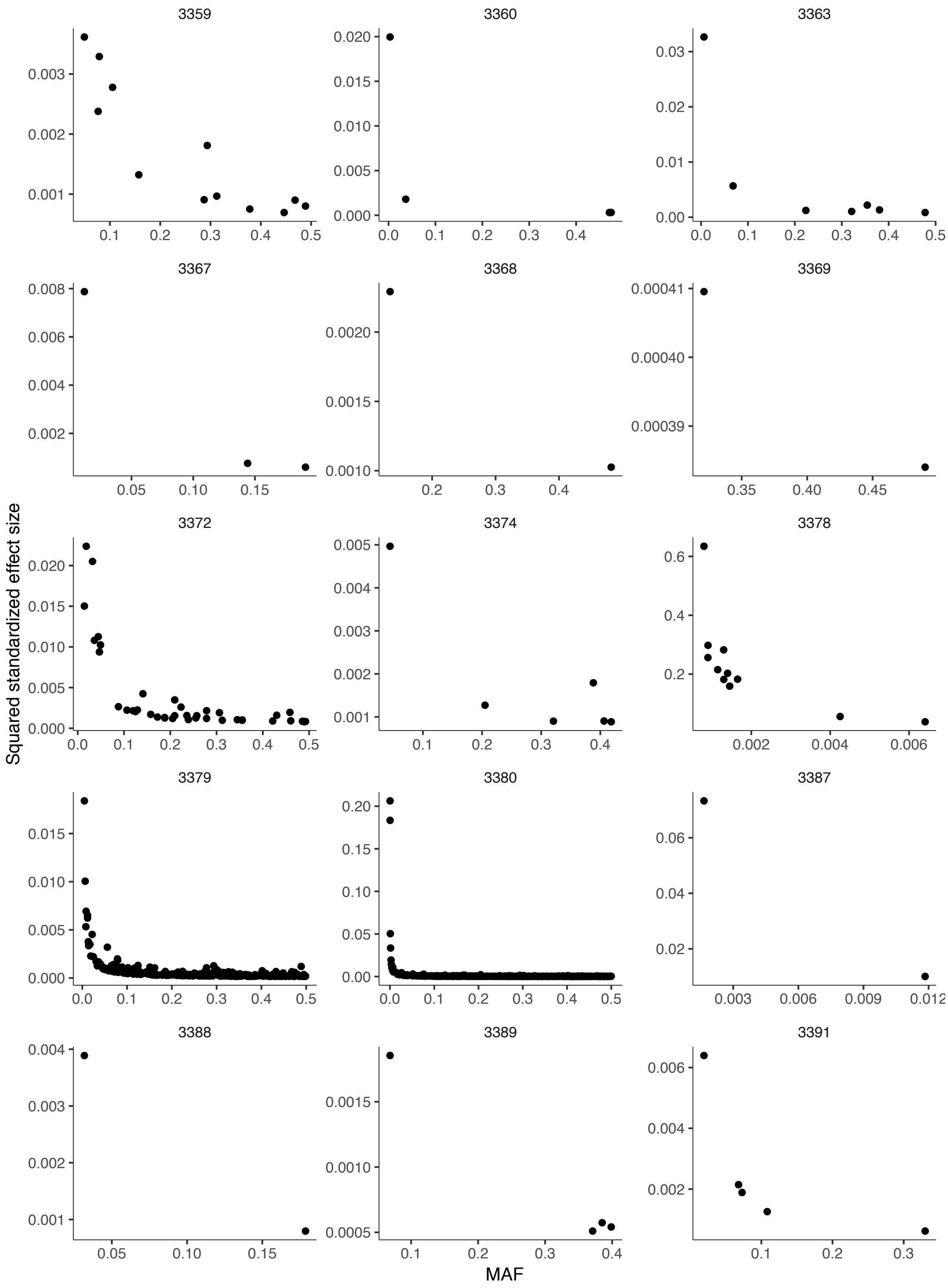

3359
3360
3363
3367
3368
3369
3372
3374
3378
3379
3380
3387
3388
3389
3391
Squared standardized effect size
MAF

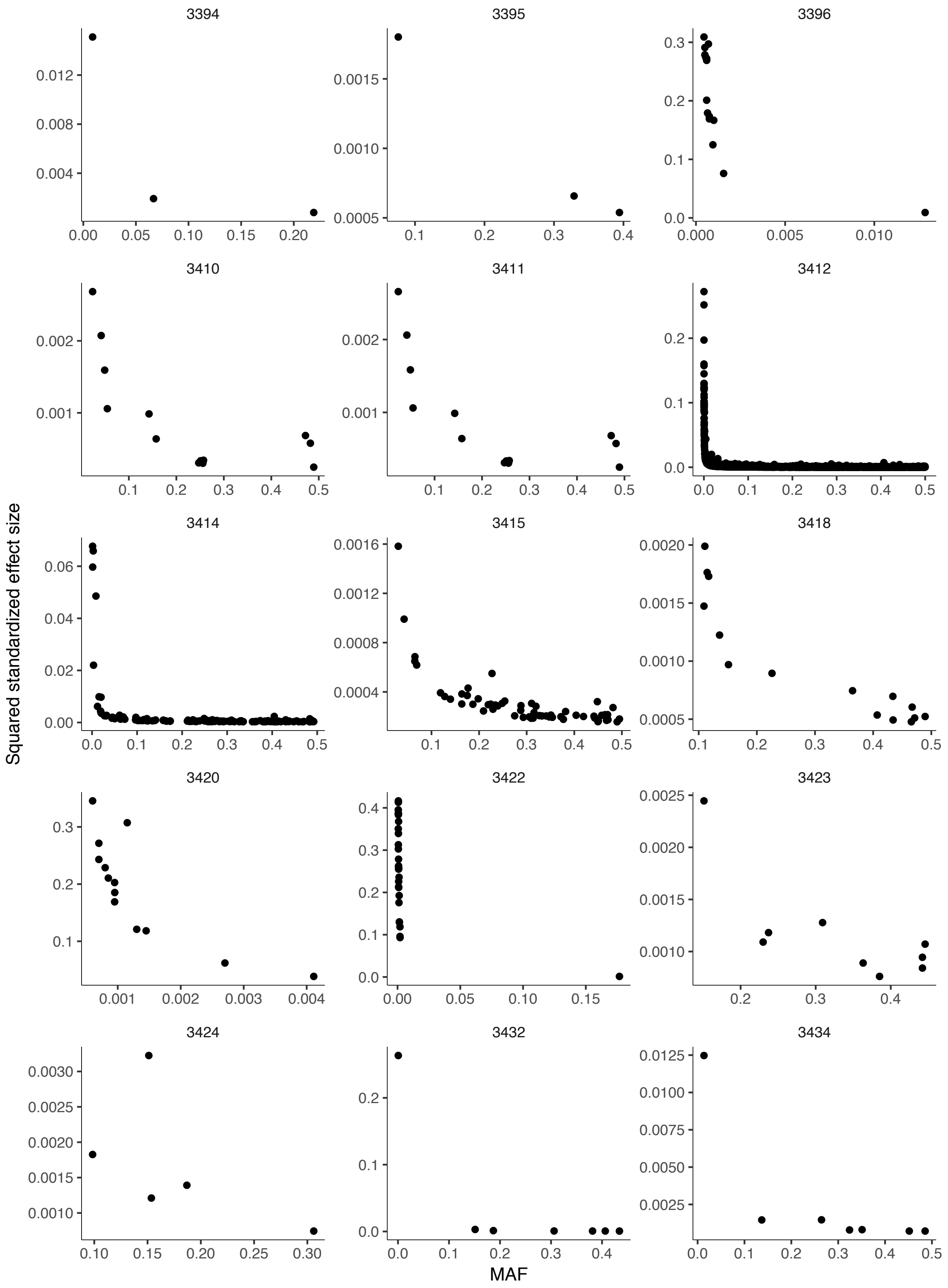
3394
3395
3396
3410
3411
3412
3414
3415
3418
3420
3422
3423
3424
3432
3434
Squared standardized effect size
MAF

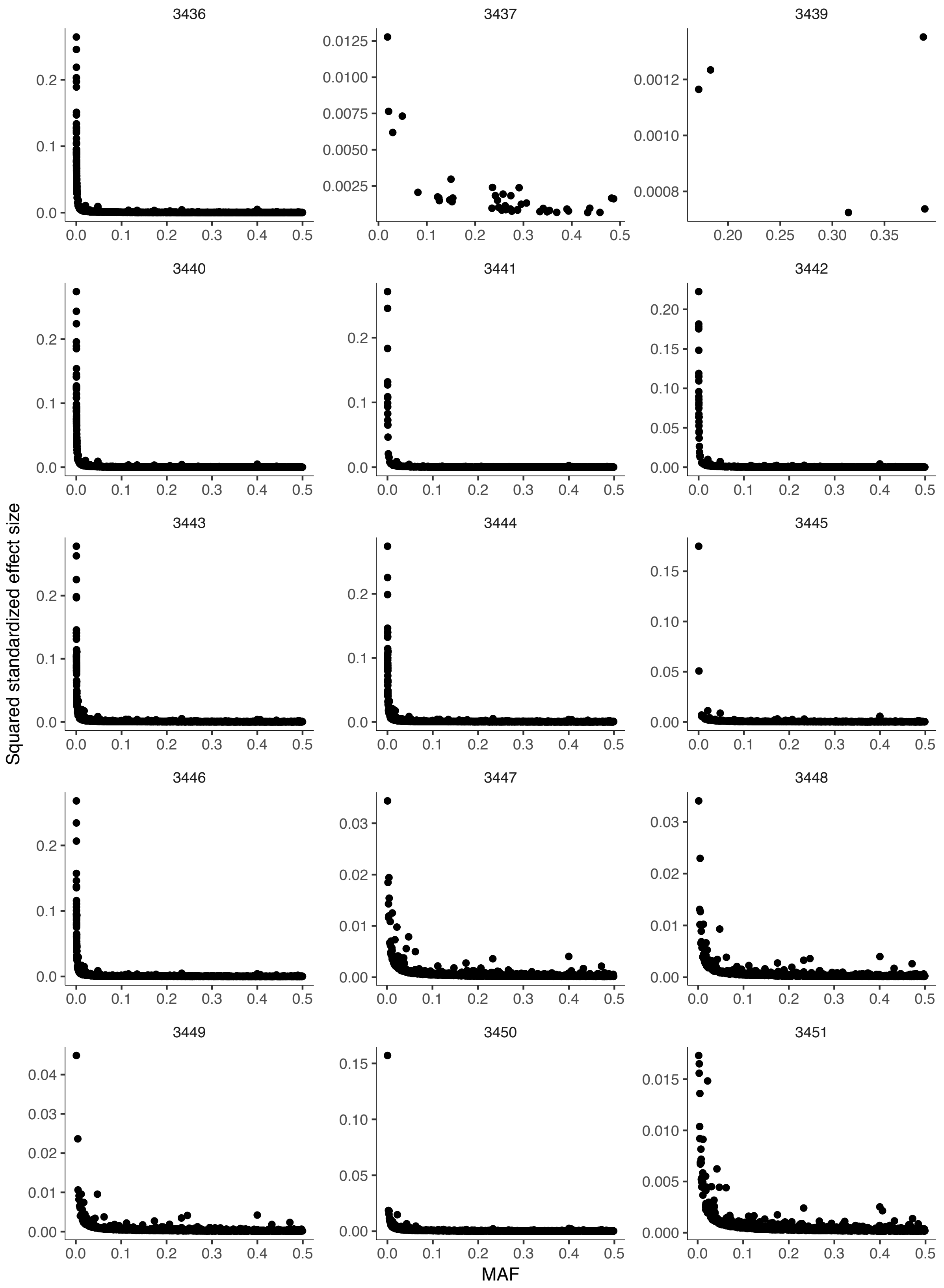

3436
3437
Squared standardized effect size
MAF

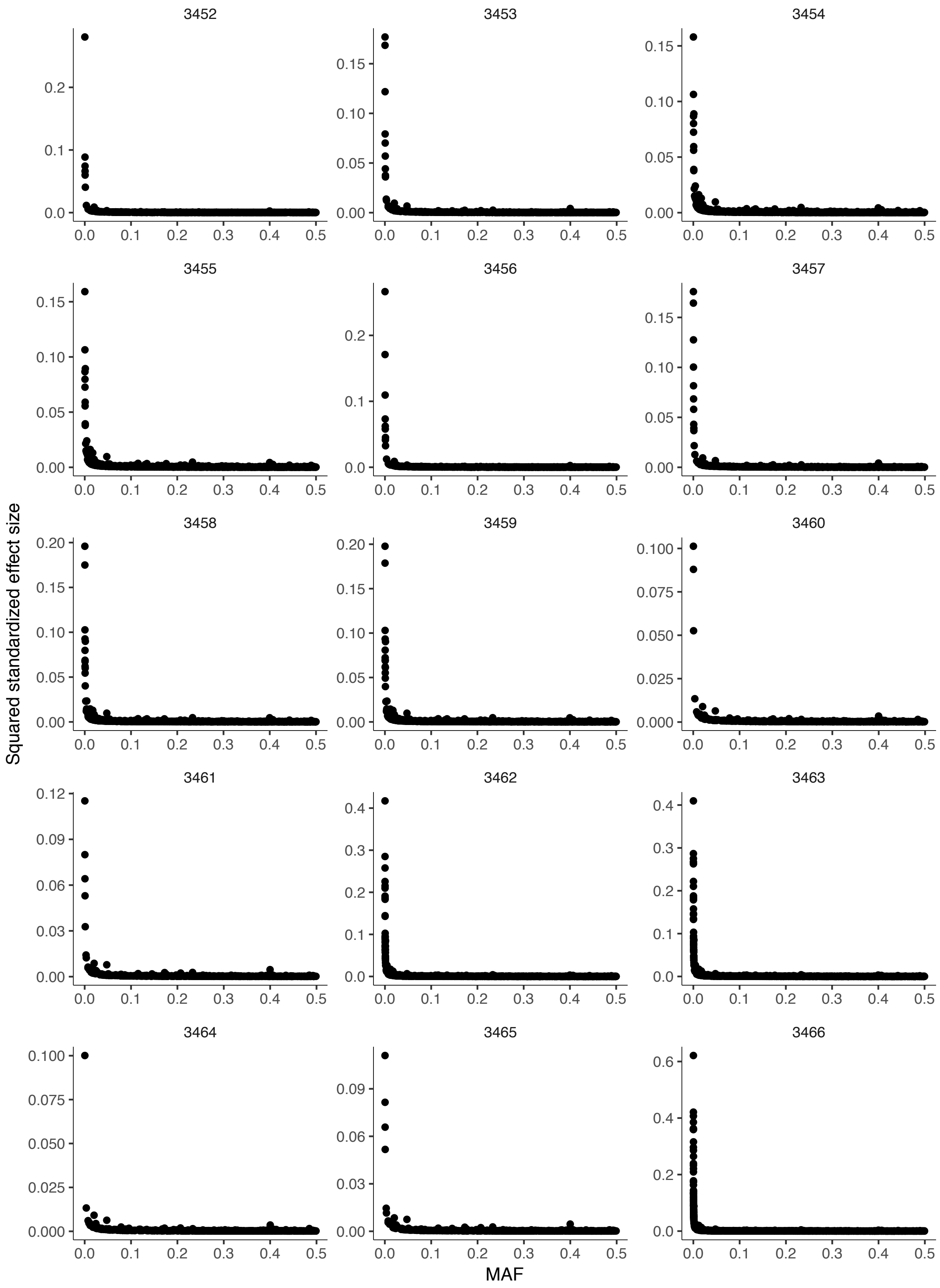
Squared standardized effect size
MAF

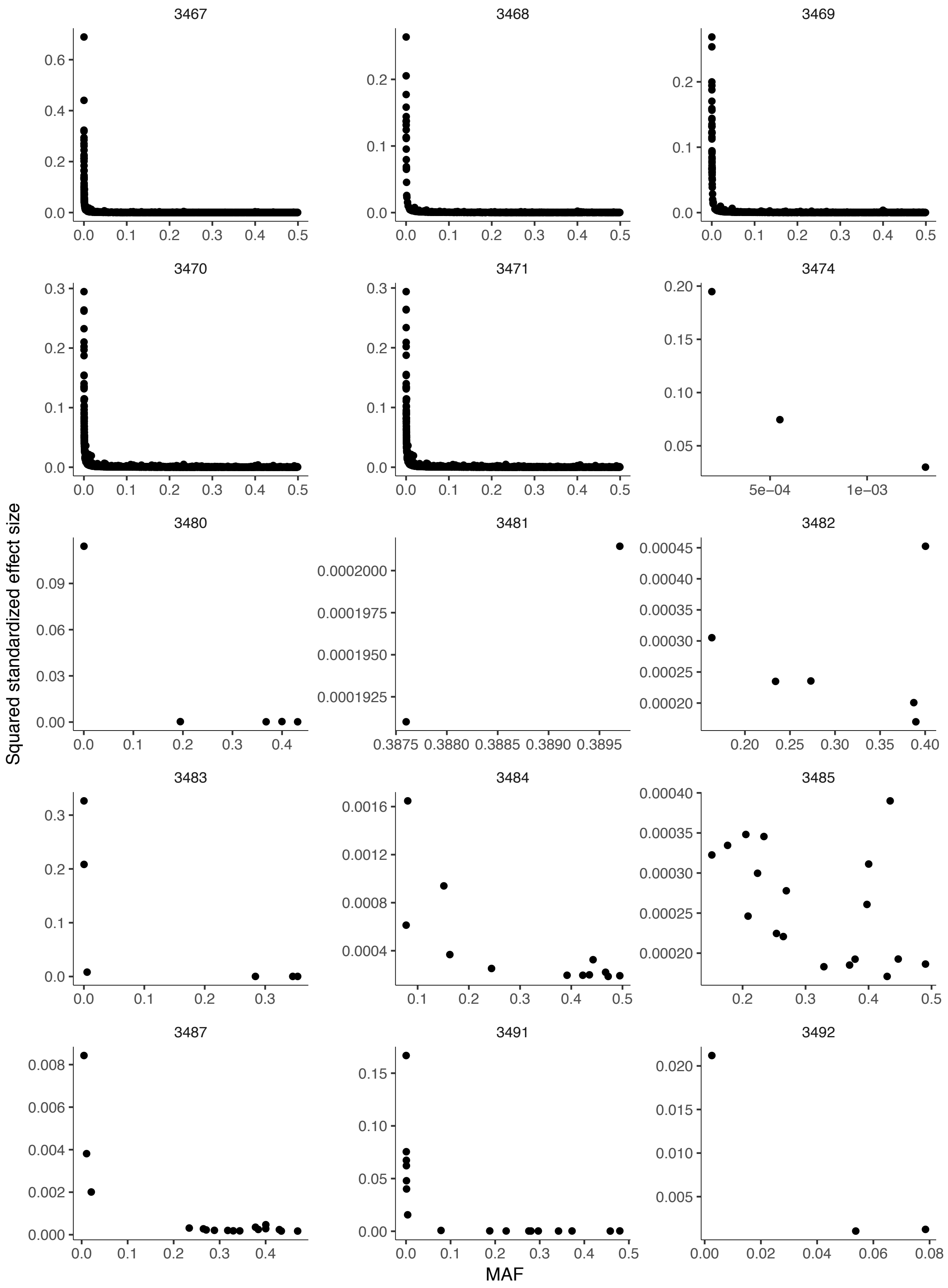

3467
3468
3469
3470
3471
3474
3480
3481
3482
3483
3484
3485
3487
3491
3492
Squared standardized effect size
MAF

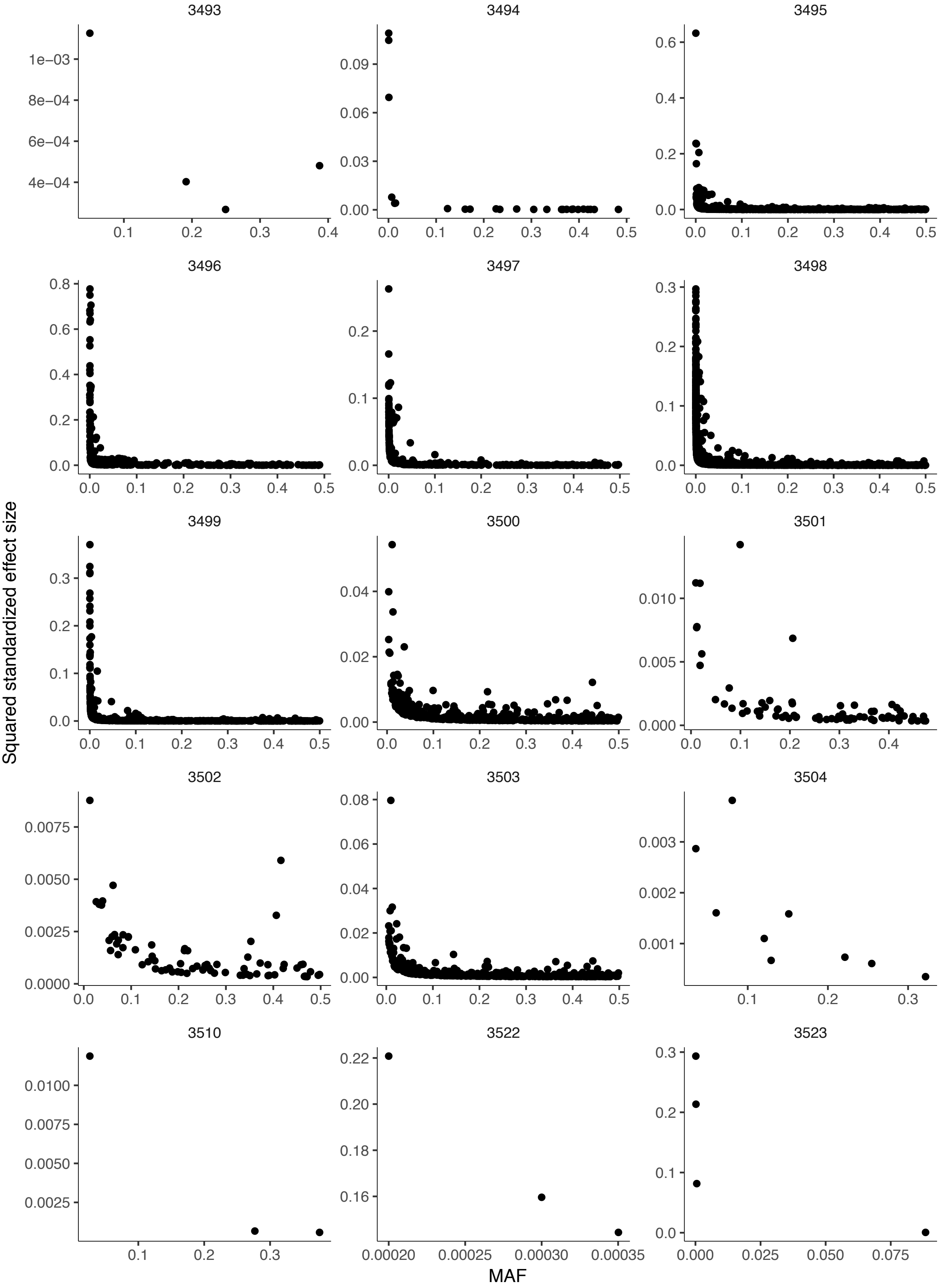
3510
3522
3523
Squared standardized effect size
MAF

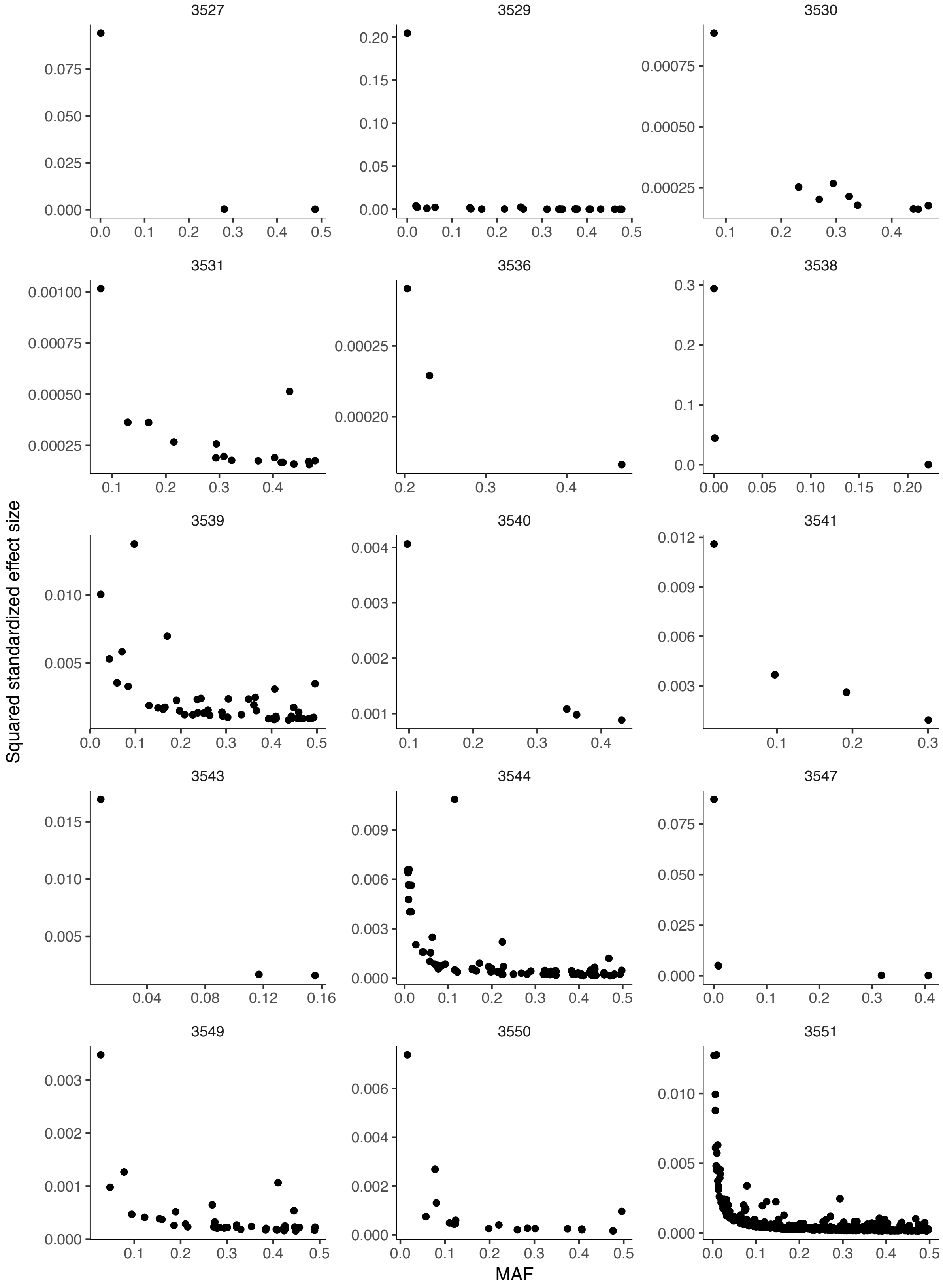
3527
3529
3530
3531
3536
3538
3539
3540
3541
3543
3544
3547
3549
3550
3551
Squared standardized effect size
MAF

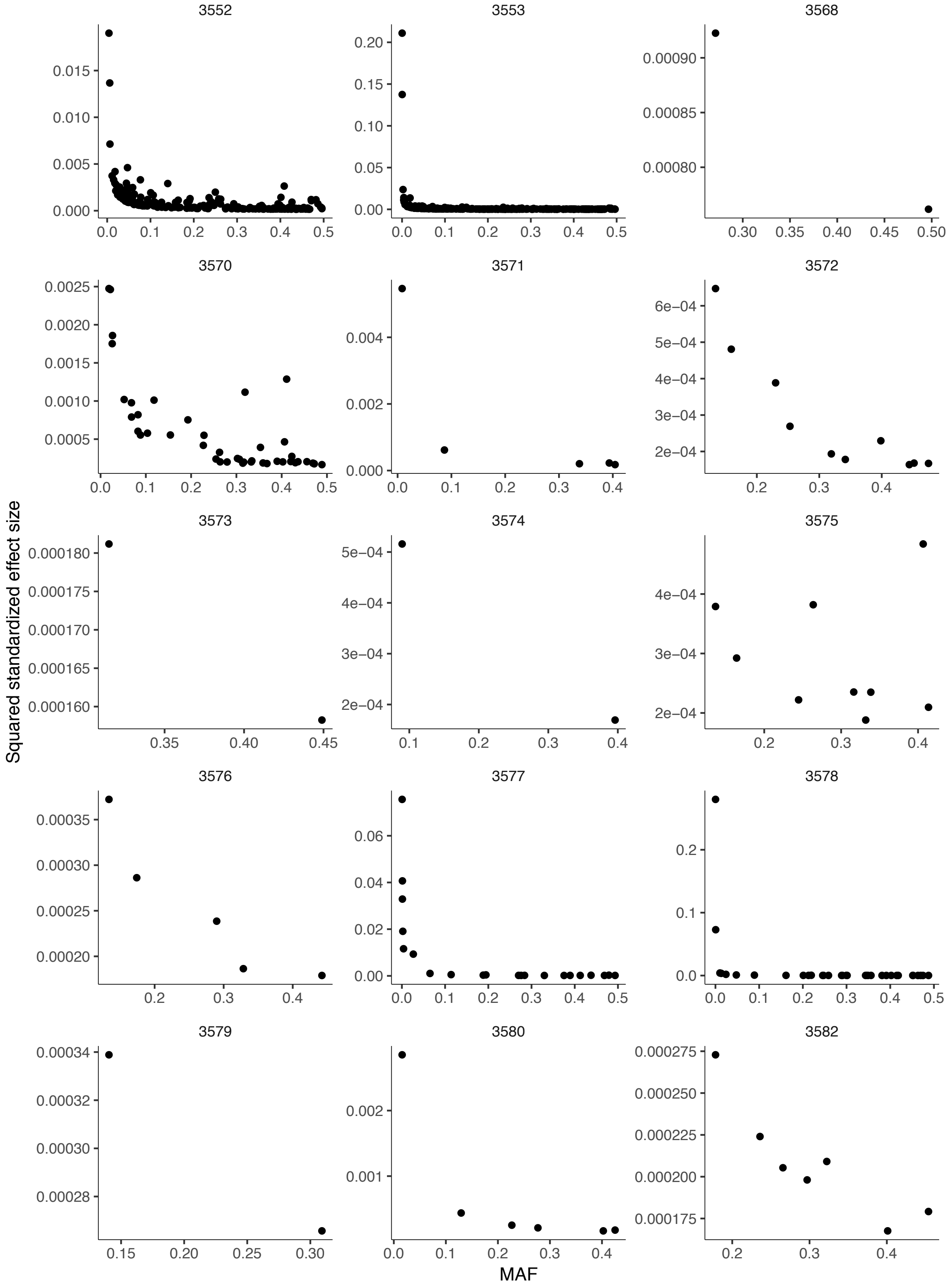
3552
3553
3568
3582
Squared standardized effect size
MAF

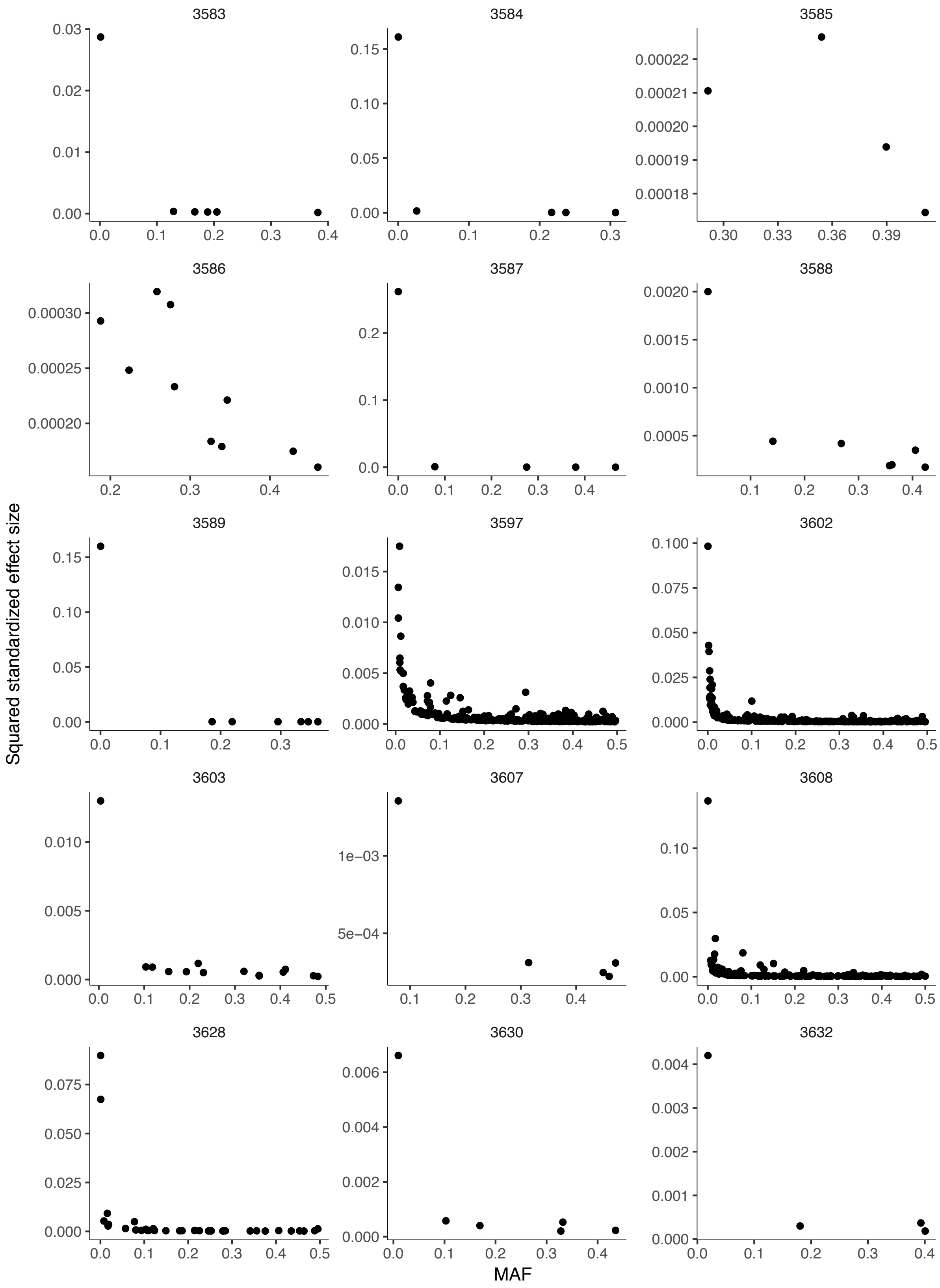
3583
3584
3585
3586
3587
3588
3589
3597
3602
3603
3607
3608
3628
3630
3632
Squared standardized effect size
MAF

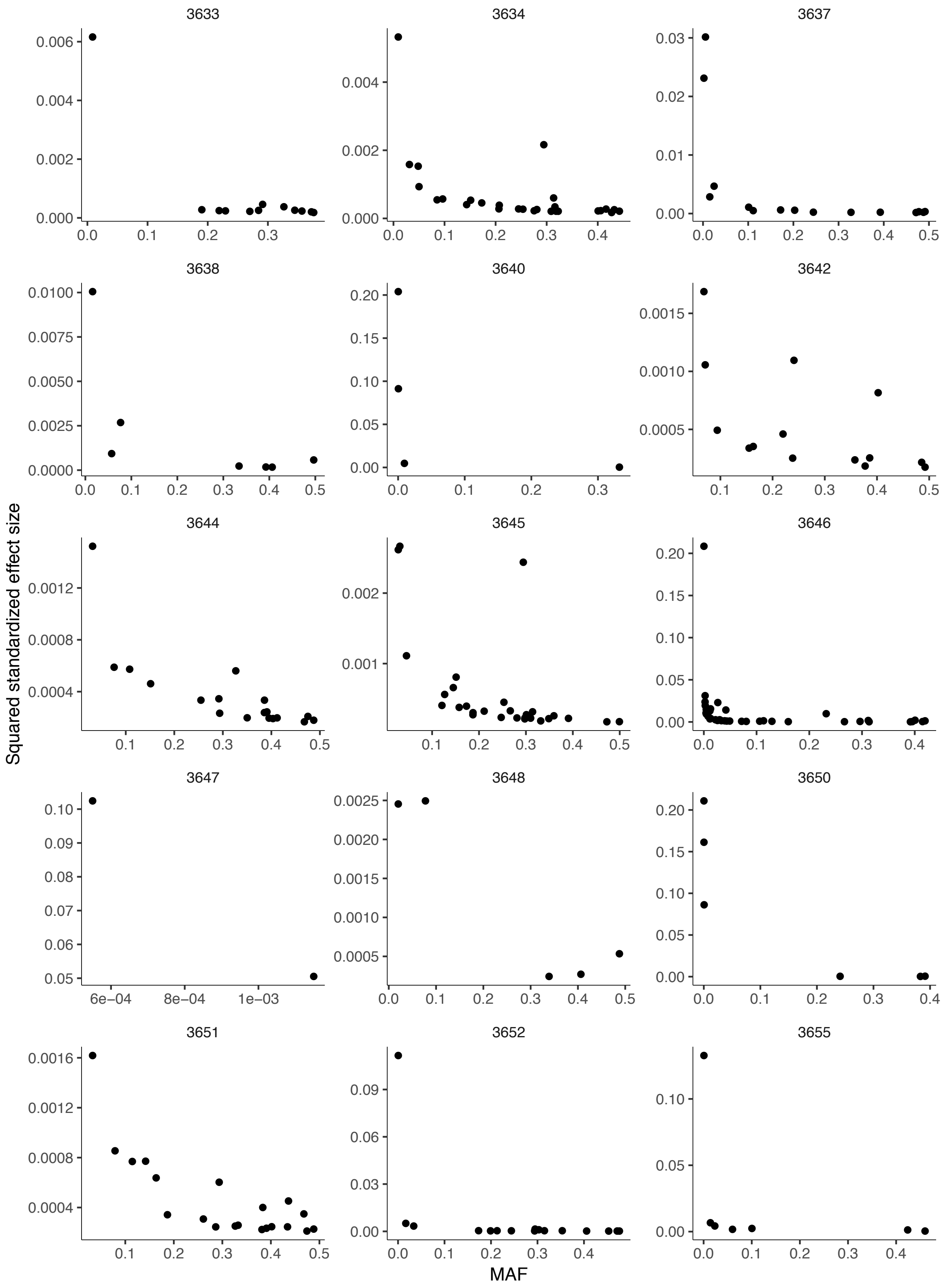

3633
3634
3637
3638
3640
3642
3644
3645
3646
3647
3648
3650
3651
3652
3655
Squared standardized effect size
MAF

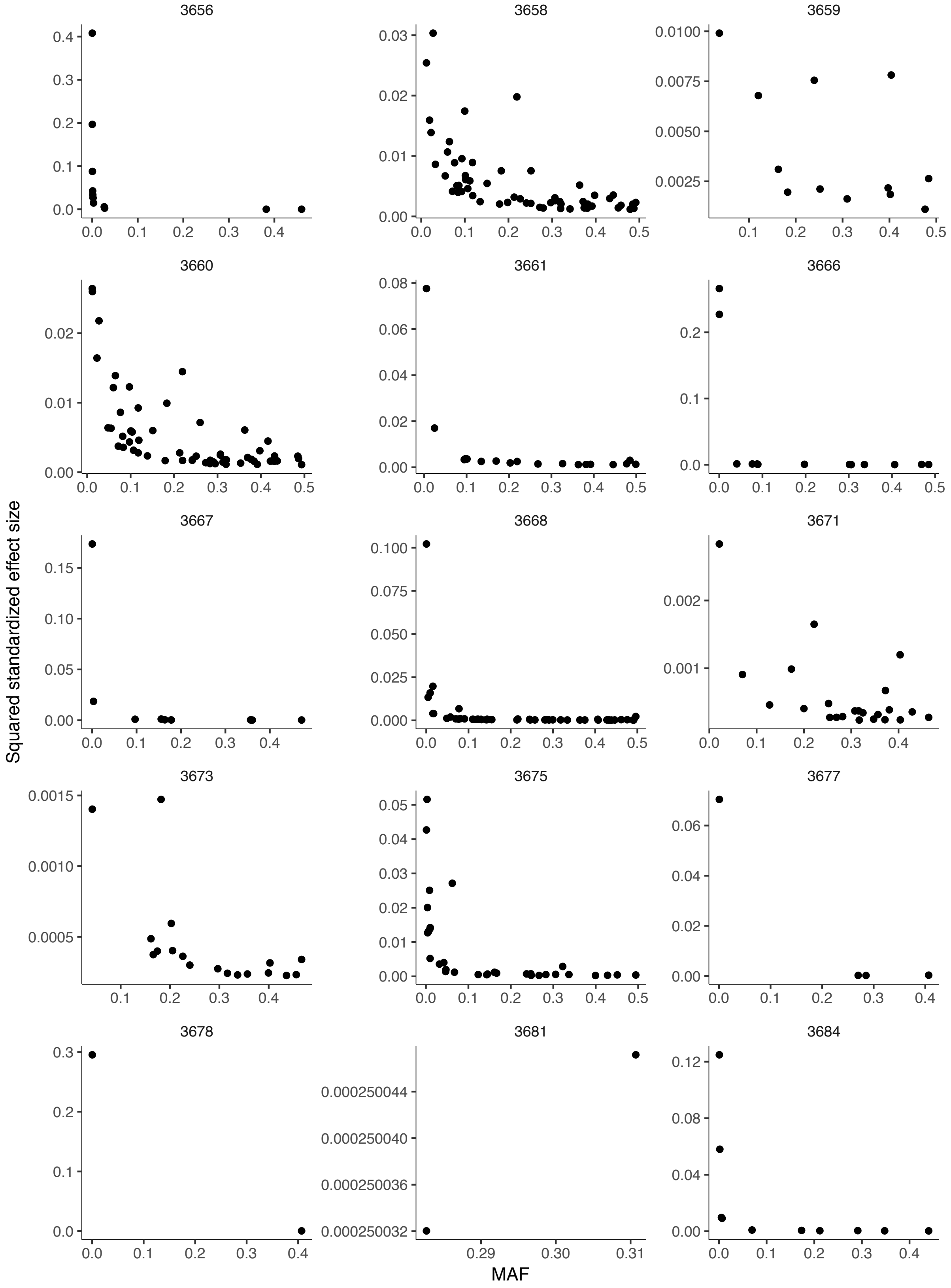
3656
3658
3659
3660
3661
3666
3667
3668
3671
3673
3675
3677
3678
3681
3684
Squared standardized effect size
MAF
0.000250044
0.000250040
0.000250036
0.000250032
0.29
0.30
0.31

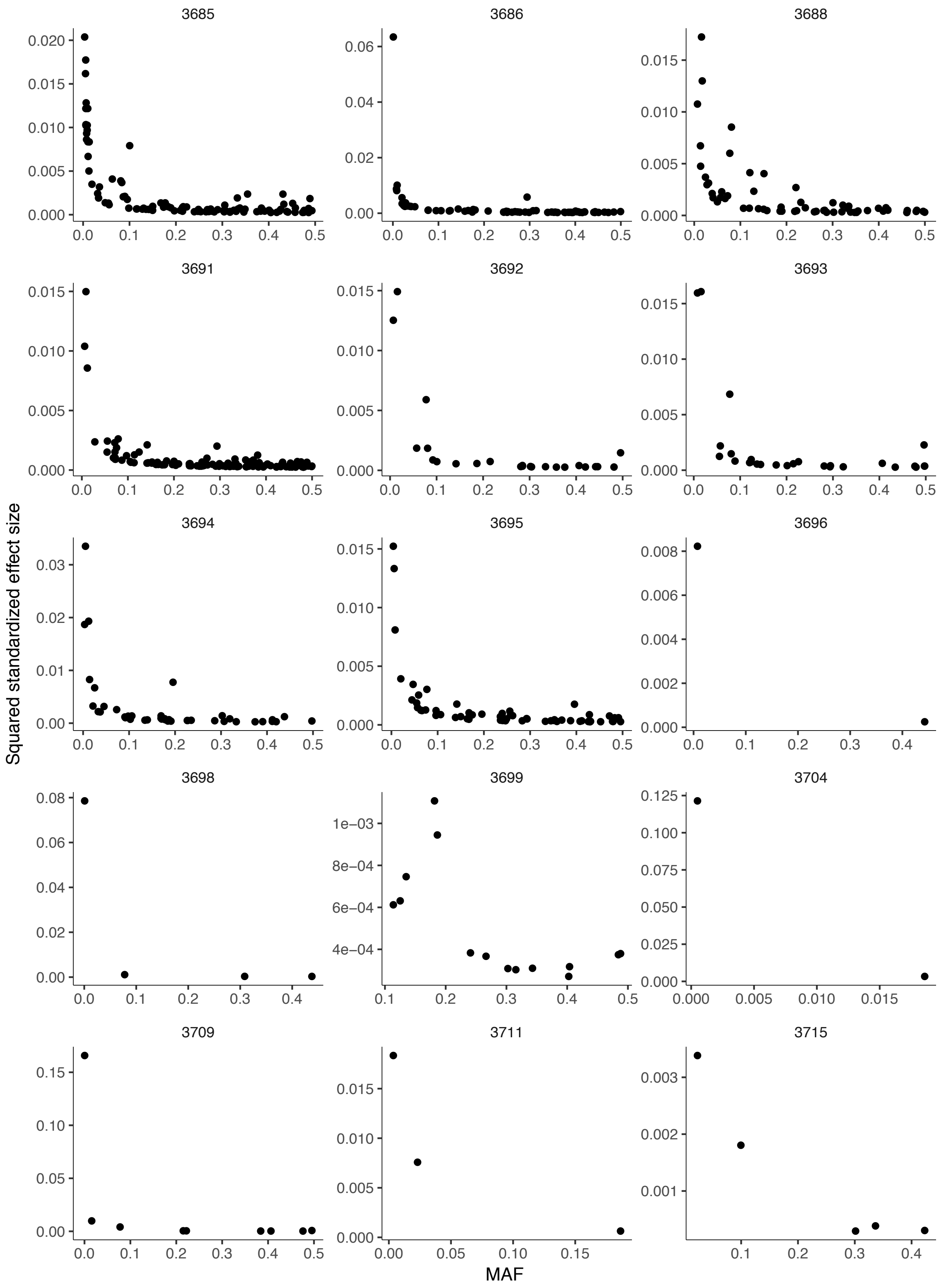
3685
3686
3688
3691
3692
3693
3694
3695
3696
3698
3699
3704
3709
3711
3715
Squared standardized effect size
MAF

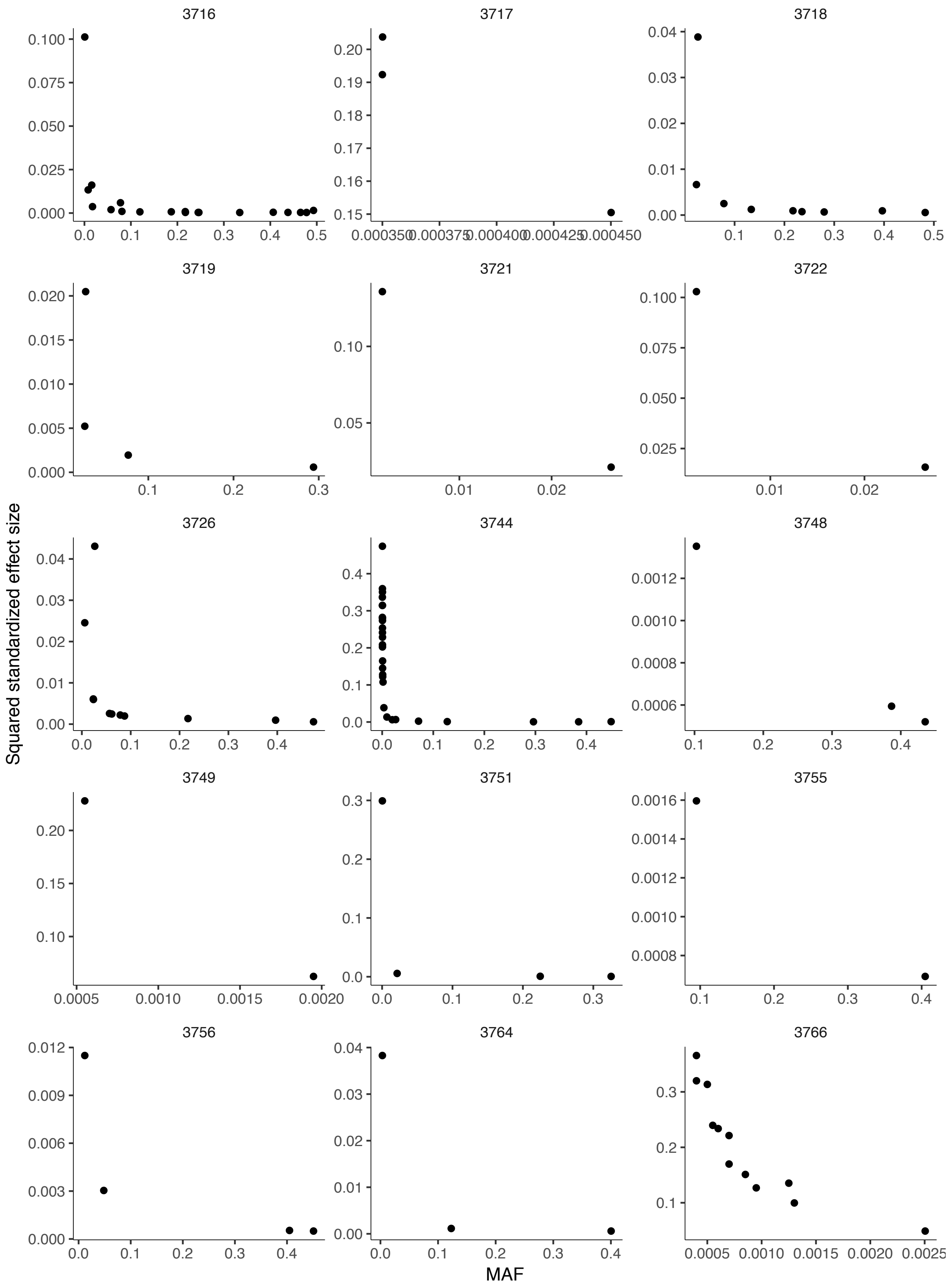
3716
3717
3718
3719
3721
3722
3726
3744
3748
3749
3751
3755
3756
3764
3766
Squared standardized effect size
MAF

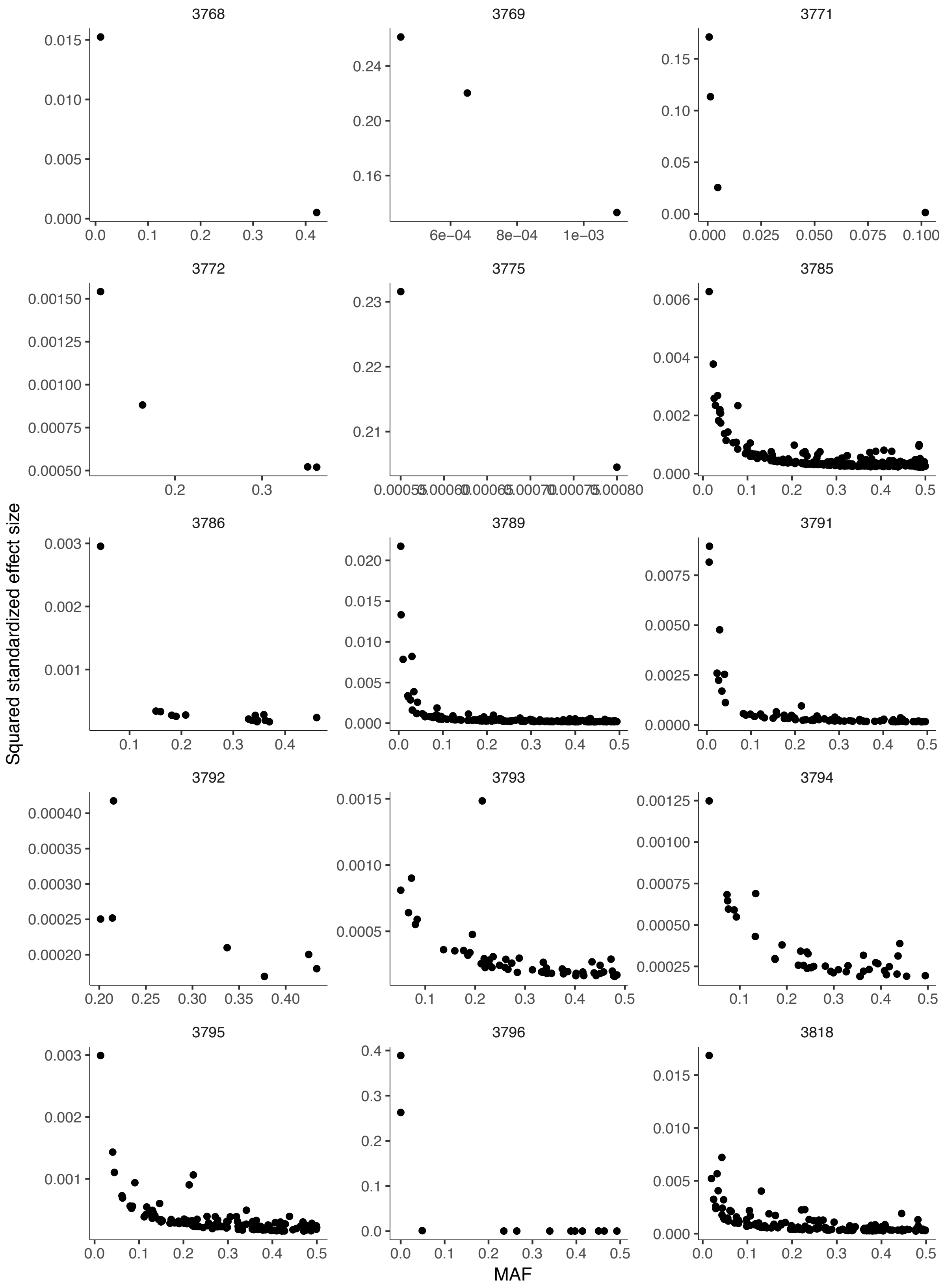
3768
3769
3771
3772
3775
3785
3786
3789
3791
3792
3793
3794
3795
3796
3818
Squared standardized effect size
MAF

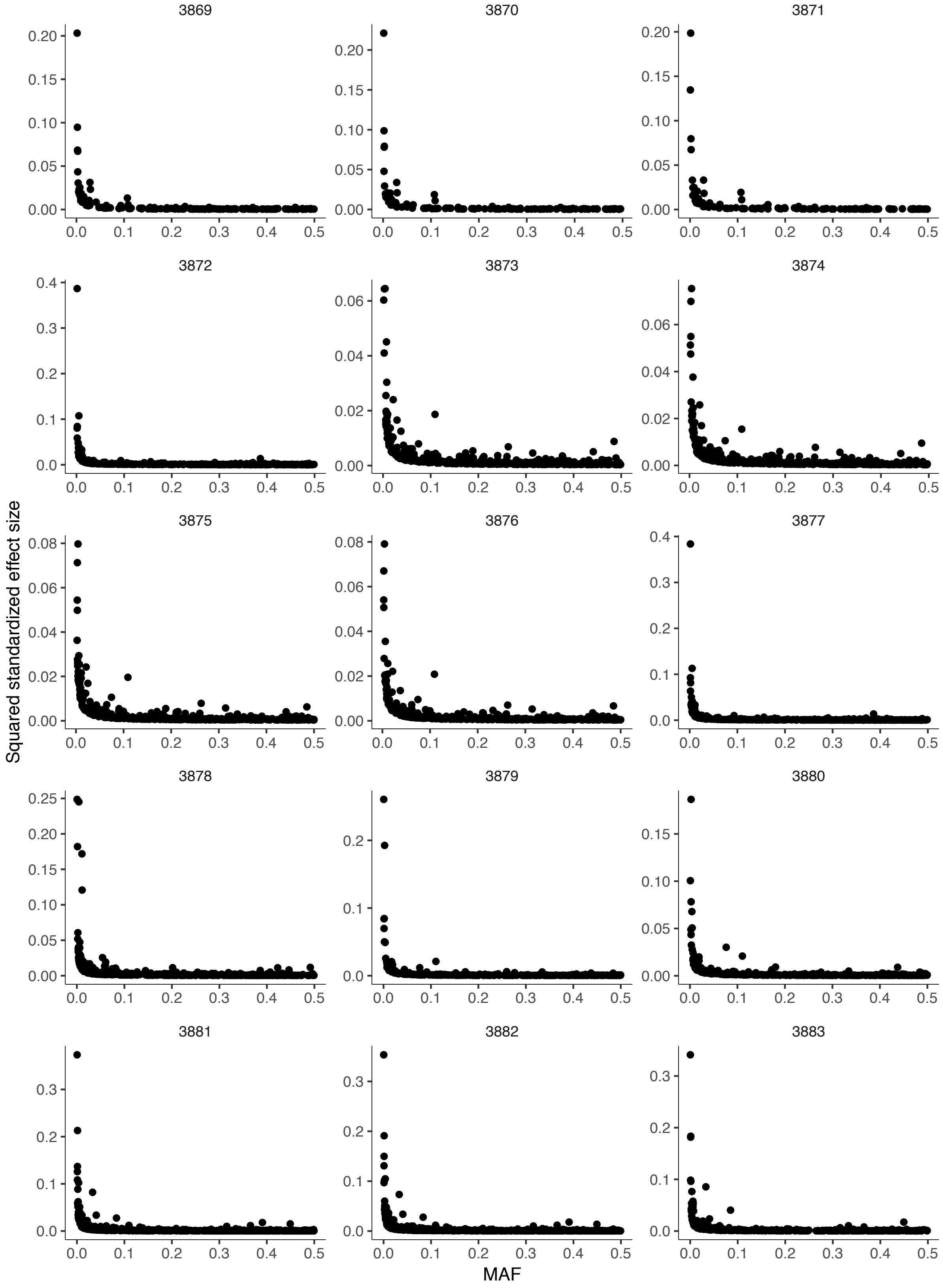
Squared standardized effect size
MAF

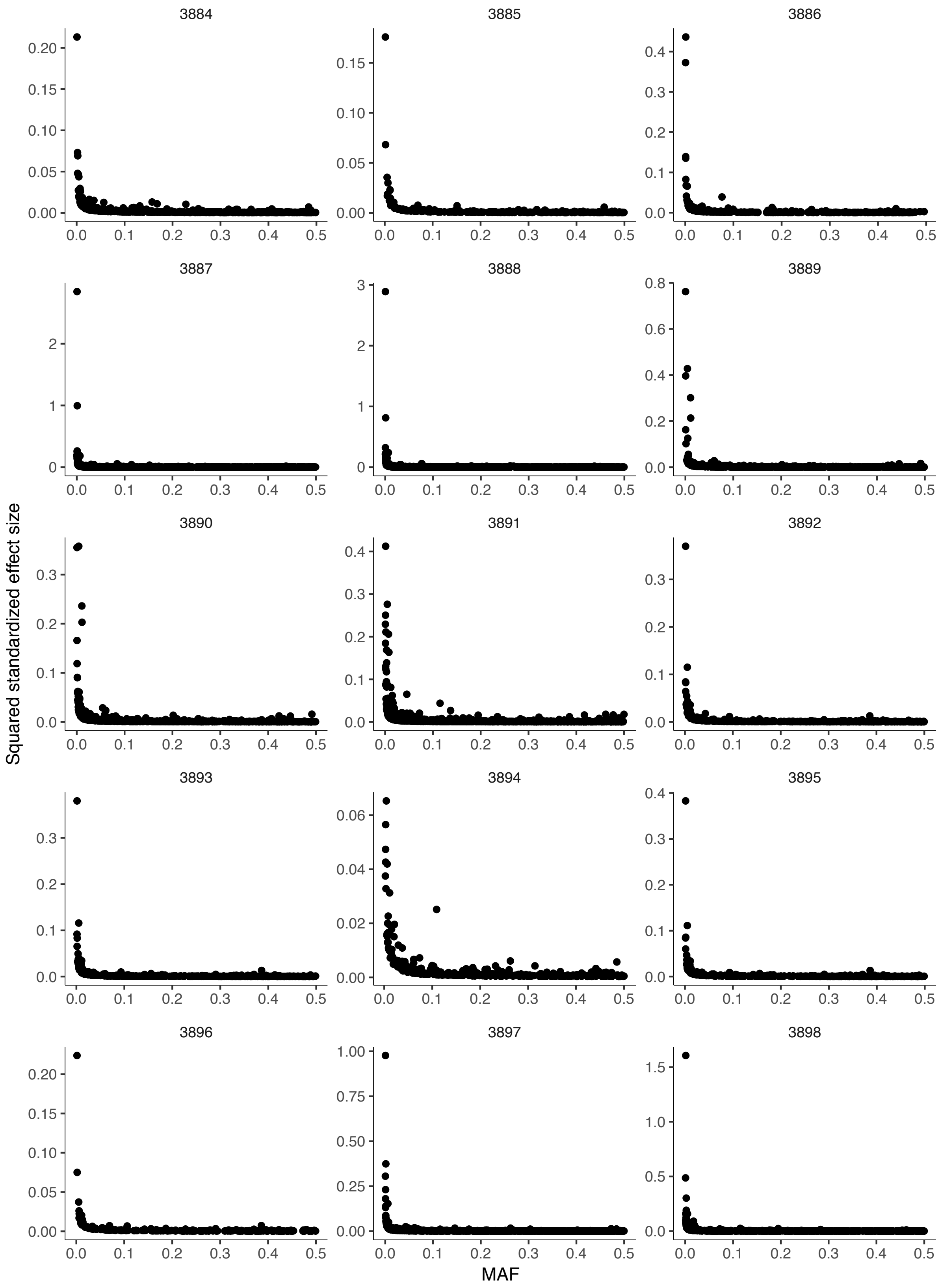
Squared standardized effect size
MAF

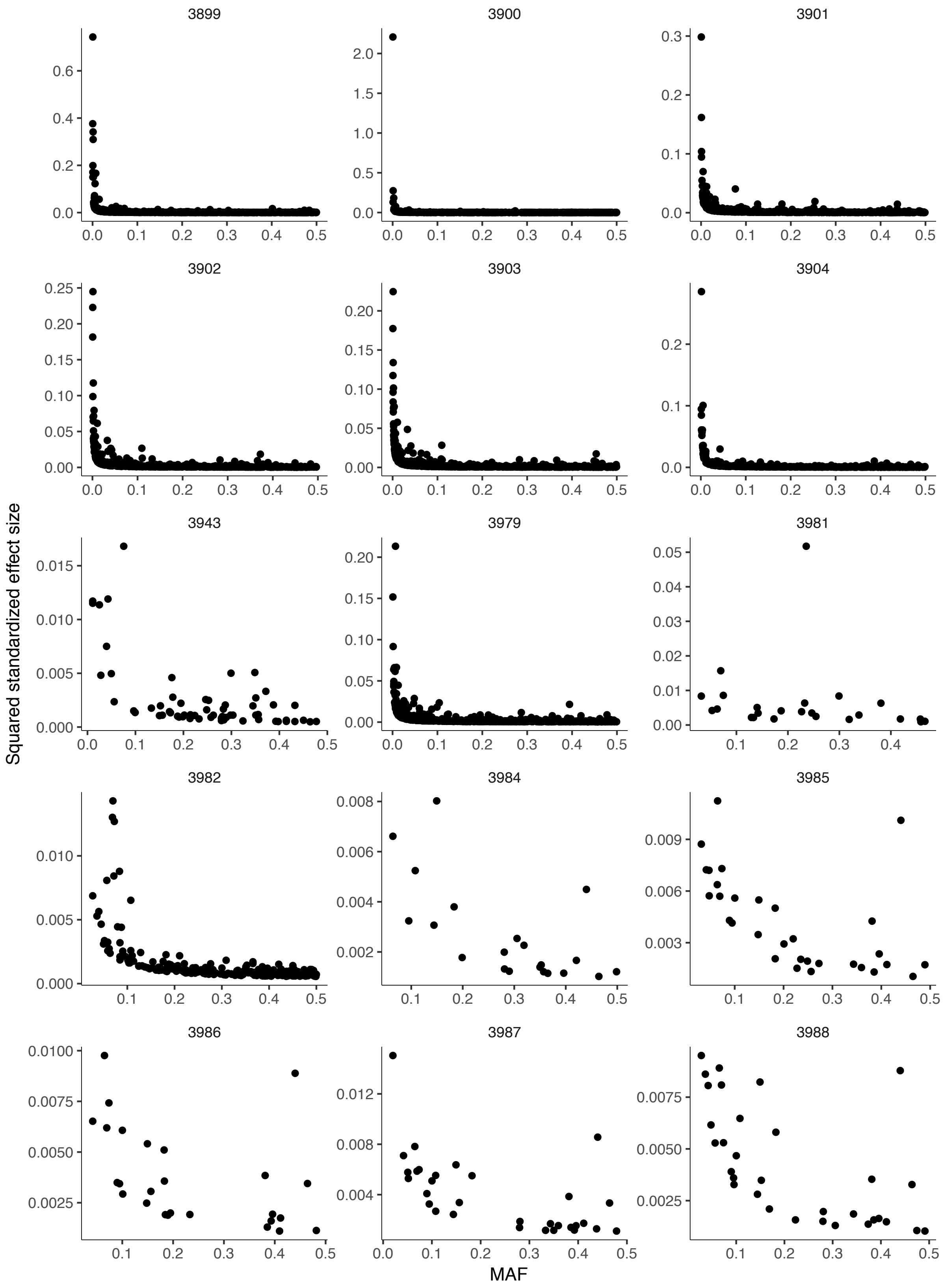

3899
3900
3901
3902
3903
3904
3943
3979
3981
3982
3984
3985
3986
3987
3988
Squared standardized effect size
MAF

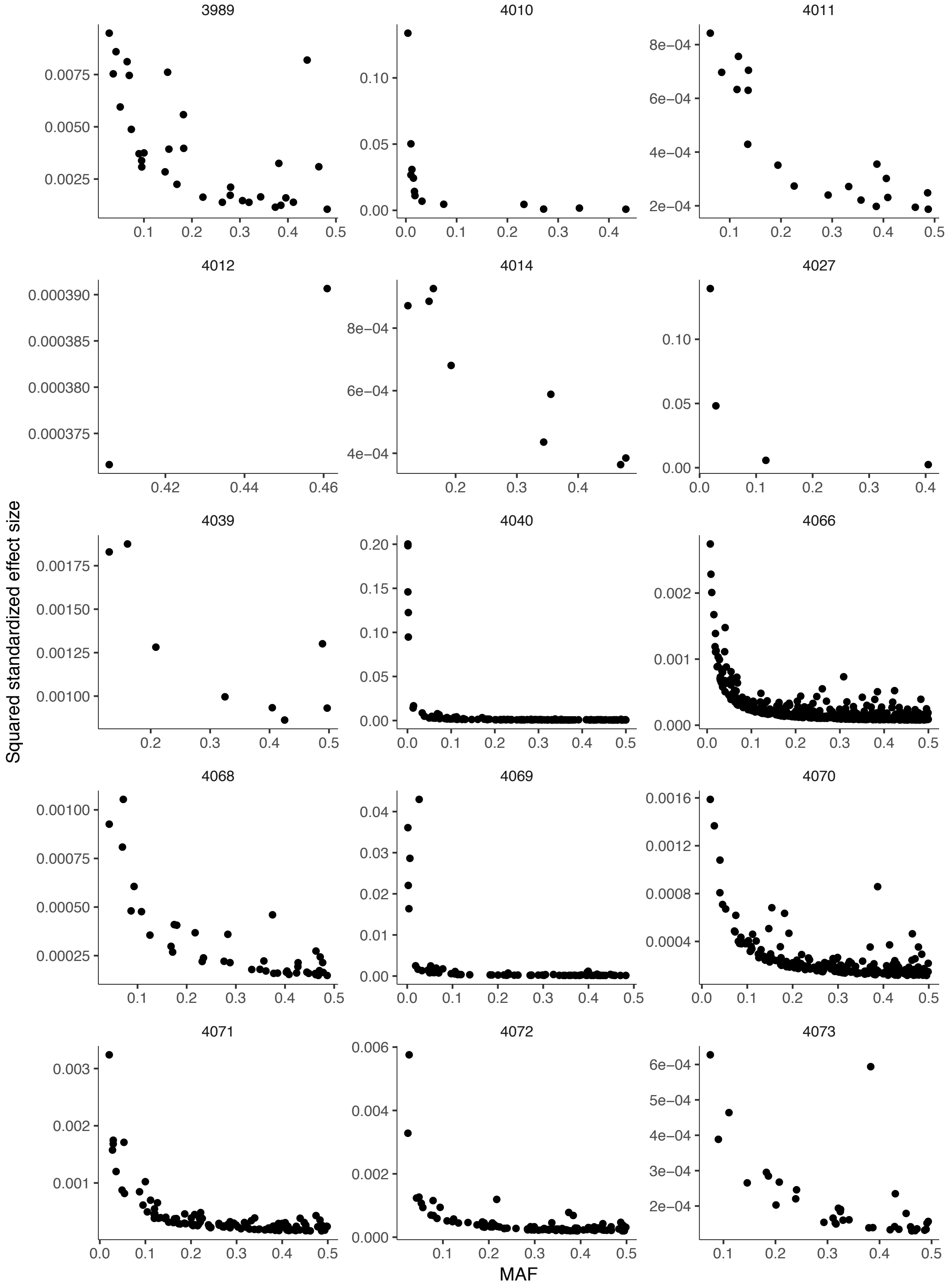

3989
4010
4011
4012
4014
4027
4039
4040
4066
4068
4069
4070
4071
4072
4073
Squared standardized effect size
MAF

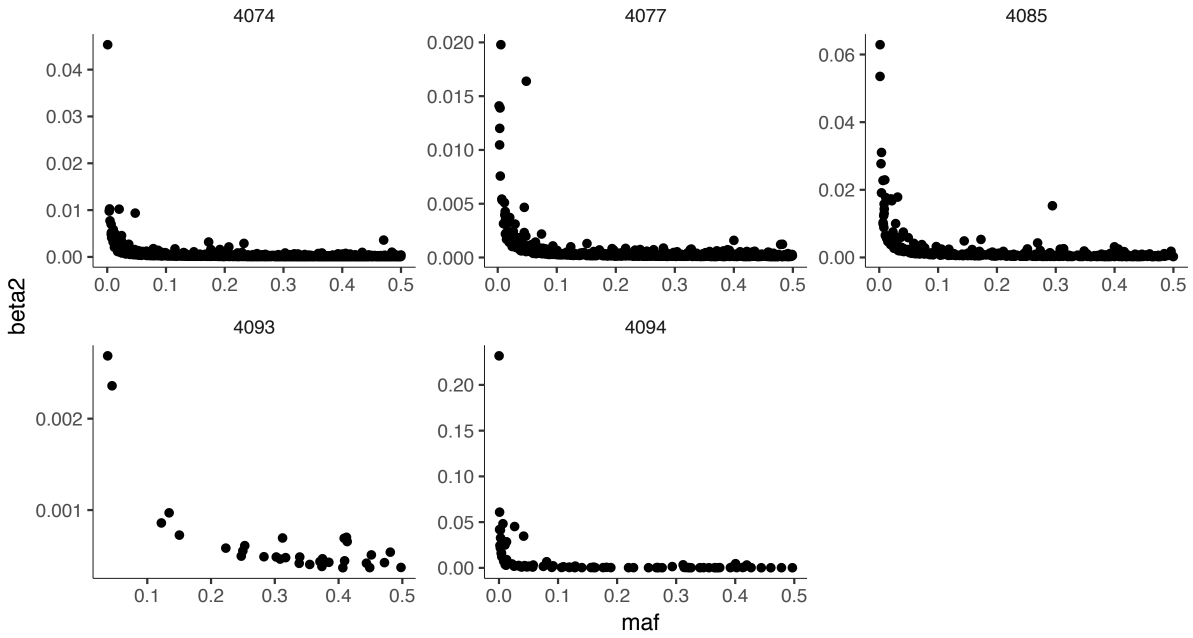

**Fig. 9. Distribution of effect size and MAF of lead SNPs for each trait.** Each of 410 traits with >1 lead SNPs are displayed. The number above the plot is the ID of the trait in the database matching with **Supplementary Table 3**.

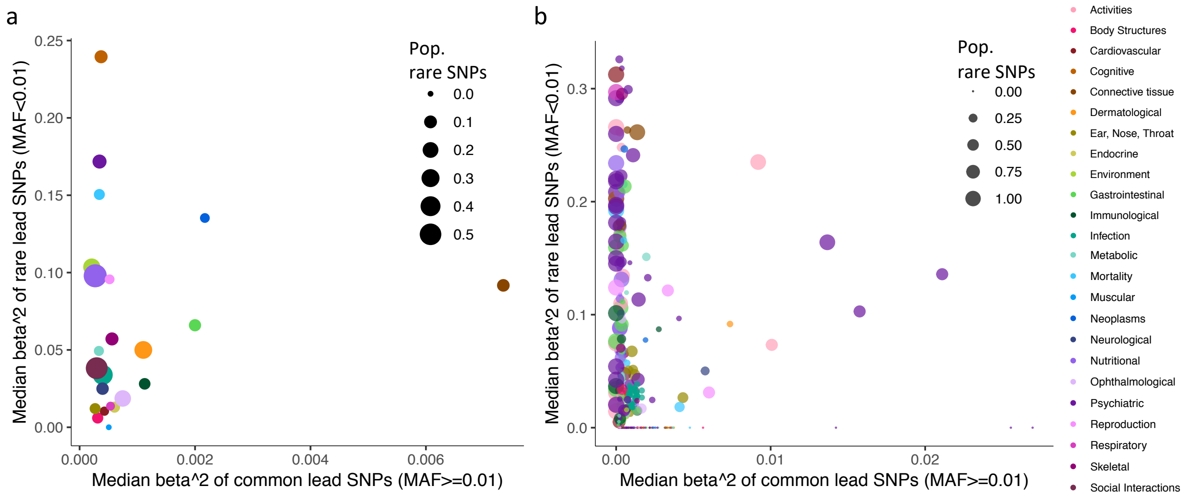

**Fig. 10. Effect size and MAF of lead SNPs. a.** Median of squared standardized effect size of unique lead SNPs per domain. **b.** Median of squared standardized effect size of unique lead SNPs per trait colored by domain.

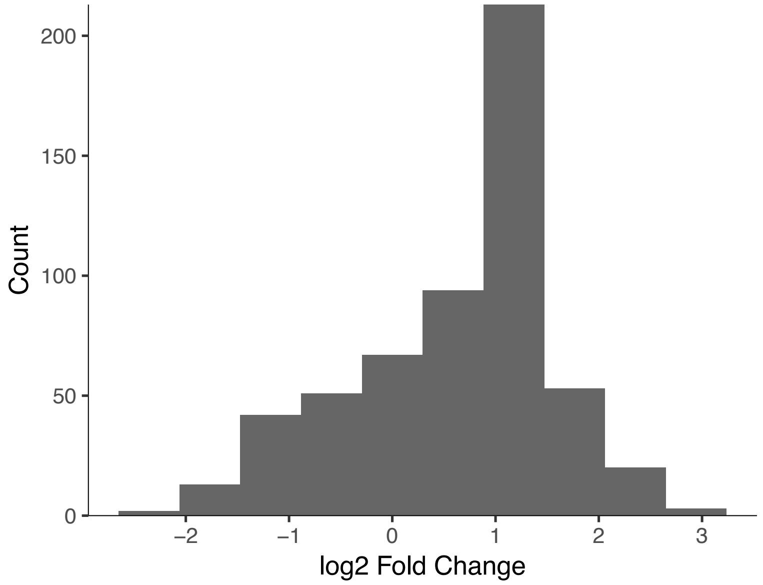

**Fig. 11. Difference of SNP heritability estimates between LDSC and SumHer.** X-axis is log 2 fold change (SumHer estimate divided by LDSC estimate).
