## Supplementary Information for "A global overview of pleiotropy and genetic architecture in complex traits"

### 1. UK Biobank release 2 dataset

#### *1.1 UK Biobank study cohort*

The current study include data from the UK Biobank study (www.ukbiobank.ac.uk)[1]. The UK Biobank is a large population-based cohort that includes over 500,000 participants and aims to improve insights into a wide variety of health-related determinants and outcomes across the UK. All participants provided written informed consent; the UK Biobank received ethical approval from the National Research Ethics Service Committee North West-Haydock (reference 11/NW/0382), and all study procedure were in accordance with the World Medical Association for medical research. Between 2006 and 2010, approximately 9.2 million invitations to participate in the study were sent to all people aged 37-72 years old who were registered with the National Health Service (NHS) and were living within 25 miles from one of the 22 study research centers. In total, 503,325 participants were recruited in the study. Access to the UK Biobank data was obtained under the application number 16406. We analyzed imputed genotype data of UK Biobank second release (July 2017)[2], which includes 92,693,895 genetic variants in 487,422 individuals. Genotypes were imputed using a combination of two reference panels; a combination of the UK10K haplotype and 1000 Genomes reference panels, and Haplotype Reference Consortium (HRC) panel[3]. If variants were imputed in both panels, the HRC imputation was retained. Details about genotyping and imputation are available in the original study[2]. We excluded variants imputed based on the combined UK10K/1000 Genomes panel according to recommendations from UK Biobank. To determine individuals of European ancestry, we projected ancestry principal components from the 1000 Genome[4] reference populations onto the called genotypes available in UK Biobank and grouped individuals into their closest ancestral population identified by the minimum Mahalanobis distance from the projected principal component scores [5]. We excluded subjects with a Mahalanobis distance >6 S.D. from their assigned population. Additional filtering of individuals was performed based on UKB-provided information on relatedness (subjects with most inferred relatives, 3rd degree or closer, were removed until no related subjects were present), discordant sex, sex aneuploidy, missing phenotype or covariate data and withdrawn consent.

Imputed variants were converted to hard calls filtering by an imputation INFO score at 0.9 and excluding multi-allelic SNPs, indels, SNPs without unique rsID and SNPs with minor allele frequency (MAF) <0.0001, resulting in a total of 10,847,151 remaining SNPs. To correct for population stratification, we computed European-specific principal components based on a set of 145,432 independent ($r^2$<0.1) SNPs with MAF>0.01 and INFO=1 using FlashPCA2[6] and first 20 PCs were used as covariates of the association analyses.

*1.2 UK Biobank phenotype definition*

Most of the phenotypes are defined as described in the method section. Quantitative and binary phenotypes with exceptional definitions are described in **Supplementary Table 1**. Multi-coded phenotypes (i.e., nominal phenotypes to which 1 (exclusive categories) or multiple (inclusive categories) answers were possible) are described in **Supplementary Table 2**.

For multi-coded phenotypes with exclusive categories (i.e., participants could select only 1 answer), we used information gathered during the first visit and first run (i.e., field codes ending with 0.0, i.e., f.xxx.0.0). These phenotypes were then dichotomized through dummy-coding (e.g., if 5 categories could be selected, 5 dummies were created in which each of the categories was contrasted against the other categories). For instance, the field f.1747.0.0 (hair color) has the following 5 categories; blonde, red, right brown, dark brown and black. In this case, the first dummy trait is blonde vs all others (red, right brown, dark brown and black) and the second is red vs all others (blonde, right brown, dark brown and black) and so on that results in 5 distinct GWAS.

For multi-coded phenotypes with inclusive categories (i.e., participants could select multiple answers, i.e., so-called multiple response questions), we used information of all runs gathered during a single visit (i.e., field codes ending with 0.x, i.e., f.xxx.0.x with exception for f.40006, f.40011 and f.40012 in which we used fields encoding f.xxx.x.0). These phenotypes were then re-coded such that every individual category was treated as a separate dichotomous question. For instance, participants could select multiple options for f.6139 (Gas or solid fuel cooking/heating; f.6139.0.0: "a gas hop or gas cooker", f.6139.0.1: "a gas fire that you used regularly in winter time", and f.6139.0.2: "an open solid fuel fire that you used regularly in winter time"). In this case, each of the three answer options was treated as a separate dichotomous question, with people endorsing the specific option coded as cases (1), and all others coded as controls (0).

The phenotypes gauging substance use (i.e., Smoking status and Alcohol Drinker status: fields f.20116.0.0 and f.20117.0.0, respectively) were originally coded as 0 (Never), 1 (Previous), and 2 (Current). For these phenotypes, we ran two separate case-control GWAS: a) Never (0) versus Ever (1+2), using the data of all participants, and b) Previous (1) versus Current (2), using only the data of people who had ever used the specific substance.

ICD10 fields often included sub-traits (e.g., ICD10 field A01 has 5 sub-traits coded A01.0 - A01.4). These sub-traits were collapsed with the main category (i.e., endorsement of any of the sub-categories A01.0 - A01.4 was treated as endorsement of the main category A01). After defining phenotypes, we recounted the total sample size used in the analyses by excluding subjects without genotype data or missing some covariates. To all available phenotypes, we applied the minimum sample size criterion of *i*) at least N=50,000 European subjects, and *ii*) in case of dichotomous phenotypes, both the number of cases and the number of controls exceeding N=10,000. All phenotypes that did not reach this criterion were excluded from further analysis.

**2. Phenotypic correlations across 457 traits**

As we expect not all traits are phenotypically independent, especially the ones in the same trait domain, using the number of associated traits to define the level of pleiotropy is not fair since the number of traits per domain varies, and trait correlations within domains may also vary. To test how dependent the traits are, we extracted phenotypes of 457 UKB2 traits out of 558 selected traits for which we have individual level phenotype information. Then computed the phenotypic correlation ($r_p$) as the Pearson's correlation of standardized phenotype. As expected, our results showed that traits within the same domain have relatively higher phenotypic correlations (average $|r_p|$=0.10) compared to traits from different domains (average $|r_p|$=0.03). We next clustered traits based on the phenotypic correlations using hierarchical clustering by optimizing the number of clusters $k$ while maximizing silhouette score. Non-significant correlations after Bonferroni correction were replaced with zero. We identified 160 clusters (**Extended Data Fig. 2a, b**). 44 traits (27.5%) did not cluster together with any other trait (i.e. clusters consisting only one trait) and 126 out of 160 clusters (78.7%) contain traits from the same domain (**Extended Data Fig. 2c**). These results show that, traits are less likely to be clustered with traits from different trait domains and generally tend to cluster within the same domain. This suggests that when assessing pleiotropy purely based on the number of associated traits, (i.e. regardless of trait domains), the observed stronger phenotypic correlations between traits within the same domain may induce pleiotropy. In addition, the 558 traits were not equally distributed across 24 trait domains. Therefore, we primarily defined the level of pleiotropy based on the number of associated domains, to avoid biasing the extent of pleiotropy due to within-domain phenotypic trait correlations. The number of associated traits was only used to distinguish trait-specific associations from potentially pleiotropic associations within domains.

**3. Colocalization of the trait-associated loci**

To distinguish pleiotropy due to LD overlap between associated loci, from likely true pleiotropy (i.e. the same causal SNP is associated with multiple trait domains) of the loci, we colocalized each trait-associated locus with all of the physically overlapping loci. We defined a pair of loci sharing a causal SNP when the posterior probability of sharing the same causal SNP was greater than 0.9. Out of 41,511 loci, 40,238 loci were physically overlapping with at least one of the other loci. 88.5% (35,603 loci) of these were colocalized with at least one locus from a different trait, and 56.5% (20,106 loci) were colocalized with trait(s) from different trait domain(s) (**Supplementary Table 6**). We then re-grouped 35,603 loci based on the colocalization pattern, so that a grouped locus is a connected component of the graph consists of loci within a grouped locus based on physical overlap and pairs of colocalized loci are linked (note that it does not require all loci within a group to be colocalized with all other loci; see Material and Methods for details). This resulted in 3,529 connected components. For each component, the number of traits and domains were counted (**Extended Data Fig. 4c** and **Supplementary Table 7**). Out of 2,089 grouped loci with more than one associated trait based on physical overlap, 1,452 loci contained more than one connected component (median 3 and average 5 independent components; **Extended Data Fig. 3**). For 1,625 multi-domain loci (defined by the physical overlap), the number of associated trait domains of the largest connected component per grouped locus showed on average 38% decrease compared to the number of domains in the grouped loci based on physical overlap (**Extended Data Fig. 4d**). In addition, we computed the proportion of colocalized pairs for each connected component. When the proportion is 1, the connected component is a clique (fully connected graph, i.e. every locus is colocalized with every other). We found that the proportion of colocalized pairs tends to decrease along with the number of trait domains in the components (**Extended Data Fig. 4e**). This suggests that larger connected components likely include “hub” loci that are colocalized with a lot of loci from different traits and domains. However, 67 grouped loci contained associations with ≥5 domains and had a colocalized proportion >0.5, indicating high pleiotropy of these loci.

**4. Gene length and LD for MAGMA gene analysis**

We showed that more pleiotropic genes are significantly longer compared to genes that are not associated with any of the 558 traits. This is unlikely to be due to artifacts, such as the fact that longer genes containing more SNPs and therefore would have a higher chance of including significant associations, since MAGMA controls for this by projecting a SNP matrix on principal components (PC) and pruning away PCs with small eigenvalues[7]. Therefore, the number of parameters per gene is independent on the length of genes. Indeed the number of parameters done not significantly differ between multi-domain, domain specific, trait specific and non-associated genes, except a slight increase between non-associated genes to multi-domain genes (**Extended Data Fig. 5c** and **Supplementary Table 10**).

**5. Distribution of pleiotropic SNPs across chromosomes**

To test if the pleiotropic SNPs are evenly distributed across chromosomes, we first pruned the 1,740,179 SNPs and extracted 822,052 independent SNPs at $r2<0.1$ with maximum distance 1Mb. We then performed the following 4 types of tests.

*i) Enrichment of significant SNPs in each chromosome*

For each chromosome, Fisher's exact test (two-sided) was performed for the number of significant SNPs in a certain chromosome against all chromosomes. When the proportion of a chromosome is higher than the total proportion, that means positive enrichment, otherwise negative.

*ii) Enrichment of pleiotropic SNPs in each chromosome*

For each chromosome, Fisher's exact test (two-sided) was performed for the number of pleiotropic SNPs associated with more than one trait out of SNPs associated with at least one trait in a certain chromosome against all chromosomes. Again, when the proportion of a chromosome is higher than the total proportion, that means positive enrichment, otherwise negative.

*iii) Enrichment of highly pleiotropic SNPs in each chromosome*

For each chromosome, Fisher's exact test (two-sided) was performed for the number of highly pleiotropic SNPs associated with more than one domain out of SNPs associated with at least one trait in a certain chromosome against all chromosomes. When the proportion of a chromosome is higher than the total proportion, that means positive enrichment, otherwise negative.

*iv) Deviation of the level of pleiotropy of SNPs across chromosomes*

For each chromosome, a Wilcoxon rank sum test (one-sided) was performed for the number of associated domains of SNPs associated with at least one trait in a certain chromosome compared to other chromosomes. A statistically significant test result suggests a significantly higher level of pleiotropy of SNPs (in terms of the number of associated domains) in the tested chromosome compared to all others.

**6. Pleiotropic gene-sets**

The level of pleiotropy of the gene-sets may depends on the definition of tested gene-sets, e.g. if only limited number of genes were included in the current gene-sets, this might bias the results. Therefore, we evaluated distribution of pleiotropic genes across gene-sets. Out of 17,444 genes analyzed in this study (that were tested in all 558 traits), 16,401 genes (94.0%) were assigned to one of the 10,650 tested gene-sets. Genes that are not included in any of the gene-sets showed significantly lower pleiotropy compared to genes involved in any of the gene-sets ($p$=2.2e-15, two-sided Mann-Whitney U test on the number of associated domains of genes). Therefore, this cannot explain that higher proportion of trait-specific gene-sets are due to exclusion of highly pleiotropic genes from the tested gene-sets.

**7. Generic correlations across 558 traits**

Although the proportion of trait pairs with a significant genetic correlation were generally higher within domain than between domains, it is important to note that the trait domains are assembled based on external definitions which might partially reflect phenotypic correlations, but not genetic similarity. To identify genetically similar clusters of traits, we performed hierarchical clustering on the $r_g$ matrix and identified 62 clusters by optimizing the number of clusters $k$ with maximizing silhouette score (non-significant $r_g$*'s* were replaced by zero, see **Methods** for details). One large cluster contained 258 traits (46.2% of 558 traits) that are clustered together mainly due to low $r_g$ (or non-significance due to low power) across traits (**Extended Data Fig. 8**). Apart from that cluster, 18 traits were not clustered with any other traits and 7 clusters contained traits from the same domain, while the remaining 36 clusters contain traits from multiple domains (**Extended Data Fig. 8b**).

**8. Distribution of MAF and effect sizes of lead SNPs**

To test whether proportion of rare lead SNPs (MAF<0.01) is higher or lower than expected, we compared with the number of SNPs with MAF<0.01 in UKB2 reference panel, as >85%

of traits are based on UKB2. In GWASs performed in this study, 10,846,943 SNPs were tested, of which 33.6% of SNPs had MAF<0.01. The proportion of rare lead SNPs (12.3%) is, therefore, less than expected ($p$<1e-323, two-sided Fisher's exact test). We note, however, that the lower proportion of rare lead SNPs may be biased by the different QC criteria across different studies, especially for non-UKB2 GWASs (i.e. it is possible that rare SNPs are excluded prior to GWASs).

Rare lead SNPs (MAF<0.01) showed the highest median of squared effect size ($\beta^2$) in the cognitive domain ($\beta^2$=0.24) followed by psychiatric ($\beta^2$=0.17) and mortality ($\beta^2$=0.15) domains (**Supplementary Table 20**). For common lead SNPS, most of the domains showed a median of $\beta^2$<0.001 except connective tissue (7.4e-3), neoplasms (2.2e-3) and dermatological (1.1e-3) domains. At a trait level, we again observed a wide variety in effect sizes, with the high median $\beta^2$ (>0.01) of common SNPs (MAF>=0.01) for lead SNPs of several psychiatric traits, such as 'Felt distant from other people in past month' and 'frequency of memory loss due to drinking alcohol in last year' (**Supplementary Table 20**). Genetic associations for the nutritional, environment and social interactions domains have a higher proportion of rare lead SNPs while the effect sizes of these are moderate compared to other domains (**Extended Data Fig. 10a**). On the other hand, the cognitive, psychiatric and neoplasms domains showed much smaller proportions of rare variants in genetic associations, yet with higher effect size. Within domains, traits also differed with respect to the frequency and effect sized of associated lead SNPs. For example, 'Frequency of tenseness / restlessness in last 2 weeks' has a very small proportion of rare lead SNPs with high effect size, while 'Frequency of memory loss due to drinking alcohol in last year' has relatively higher proportion of rare lead SNPs with low effect size but higher effect size for common lead SNPs (**Extended Data Fig. 10b** and **Supplementary Table 21**).

**9. Fine-mapping of the trait-associated loci**

Fine-mapping was performed for a region of 50kb centered around the top SNP of the locus (unless the locus was larger than that window). Loci overlapping with the MHC region (chr6:25Mb-36Mb) and loci with less than 10 SNPs were excluded. In total, 40,875 loci from 459 traits containing 76,162 lead SNPs were fine-mapped. We then extracted 95% credible sets by taking configs (sets of $k$ causal SNPs) until the cumulative sum of posterior probably reaches 0.9 (see Materials and Methods for details). The credible sets consist of 1,110,733 SNPs of which 563,382 are unique. 53% (21,690) of loci contained the top SNPs in credible sets and 32% (24,678) of lead SNPs were included in credible sets. In the 89.9% (36,771) of loci, the number of causal SNPs was optimized at $k$=10, which also suggests that there might be more than 10 as it was restricted to the maximum of 10, while only 1,191 loci were optimized at $k$=1. Each SNP in 95% credible sets also has posterior inclusion probability (PIP) which measures how likely the SNP is to be included in the causal configs. In total 27.0% (196,542) of SNPs showed PIP>0.95 from 38,356 loci which suggests that a large proportion of SNPs in credible sets are unsolved mainly due to strong LD. In addition, false positive rate is much higher when a reference panel is used rather than the matched genotype data for fine-mapping [8]. We therefore focused on SNPs with PIP>0.95 which are referred as credible SNPs.

**10. SNP heritability estimate based on LD score regression and SumHer**

$h^2_{SNp}$ is different from the broad sense heritability ($H^2$) which is generally based on twin or family studies and inferred from known genetic relationships. When $h^2_{SNP}$ is lower than $H^2$, this 'still-missing heritability' may indicate that non-additive effects are responsible for some of the $H^2$ or that variants not included or captured in the GWAS such as rare variants or structural variants make a substantial contribution. In addition, phenotypic heterogeneity of the trait might also explain the low $h^2_{SNP}$ compared to $H^2$[29]. Although $h^2_{SNP}$ itself is not directly informative of the genetic architecture of a trait (in terms of the number of associated SNPs, allele frequencies or the effect sizes), it does provide insights into the relative contribution of additive SNPs effects across traits.

LDSC was introduced in 2015 and has been widely used since then to estimate $h^2_{SNP}$ from summary statistics. LDSC assumes that the contribution of each SNP to the total $h^2_{SNP}$ is constant[10]. Recently, the LDAK model has been introduced as an alternative, relying on the assumption that the contribution of each SNP to the total $h^2_{SNP}$ is a function of its MAF, and SNPs are weighted by their local LD[11]. SumHer implements the LDAK model, and like LDSC, it can be used to estimate $h^2_{SNP}$ from summary statistics[12]. The models underlying LDSC and SumHer thus rely on different assumptions, and therefore estimated $h^2_{SNP}$ values may differ between LDSC and SumHer. To facilitate a fair comparison between the two methods, we used the 1000 Genome Phase 3 European reference panel[4] and limited our analyses to HapMap3 SNPs (see **Methods**).

Overall, 74.7% (417/558) of traits showed higher estimates in SumHer than in LDSC (**Fig. 4a** and **Supplementary Table 22**), of which 268 traits showed more than a 2-fold increase (**Extended Data Fig. 11**). As Speed *et al.* reported in their studies based on simulations,

LDSC tends to under-estimate the $h^2_{SNP}$ when a phenotype follows the LDAK model but at the same time SumHer tends to overestimate when a trait follows the LDSC model[11,12]. Notably, SumHer estimated $h^2_{SNP}>1$ for 8 metabolic traits which also had very high standard error (average 0.33). In addition, as presented by Luke et al.[13], LDAK is highly sensitive to misspecification of the model assumptions; if causal variants of a trait are mainly common, weighting SNPs by the inverse proportion of MAF can highly bias the $h^2_{SNP}$ estimate[13]. Since the true underlying genetic model (whether the contribution of each SNP is constant or a function of MAF, or even a mixture of these two) of a trait is unknown, the true value of $h^2_{SNP}$ could be much higher than the LDSC estimates (which can be considered as a lower bound), but lower than the SumHer estimates.

**References**


1. Sudlow, C. *et al.* UK Biobank: An open access resource for identifying the causes of a wide range of complex diseases of middle and old age. *PLoS Med.* **12,** 1–10 (2015).
2. Bycroft, C. *et al.* The UK Biobank resource with deep phenotyping and genomic data. *Nature* **562,** 203–209 (2018).
3. Mccarthy, S. *et al.* A reference panel of 64,976 haplotypes for genotype imputation. *Nat. Genet.* **48,** 1279–1283 (2016).
4. Auton, A. *et al.* A global reference for human genetic variation. *Nature* **526,** 68–74 (2015).
5. Webb, B. T. *et al.* Molecular Genetic Influences on Normative and Problematic Alcohol Use in a Population-Based Sample of College Students. *Front. Genet.* **8,** 30 (2017).
6. Abraham, G., Qiu, Y. & Inouye, M. FlashPCA2 : principal component analysis of Biobank-scale genotype datasets. *Bioinformatics* **33,** 2776–2778 (2017).
7. de Leeuw, C. A., Mooij, J. M., Heskes, T. & Posthuma, D. MAGMA: generalized gene-set analysis of GWAS data. *PLoS Comput. Biol.* **11,** e1004219 (2015).
8. Benner, C. *et al.* Prospects of fine-mapping trait-associated genomic regions by using summary statistics from genome-wide association studies. *Am. J. Hum. Genet.* **101,** 539–551 (2017).
9. Wray, N. R. & Maier, R. Genetic Basis of Complex Genetic Disease: The Contribution of Disease Heterogeneity to Missing Heritability. *Curr. Epidemiol. Reports* **1,** 220–227 (2014).
10. Bulik-sullivan, B. K. *et al.* LD Score regression distinguishes confounding from polygenicity in genome-wide association studies. *Nat. Genet.* **47,** 291–295 (2015).
11. Speed, D. *et al.* Reevaluation of SNP heritability in complex human traits. *Nat. Genet.* **49,** 986–992 (2017).
12. Speed, D. & Balding, D. J. Better estimation of SNP heritability from summary statistics provides a new understanding of the genetic architecture of complex traits. *Nat. Genet.* (2018).
13. Evans, L. M. *et al.* Comparison of methods that use whole genome data to estimate the heritability and genetic architecture of complex traits. *Nat. Genet.* **50,** 737–745 (2018).

**Supplementary Tables**

**Table 1. Exceptional phenotype definition for UK Biobank traits.**

**Table 2. Definition of multi-coded phenotypes for UK Biobank traits.**

**Table 3. GWAS summary statistics in the atlas database.**

**Table 4. Physically overlapping trait-associated loci groups.**

**Table 5. Wilcoxon rank sum test of gene density for grouped loci across 4 categories of loci associations.**

**Table 6. Trait-associated loci grouped based on colocalization pattern (connected components of colocalized loci).**

**Table 7. Genes significantly associated with at least one trait.**

**Table 8. T-test of the gene length (in log scale) across 4 categories of gene associations.**

**Table 9. T-test of the number of parameters for MAGMA gene analysis (in log scale) across 4 categories of gene associations.**

**Table 10. Wilcoxon rank sum test of pLI score of genes across 4 categories of gene associations.**

**Table 11. Level of gene pleiotropy and tissue specificity of gene expression.**

**Table 12. SNPs significantly associated with at least one traits.**

**Table 13. Distribution of pleiotropic SNPs across chromosomes.**

**Table 14. Level of SNP pleiotropy and functional consequences.**

**Table 15. Level of SNP pleiotropy and tissue specificity based on eQTLs.**

**Table 16. Gene sets significantly associated with at least one traits.**

**Table 17. T-test of the number of genes in gene sets (in log scale) across 4 categories of gene set associations.**

**Table 18. Within domain density of genetic correlation.**

**Table 19. Between domain density of genetic correlations.**

**Table 20. Median beta^2 of lead SNPs per domain.**

**Table 21. Median beta^2 of lead SNPs per trait.**

**Table 22. SNP heritability estimates by LDSC and SumHer.**

**Table 23. Polygenicity and discoverability estimates by UGMG.**

**Table 24. Reasons of exclusion from database for dbGAP GWAS entry.**

**Table 25. Curated population prevalence of binary traits.**
